# Loss of multi-level 3D genome organization during breast cancer progression

**DOI:** 10.1101/2023.11.26.568711

**Authors:** Roberto Rossini, Mohammadsaleh Oshaghi, Maxim Nekrasov, Aurélie Bellanger, Renae Domaschenz, Yasmin Dijkwel, Mohamed Abdelhalim, Philippe Collas, David Tremethick, Jonas Paulsen

**Affiliations:** Department of Biosciences, Faculty of Mathematics and Natural Sciences, University of Oslo, 0316 Oslo, Norway; Department of Genome Sciences, The John Curtin School of Medical Research, The Australian National University, Canberra, Australian Capital Territory, Australia; Department of Molecular Medicine, Institute of Basic Medical Sciences, Faculty of Medicine, University of Oslo, 0317 Oslo, Norway; Department of Immunology and Transfusion Medicine, Oslo University Hospital, 0424 Oslo, Norway; Centre for Bioinformatics, Department of Informatics, University of Oslo, 0316 Oslo, Norway

**Author notes:** Correspondence (PC); (DT); (JP).

## Abstract

Breast cancer entails intricate alterations in genome organization and expression. However, how three-dimensional (3D) chromatin structure changes in the progression from a normal to a breast cancer malignant state remains unknown. To address this, we conducted an analysis combining Hi-C data with lamina-associated domains (LADs), epigenomic marks, and gene expression in an *in vitro* model of breast cancer progression. Our results reveal that while the fundamental properties of topologically associating domains (TADs) are overall maintained, significant changes occur in the organization of compartments and subcompartments. These changes are closely correlated with alterations in the expression of oncogenic genes. We also observe a restructuring of TAD-TAD interactions, coinciding with a loss of spatial compartmentalization and radial positioning of the 3D genome. Notably, we identify a previously unrecognized interchromosomal insertion event, wherein a locus on chromosome 8 housing the *MYC* oncogene is inserted into a highly active subcompartment on chromosome 10. This insertion is accompanied by the formation of *de novo* enhancer contacts and activation of *MYC*, illustrating how structural genomic variants can alter the 3D genome to drive oncogenic states. In summary, our findings provide evidence for the loss of genome organization at multiple scales during breast cancer progression revealing novel relationships between genome 3D structure and oncogenic processes.

## Introduction

The mammalian genome is folded into large-scale dynamic 3-dimensional (3D) chromatin conformations that provide a framework for regulated gene expression. In the interphase nucleus, chromosomes exist as territories (Rowley and Corces 2018; Cremer and Cremer 2010) within which gene-rich and active A-compartments are segregated from more gene-poor and inactive B compartments (Misteli 2004; Crosetto and Bienko 2020; Lieberman-Aiden et al. 2009), both ranging in size from single promoters up to 2 megabase-pairs (Mbp) (Sood and Misteli 2022; Harris et al. 2023). Compartments consist of multiple sub-compartments that differ in their chromosomal contact frequencies, enrichment in histone modifications and chromatin-binding proteins (Spracklin et al. 2023). At a resolution ranging from ∼10-800 kilobase-pairs (kbp), topologically associated domains (TADs) emerge as local, highly interacting domains (Rao et al. 2014; Nora et al. 2012; Dixon et al. 2012). TADs are further organized into sub-TADs and support the formation of internal smaller chromatin loops, which includes enhancer-promoter interactions (Berlivet et al. 2013). TADs and sub-TADs seemingly form independent of compartments (Rowley and Corces 2018). It was initially proposed that TADs provide a platform for the coordinated regulation of subsets of genes by enhancers located within the boundaries of the TAD. However, disrupting TAD structure minimally affects gene expression (Nora et al. 2017; Rao et al. 2017; Schwarzer et al. 2017), suggesting that other levels of genome organization have a more important regulatory role.

An intriguing possibility is that long-range TAD-TAD interactions, forming TAD cliques (Zhao et al. 2023; Pei et al. 2022; Li et al. 2022; Arnould et al. 2023; Paulsen et al. 2019), sculpt the 3D genome into distinct functional domains. TAD cliques are enriched in B compartments, and can gain or lose TADs during differentiation, generally in correlation with gene repression or activation, respectively (Paulsen et al. 2019). Additionally, the nuclear envelope imposes positional constraints on chromatin by anchoring heterochromatin through lamina-associated domains (LADs) of ∼0.1-10 Mbp in size (Gonzalez-Sandoval and Gasser 2016; Rønningen et al. 2015; van Steensel and Belmont 2017). Accordingly, genes located at the nuclear periphery are repressed or expressed at lower levels than genes localized towards the nuclear center (Misteli 2004; Crosetto and Bienko 2020). Interestingly, integrating Hi-C and LAD data allows the generation of 3D structural genome models that accurately recapitulate the radial nuclear positions of TADs (Paulsen et al. 2017, 2018; Li et al. 2017; Boninsegna et al. 2022), and provide an enhanced understanding of the link between chromatin architecture and gene regulation in disease contexts (Paulsen et al. 2017).

Indeed, diseases including cancer have attributed transcriptional dysregulation to alterations in 3D genome organization (Feng and Pauklin 2020; Osman et al. 2022). Yet, analysis of multiple cancers shows that TAD boundary deletions only rarely change gene expression, emphasizing their rare involvement in gene expression dysregulation in cancer (Akdemir et al. 2023). Even with shifts in A and B compartments between colon tumors and normal intestinal cells, TADs remain relatively unchanged (Johnstone et al. 2020), suggesting compartment switching significantly impacts gene expression control (Sood and Misteli 2022; Ibrahim and Mundlos 2020).

Breast cancer is the most common cancer in women (Bobbitt et al. 2023). These cancers have been classified into five subtypes based on the expression of receptors for human epidermal growth factor, estrogen and progestogen, and clinical features (Bobbitt et al. 2023). Among these subtypes, triple-negative breast cancers, which do not express any of these receptors, are the most aggressive (Bobbitt et al. 2023). Few studies have examined the chromatin architectural features of breast cancer cells or tissues (Dozmorov et al. 2023; Kim et al. 2022; Wang et al. 2022; Zhou et al. 2019). Nonetheless, one study reports that triple-negative breast cancer cells display the most severe disruption of the 3D genome, including weakening of TAD borders, loss of 3D chromatin interactions leading to fewer chromatin loops, and dynamic compartmental changes (Kim et al. 2022). However, how 3D chromatin organization is associated with the progression from a normal to a breast cancer malignant state remains unexplored. Here, we rely on an *in vitro* isogenic breast cancer progression model (Dawson et al. 1996; Santner et al. 2001) to investigate 3D genome organization changes in this process.

## Results

### TAD properties are conserved across breast cancer stages

We performed duplicate Hi-C experiments at three stages of an *in vitro* human breast cancer progression model, corresponding to a nonmalignant state (using MCF10A cells; “10A” from here on), a premalignant state (MCF10AT1 cells; “T1”) and a malignant tumorigenic state (MCF10Ca1a cells; “C1”). We obtained an average of >560 million pairwise interactions after filtering (Supplementary Table S1), supporting analyses in the range of 1-10 kbp bin resolution (Lajoie et al. 2015; Ay and Noble 2015). Filtering statistics and the relative fraction of intra- (cis) and inter- (trans) chromosomal contacts indicate high quality libraries (Lajoie et al. 2015), and samples show high reproducibility between replicates (Supplementary Fig. S1 and S2) (Yang et al. 2017). We identified and masked out regions with translocations and used ICE-normalized interactions which adjusts for bias resulting from copy-number alterations (see Methods) (Kim et al. 2022).

TAD comparisons across samples and replicates (Fig. 1A) show similar TAD numbers (∼4000 TADs) and genomic size (∼0.6Mbp; Supplementary Fig. S3), TAD border insulation strength (R^2^ = 0.77-1.00; Supplementary Fig. S4) and intra-TAD contact frequencies (R^2^ = 0.91-1.00; Supplementary Fig. S5), and genomic positions of both TADs (Jaccard Index [JI] = 0.76-0.80; called at 50kb resolution), and sub-TADs (JI = 0.71-0.75; 10kb resolution) (Supplementary Fig. S6). Inspecting TADs surrounding known breast-cancer related genes (Lee and Muller 2010) reveals minor differences that were not consistently reproducible across replicates (Supplementary Fig. S7-S13). We conclude that TADs are overall similar across the three cell types and are therefore likely not the main oncogenic drivers in this progression model of breast cancer.

**Fig. 1.**
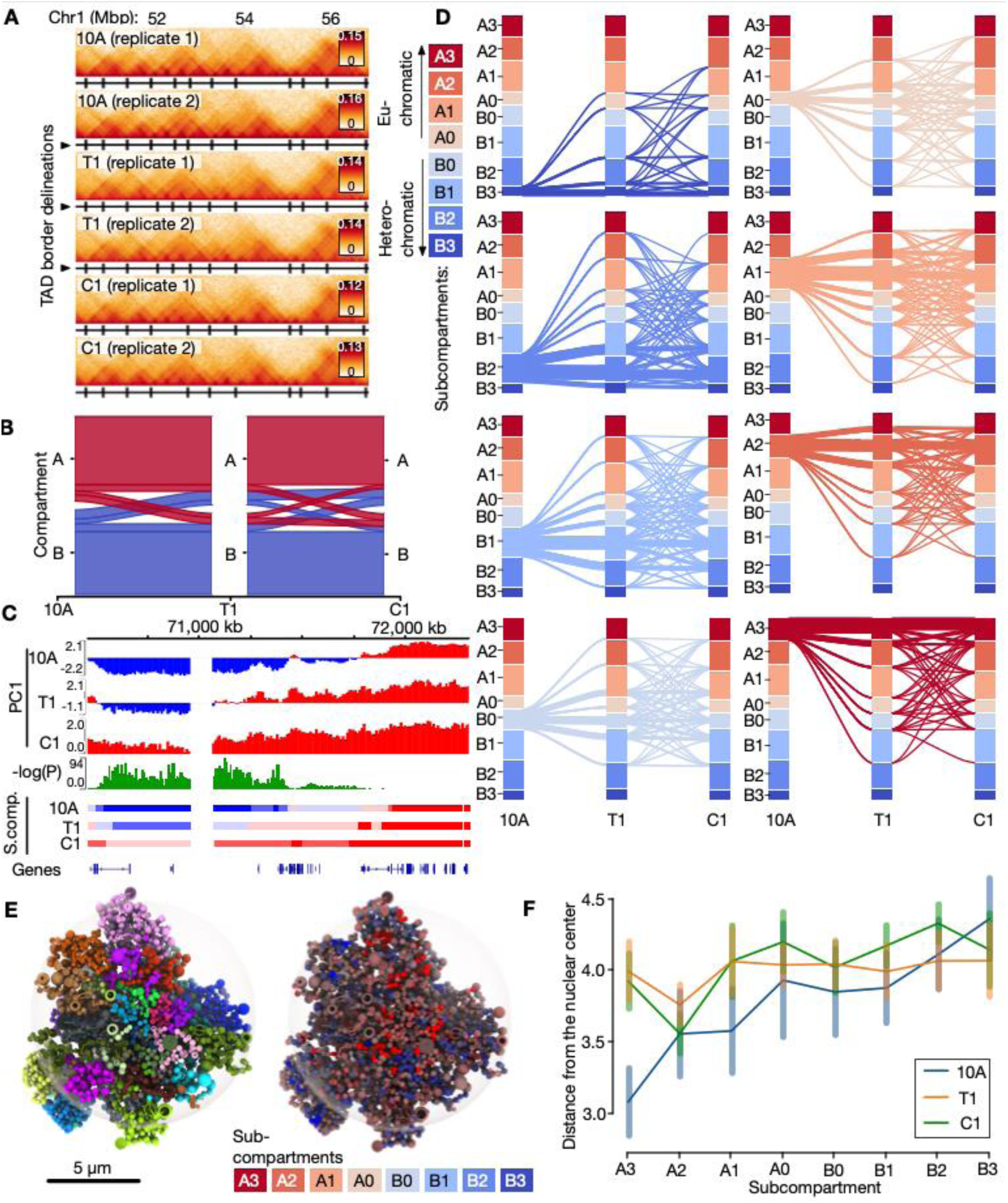
Alterations in genome compartment properties during breast cancer progression stages. **A:** Example Hi-C data for a region on chromosome 1. TAD delineations are shown as black vertical lines below each track. **B**: Alluvial plot showing A/B compartment conservation and switching during progression from 10A (left), via T1 (middle) and to C1 (right). **C**: Example region on chromosome 11 showing a statistically significant switch in the first principal component (PC1) in the three stages. Corresponding subcompartments in the three stages (10A, T1, C1) are shown below. **D**: Switching between four A-type subcompartments (A0-A3), and four B-type (B0-B3) called from the Hi-C data (shown in distinct colors). Subcompartment switches across 10A, T1 and C1 shown as separate alluvial plots starting at each of the eight different subcompartments in 10A. **E**: Left: Tomographic view of exemplary Chrom3D model from 10A cells with chromosomes colored individually. Right: The same model with regions colored by their subcompartment associations. **F**: Median distance (with standard deviation) from the nuclear center for each subcompartment in each condition.

### Compartments and subcompartments engage in major switching events during breast cancer progression

To map differences in long-range genome contacts in our progression model, we searched for any A/B compartment switching, a property reported to be implicated in breast cancer (Barutcu et al. 2015). Most of the genome (2124Mbp; ∼78%) remains in the same compartment across stages (Fig. 1B). However, for the fraction that switches between stages, we note 385 Mbp (∼14%) in 10A switching to B (162 Mbp; ∼6%) or A (222 Mbp; ∼8%) compartments in T1 (Fig. 1B; Supplementary Table S2). Most of the switched compartments remain in their switched state in C1, but a fraction switches back to B (76 Mbp; ∼3%) or A (61 Mbp; ∼2%). The extent of switching observed in our model is comparable to that found in other cellular contexts involving changes in cell fate (Vilarrasa-Blasi et al. 2021; Dixon et al. 2015; Liu et al. 2021), indicating that compartment switching is a prominent feature.

We next used dcHiC to identify subcompartments (Fig. 1C) (Chakraborty et al. 2022). We predict four A sub-compartments (A0-A3) and four B sub-compartments (B0-B3), with A1, A2 and B1 being the most prominent and A0 and B0 the least prominent for all cell stages (Supplementary Fig. S14; Supplementary Table S3). Overlaying a range of active and repressive histone modifications from public ChIP-seq datasets onto the subcompartments reveals a gradual enrichment of the ratio of heterochromatin to euchromatin features, going from B3 to A3 subcompartments (Supplementary Fig. S15). From this analysis, B2 and B3 show features of heterochromatin, with enrichment of repressive histone marks (H3K27me3) and Lamin B1. Compared to the other subcompartments, A0 and B0 are the least enriched in active and repressive histone marks respectively, raising the interesting possibility that they could be more easily remodeled into other stronger euchromatic or heterochromatic subcompartments. In contrast, A3 and B3 display the strongest active and inactive states respectively, which would arguably be more resistant to configuration changes (Supplementary Fig. S15).

Indeed, we find that (i) weaker, intermediate subcompartments (A0, A1, B0, B1) undergo more dynamic switches than more prominent and stronger subcompartments (A2, A3, B2, B3) (Fig. 1D; Supplementary Fig. S16-18). Yet, interestingly, (ii) switching between B0 and A0 (B0↔A0) is minimal; whereas (iii) B0↔A1 is the most prominent inter-compartment switch (Fig. 1D; Supplementary Fig. S16-18), indicating specific switching between weakly heterochromatic and euchromatic states. A further characterization of paths of subcompartment switching across the three stages shows that (iv) a similar fraction of 24-26% of subcompartments ends up in either a more closed (and thus more B-like) subcompartment in C1, or a more open (A-like) subcompartment. Further, reverting back to the initial 10A-subcompartment landscape in C1 rarely happens (Supplementary Fig. S19). Lastly, (v) switches mainly involve a transition between consecutive subcompartment strengths, occurring with similar frequencies towards both more open (e.g. A2→A3) and less open (e.g. B2→B3) subcompartments (Supplementary Fig S19). Comparing the switching levels between the two replicates shows a high degree of consistency, with correlation coefficients (R^2^) ranging from 0.87 to 0.94 (P<1e-20; see Supplementary Fig. S20).

To determine implications of subcompartment switching on 3D genome organization, we generated 3D genome models (Paulsen et al. 2017) integrating Hi-C data with nuclear lamina-chromatin interactions (LADs) mapped by ChIP-seq of lamin B1 in each cell type (see Methods). The resulting models (Fig. 1E; Supplementary Fig. S21-23) display expected genomic features where more gene-poor chromosomes are located more frequently towards the nuclear periphery (Supplementary Fig. S24-26). Strikingly, 3D modeling of all three cell types reveals major differences between 10A cells versus T1 and C1 cells. As expected, there is a gradual increase in nuclear peripheral localization in subcompartments going from A3 towards B3 in 10A cells. This, however, is less apparent in T1 and C1 cells (Fig. 1F). Moreover, A3 subcompartments significantly shift away from the nuclear center in T1 and C1 cells, compared to 10A (Fig. 1F; Supplementary Table S4). B3 subcompartments appear to be slightly shifted towards the nuclear interior, which concurs with a reduction in LAD coverage in T1 and C1 cells (Supplementary Fig. S27). Taken together, our data indicate that subcompartment switching characterizing the 10A-T1-C1 cell transition is accompanied by a partial loss of radial disposition of chromatin.

### Subcompartment switching reflects changes in gene expression

We next investigated whether the 3D genome structural changes observed were linked to transcriptional changes. Differential gene expression analysis from RNA-seq data between the cell types reveals 3180 differentially expressed (DE) genes between 10A and T1 cells, and 8362 DE genes between 10A and C1 cells (|LFC|>0.5; P<0.01; Supplementary Table S5). Analysis of disease-linked gene set enrichment among DE genes in T1 and C1, reveals “breast carcinoma” as the most significant associated disease term, followed by other cancer types. “Organ system benign neoplasm” is the second most associated term for T1, whereas this term is absent for C1, reflecting the expected transcriptomic differences between the transformed (T1) versus the malignant (C1) cell line (Supplementary Fig. S28). Overlaying the RNA-seq data onto Hi-C-based subcompartments in all cell types reveals anticipated associations where gene expression levels gradually increase from B3 to A3 (Fig. 2A, Supplementary Fig. S29), reinforcing the validity of the subcompartment classification and the RNA-seq data in our model.

**Fig. 2.**
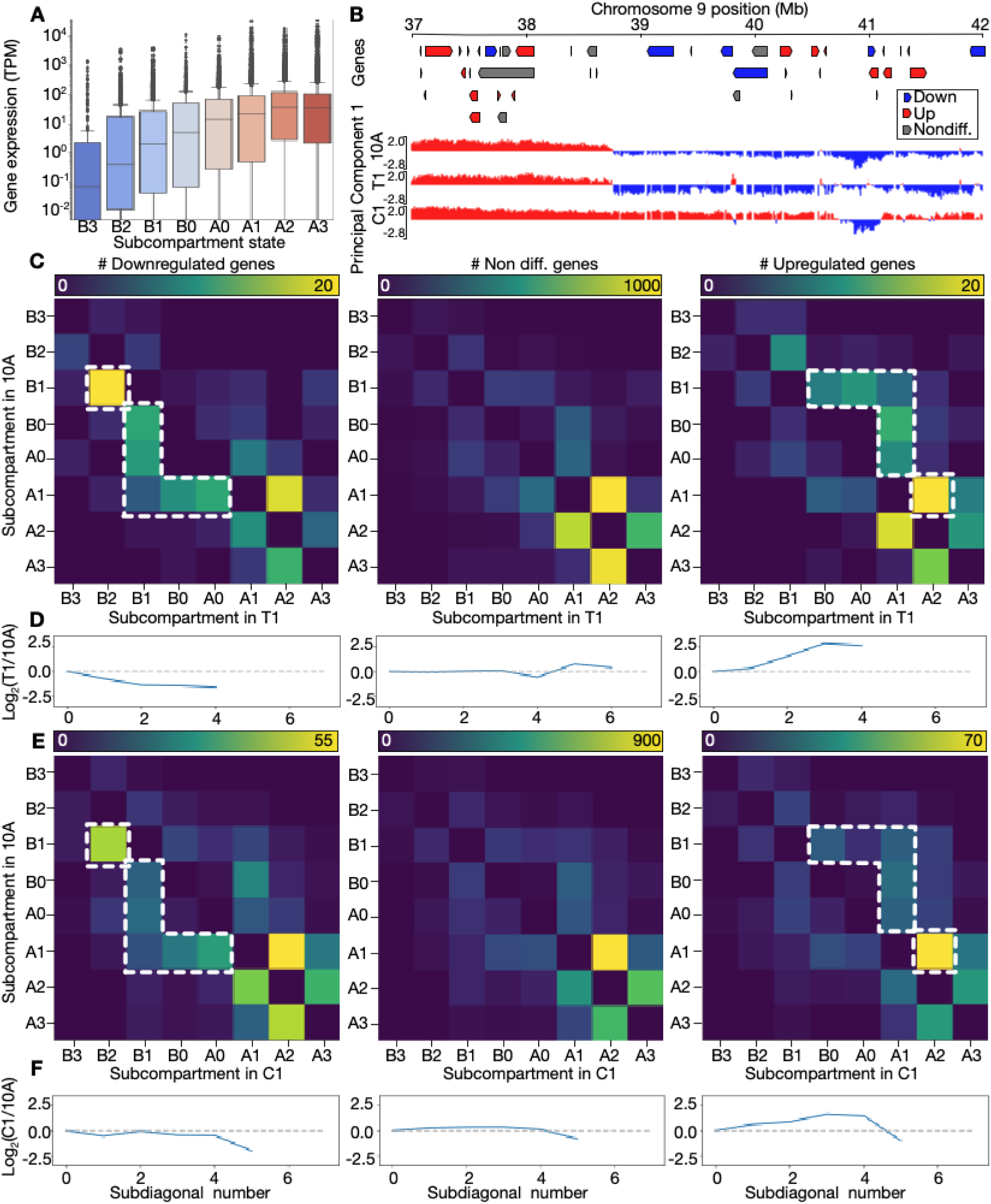
Coordinated changes in subcompartments and gene expression. **A**: Gene expression levels (TPM) by subcompartment state. Vertical axis in log scale. **B**: Example of subcompartment switching on chromosome 9. Up- and downregulated genes in C1 vs. 10A indicated in red and blue, respectively. **C**: Heatmaps showing number of DE downregulated genes (left panel), non-differential genes (middle panel) and upregulated genes (right panel) in subcompartments switching between 10A (vertical axis) and T1 (horizontal axis). Dotted lines highlight regions with enrichment relative to non-differential and upregulated genes. **D**: Log2-ratio of number of genes in subdiagonal sums in the upper vs. lower triangular of the corresponding heatmap from C. Ratio not displayed for subdiagonal sums that are equal to zero. **E**: Heatmaps as in C but contrasting 10A with C1. Dotted lines highlight regions with enrichment relative to non-differential and downregulated genes. **F**: Log2-ratio plots as in D, but contrasting 10A with C1.

Next, we analyzed up- and downregulated DEGs in conjunction with the corresponding subcompartment switches. As expected, and as previously reported (Nagai et al. 2019; Liu et al. 2021), this analysis shows that at specific chromosomal regions (Fig. 2B), or at genome-wide levels (Fig. 2C-F), there are correlated patterns of subcompartment switching and differential gene regulation. Inspecting the presence of downregulated genes in T1 relative to 10A reveals that B1(10A)→B2(T1) is the most frequent switch for these genes (Fig. 2C left panel). Switching between B0/A0(10A)→B1(T1) or from A1(10A)→B1/B0/A0(T1) is enriched relative to what is seen for non-differentially expressed genes (Fig. 2C middle panel) or upregulated genes (Fig 2C right panel).

To quantify whether up- and down-regulated were associated with expected subcompartment switches towards more open (or A-like), or closed (or B-like) subcompartments, respectively, we analyzed enrichment of DE genes in each subcompartment switch relative to a switch in opposite direction. This analysis revealed a significant enrichment of upregulated genes in switching of B1→B2 (P<1e-20; Binomial test), B0→B1 (P<1e-20), A0→B1 (P=0.004) and A1→B0 (P=0.038) between 10A and T1. Downregulated genes were significantly enriched in B1→A1 (P=0.004) and A1→A3 (P=0.019) (Supplementary Table S6). Comparing 10A and C1, downregulated genes were enriched in A0→B2 (P=0.019), B0→B1 (P=0.008) and A1→A0 (P=0.014), whereas upregulated genes were enriched in B3→B1 (P=0.031), B1→A1 (P=0.001), B1→A2 (P=0.013), B0→B2 (P=0.004), A0→A2 (P=0.004), A1→A2 (P<1e-20), A1→A3 (P=0.001) (Supplementary Table S7). This suggests that specific subcompartment switching towards more open or closed states is associated with up- or downregulated DEGs, respectively.

Further, we computed the log-ratio of subdiagonal sums in the upper-compared to the lower triangular matrices from Fig. 2C. When this log-ratio is negative, DE genes are more present in 10A, and when it is positive, DE genes are more present in T1 (Fig. 2D) or C1 (Fig. 2F). We confirm switching trends towards more open subcompartments for upregulated genes, and the opposite for downregulated genes (Fig. 2D right) (P=4.04e-09; Fisher’s exact test; Supplementary Table S8), while no trend is seen for non-differentially expressed genes (Fig 2D middle). When contrasting 10A and C1, similar trends are seen (Fig. 2E-F; P=6.66e-13; Supplementary Table S9). Thus, subcompartment switching aligns with changes in gene expression.

### Reorganization of TAD-TAD interactions coincides with subcompartment alterations

Beyond their subcompartment organization, we have previously shown that TADs can arrange in densely inter-connected long-range contact configurations termed TAD cliques (Paulsen et al. 2019). Briefly, to detect cliques, we view each TAD as a graph node linked by edges indicating significant Hi-C interactions. A “TAD clique” is maximal when it displays a fully-connected subset of nodes that cannot be expanded without losing full connectivity. The “TAD clique size” is determined by the number of nodes it contains in a maximal clique (see Supplementary Fig. S30). We computed TAD cliques in 10A, T1 and C1 cells (see Fig. 3A), and found that the distribution of maximal clique sizes overall show an expected decrease in their frequency as their size increases (Fig. 3B; Supplementary Fig. S31A). Notably, in 10A TAD cliques of size 3 are 17% less frequent than in T1 (P=0.036; Suppl. Table S10), and 24% less frequent than in C1 (P=1.8E-07; Suppl. Table S10). Larger TAD cliques (size>4) on the other hand seem generally more enriched in 10A, than in T1 and C1. T1 generally exhibits fewer TAD cliques than 10A and C1 (Fig. 3B). Therefore, larger clique sizes are more apparent in normal 10A cells. These observed trends and differences are likely not related to structural variants in the T1 and C1, as masking these out reveal similar distributions as in non-masked data (Supplementary Fig. S31)

**Fig. 3.**
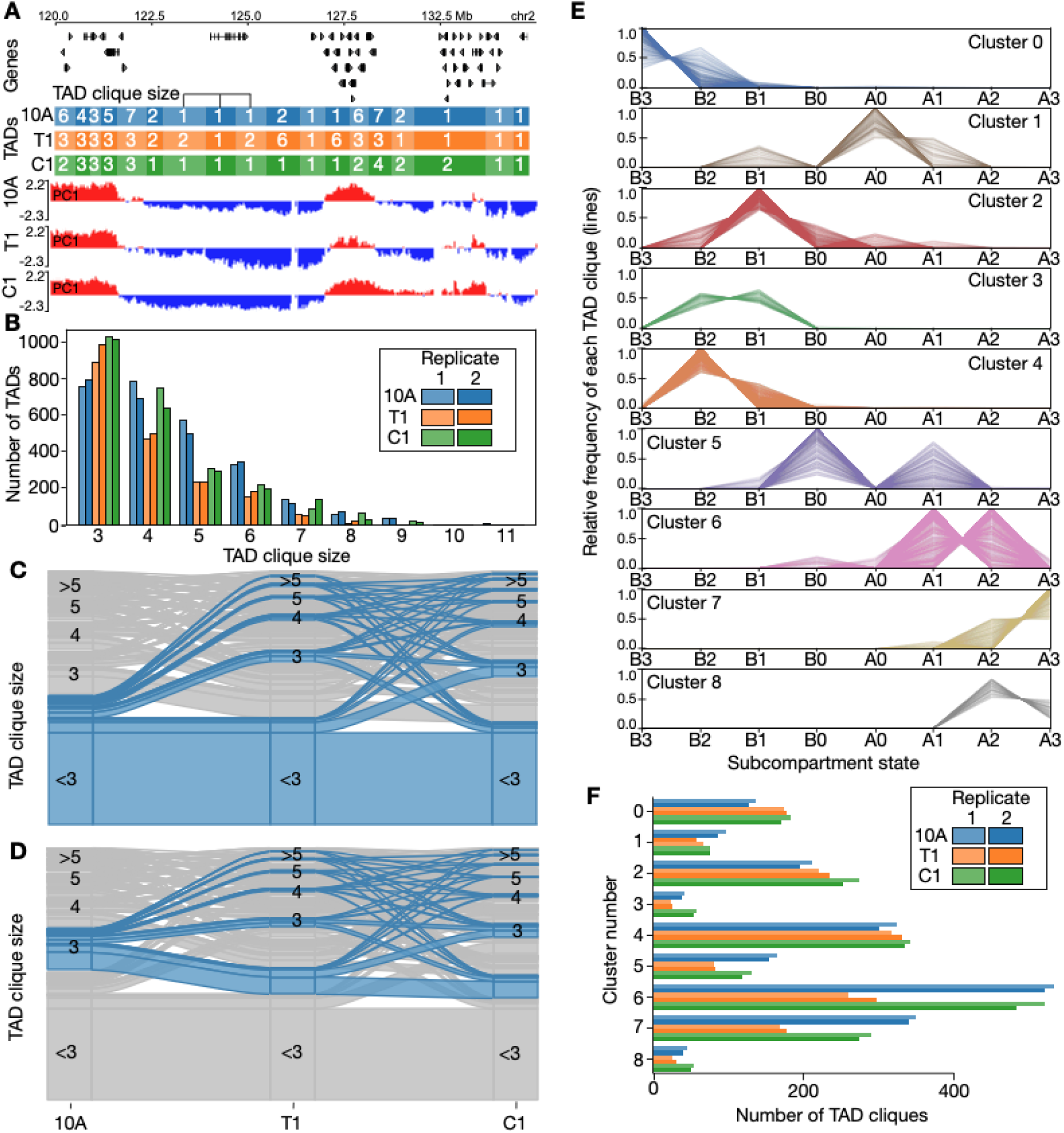
TAD clique dynamics during breast cancer progression stages. **A**: Example region on chromosome 2 showing changes in TAD clique sizes and compartment eigenvalues. **B**: Absolute number of TADs in cliques with maximal size ranging from 3 to 11 shown for each of the three breast cancer stages. **C**: Alluvial plots highlighting the alluvial path of a non-clique in 10A across the three cancer stages. **D**: Alluvial path of a TAD clique of size 3 in 10A. **E**: TAD clique clustering into 9 clusters based on their enrichment in subcompartments in the same cell type (vertical axis). Each line represents a TAD clique. **F**: Numbers of TAD cliques belonging to each of the clusters in D.

We overlapped TAD cliques of different sizes with subcompartments, and observed that association of B-type subcompartments increases with TAD clique size (Supplementary Fig. S32). To characterize TAD clique dynamics, we analyzed alterations in TAD maximal clique sizes across cell types (Fig. 3C-D; Supplementary Fig. S33-S39). A large fraction of TADs do not engage in cliques (Fig. 3C). However, those TADs that do form cliques grow or decrease in size (Fig. 3C-D; Supplementary Fig. S33-S39) across all stages. Thus, like subcompartments, TAD cliques also reconfigure extensively in this model system of breast cancer progression.

Our previous observations do not exclude the possibility that growing TAD cliques involve TADs of various chromatin composition, thus belonging to various subcompartments. To explore this, we used HDBSCAN (McInnes et al. 2017; McInnes and Healy 2017) to cluster each clique based on their TAD-wise enrichment in subcompartments. Clustering was performed targeting clusters with a minimum cluster size of 200 TAD cliques, plus a cluster dedicated to collect outliers (see Methods for more details). The resulting nine clusters (Fig. 3E) show that individual TAD cliques frequently involve multiple types of subcompartments (Supplementary Fig. S40), and thus represent a level of TAD association where subcompartments of different types potentially intermix.

To investigate the dynamics of TAD clique clusters, we explored the frequency of each of the nine TAD clique clusters in each cell type (Fig. 3F; Supplementary Table S11). We find that most clusters display a cancer progression stage-specific association, such that a cluster tends to be more enriched in one or two of the three cell types examined here (Fig. 3E,F). Some clusters involving B-type subcompartments (Cluster 0 and 2) appear to be significantly more prevalent in T1 and C1 compared to 10A. Conversely, some clusters with A-type subcompartments (Cluster 1, 5-7) seem to be diminished in T1, but exhibit partial recovery in C1 (Fig. 3F; Supplementary Table S12).

In summary, TAD cliques bring together TADs overlapping with various subcompartments, which differ in 10A, T1 and C1 cells. Overall, this signifies the role of subcompartment alterations and intermixing during breast cancer progression in this model system.

### Interchromosomal insertion of the *MYC* locus coincides with de novo enhancer contacts and oncogene activation

As expected, the cell types used in our breast cancer progression model harbor structural variations (SVs), including copy number alterations, insertions, deletions and translocations. We exploited the ability of Hi-C to detect many of these SVs, to call SVs genome wide (Supplementary Fig. S41-S43). We manually inspected the called variants and confirmed known SVs (Santner et al. 2001), including t(3;9), t(3;5), t(3;17), t(6;19). In addition, we identified previously undescribed SVs, including t(7;9) and t(10;17). Analyzing the 3D structural consequences of all SVs in these cell lines is computationally infeasible. Nevertheless, to investigate 3D genome consequences of select SVs, we focused on a striking pair of regions on chromosome 8 (126330000-128235000 bp; hg38) (Fig. 4A) and chromosome 10 (71280000-73310000 bp; hg38) (Fig. 4B). Both regions show a gradual and coordinated increase in copy number, going from T1 to C1 (Fig. 4A,B). A large region at the end of chromosome 8 is amplified in 10A, but not in T1 and C1 (Fig. 4A). The region on chromosome 8 is well-characterized (Visscher et al. 1997; Grisanzio and Freedman 2010) and contains several breast cancer related genes. These notably include *MYC*, an oncogenic transcription factor with a pivotal role in breast cancer progression (Liao and Dickson 2000) and whose amplification and overexpression is a marker of aggressive and invasive breast cancer (Corzo et al. 2006; Berns et al. 1992).

**Fig. 4.**
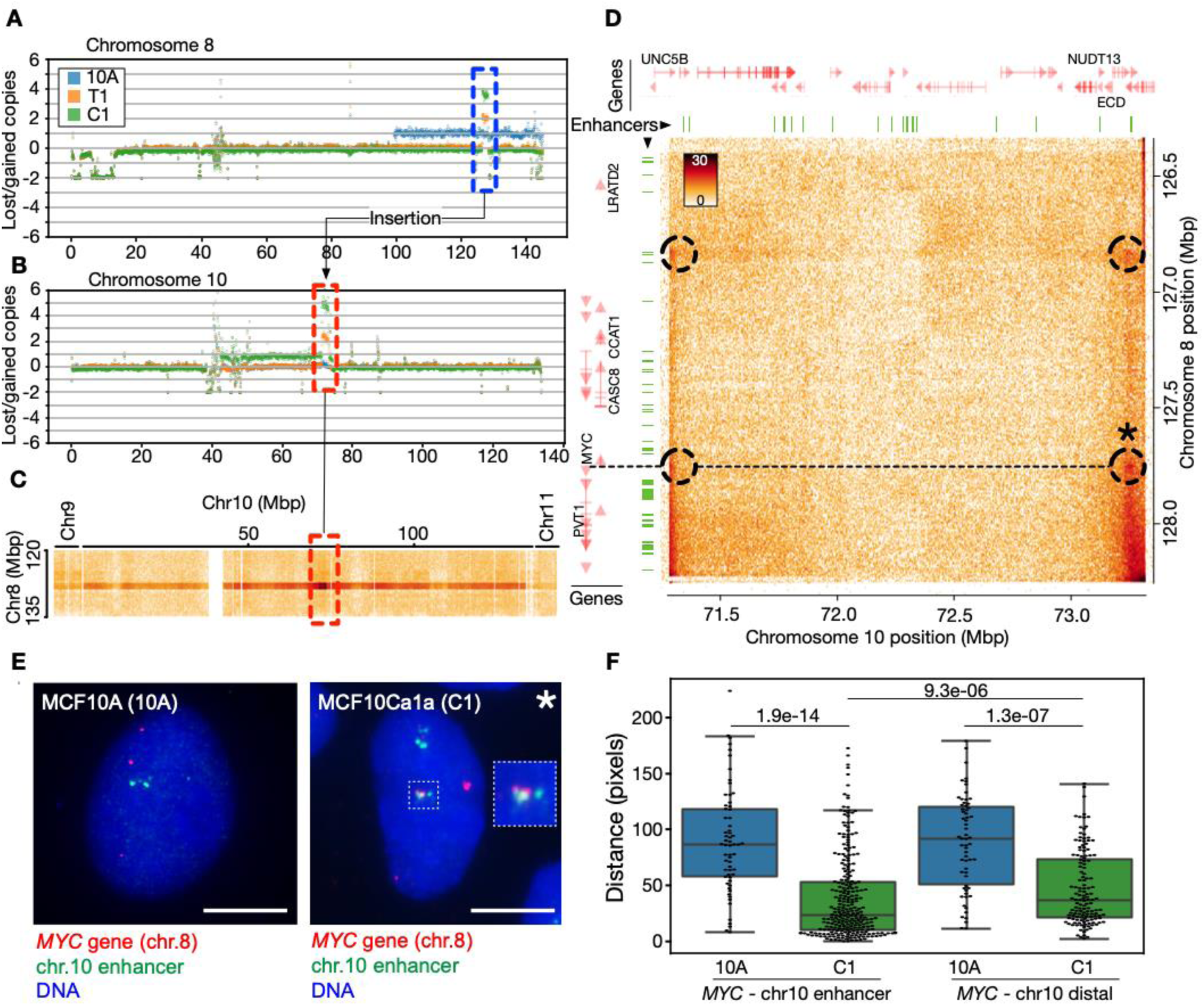
*MYC* locus insertion on chromosome 10 is accompanied by *de novo* contacts with a potential enhancer element. **A**: Lost and gained copies for the entire chromosome 8 in 10A, T1, and C1 cells. The dotted blue box highlights specific amplification of the chromosome 10 region. **B**: Lost and gained copies for the entire chromosome 10, with a dotted red box highlighting specific amplifications. **C**: Interchromosomal C1 Hi-C contacts between the region on chromosome 8 (vertical axis) and the entire chromosome 10. Highlighted region from B indicated in red. End of chromosome 9 and beginning of chromosome 11 shown on the left and right side, respectively. **D**: Zoom-in on the C1 Hi-C map of the chromosome 8/10 amplification unit. Dotted circles indicate enriched “dots” of contacts involving enhancers and *MYC*. Asterisk (*) marks the enhancer-interaction validated in FISH in panels E-F. The dotted line shows the position of the *MYC* gene on chromosome 8 relative to contacts on chromosome 10 within the amplification unit. Positions of enhancers from MCF10A indicated as green segments (top and left). Genes shown in red (top and left), with names for selected genes. **E**: FISH image of the *MYC* gene (red) and the chromosome 10 enhancer (green; chr.10 position 73117711-73267436) expected to interact with MYC from the Hi-C data, in 10A and C1 cells. Bars, 7 µm. **F**: Quantification of probe-probe distances in pixels, for the *MYC*-chr10 enhancer probe pair (left; n = 62 [10A] and 269 [C1] observations) and the *MYC*-chr10 distal probe pair (position 73794997-73992612; n = 62 [10A] and 140 [C1] observations); N = 28-95 nuclei analyzed; P-values are shown (Kolmogorov test).

Further inspection of the Hi-C data at the intersection of these two regions in the Hi-C matrix reveals, after ICE balancing, an enrichment of contacts of the region towards the entire chromosome 10, clearly indicating that the *MYC* locus is inserted in the chromosome 10 region (Fig. 4C). Enriched contacts with other chromosomes are also seen, due to the amplification of the locus (Fig. 4C). Zooming in on the specific interchromosomal contacts between the *MYC*-locus and the locus on chromosome 10, we note Hi-C patterns that typically suggest chromatin looping and extrusion of intra-chromosomal DNA (Fig. 4D). This again indicates an insertion event. Investigating the region using a different balancing scheme which explicitly models and accounts for copy number alterations (Servant et al. 2018) reveals similar trends. This indicates that the copy number alterations themselves are not distorting the Hi-C data in this region (Supplementary Fig. S44).

We next examined the chromatin context of this insertion event. Inspection of subcompartments in this region shows that the insertion site on chromosome 10 is nearly entirely covered by A3-subcompartments (Supplementary Fig. S45A) in all three cell types, indicating that the *MYC* locus is inserted into a highly active genome region on chromosome 10. Initially, the *MYC* gene resides in an A2 subcompartment on chromosome 8 in the 10A cell line. However, it switches to a more active or “open” state (A2→A3) in T1 and C1 (Supplementary Fig. S45B).

Next, we assessed the nature of the *de novo* contacts detected between the *MYC* locus and sites on the host chromosome 10. To this end, we overlaid positions of MCF10A enhancers predicted from EnhancerAtlas 2.0 (Gao and Qian 2020) onto the chromosome 8 and 10 amplification unit (Fig. 4D; green segments). Strikingly, the four distinct *MYC* interactions detected in the Hi-C map (Fig. 4D, circles) evidently corresponds to (i) two *de novo* contact points between enhancer elements on chromosome 10 and the *MYC* promoter, and (ii) an interaction with enhancer elements ∼1 Mbp upstream of the *MYC* promoter (Fig. 4D).

To validate the *de novo* interactions (proximity) of the *MYC* gene with elements on chromosome 10 at the cellular level, we carried out a fluorescence in situ hybridization (FISH) analysis of the relative positions of the *MYC* gene (*‘MYC’* probe on chromosome 8 at position 127543567-127702694) and either a *de novo* interacting enhancer on chromosome 10 (“chr10 enhancer” probe at position 73117711-73267436), or another site on chromosome 10 ∼500 kbp downstream (“chr10 downstream” probe at position 73794997-73992612; see Methods). The data show that both the *“MYC*-chr10 enhancer” and the “*MYC*-chr10 distal” distances are significantly shorter in C1 cells than in 10A cells (P=1.9e-14 and 1.3e-07 respectively; Fig. 4E,F; Supplementary Fig. S46A-D), yet the *MYC* enhancer contact is significantly shorter than the distal interaction (P=9.3e-06). These findings strongly support the view that the *MYC* locus has undergone an insertion into chromosome 10 in C1 cells and engages with a nearby enhancer element.

Lastly, to provide an element of functional significance to this *MYC* insertion, we examined RNA-seq data for these two regions. The data show an upregulation of breast cancer related genes in T1 and C1 relative to 10A, including *LRATD2, PCAT1, CASC19, CASC8, POU5F1B, PVT1* and *MYC* on chromosome 8. Several breast cancer related genes in the chromosome 10 region are also upregulated, such as *UNC5B, CDH23, PSAP, SPOCK2, ASCC1, DDIT4, NUDT13, ECD* and others (Supplementary Tables S13 and S14).

Collectively, these observations indicate that the *MYC* locus has not only been amplified but also inserted into a transcriptionally active region of chromosome 10 in C1 cells, with the formation of de novo cis-contacts between *MYC* and regulatory elements in the host chromosome. Up-regulation of *MYC* and of other breast cancer-related genes in malignant C1 cells speculatively suggests functionality of these de novo interactions. Our results highlight how structural variants in the genome may cooperate with 3D genomic states to drive oncogenesis.

## Discussion

The intricate 3D organization of the genome plays a pivotal role in gene regulation and cell function. Using a multilayered (epi)genomic approach, we report here for the first time, to our knowledge, significant alterations in higher-order 3D chromatin organization in an *in vitro* breast cancer progression model.

First, major compartment and subcompartment switching emerges as a prominent dynamic feature, which as expected based on previous studies (Nagai et al. 2019; Liu et al. 2021), correlates with altered expression of genes implicated in breast cancer. Second, we identify a reconfiguration of preferred associations between TADs, or TAD cliques (Paulsen et al. 2019), which bring together new A- and B-type subcompartment associations. Third, spatial sub-compartment organization changes in the pre-malignant and malignant states potentially also contributing to the loss of gene expression control. Collectively, there is a coordinated and progressive reorganization of higher-order genomic structure to an abnormal state. However, casual relationships remain to be determined.

The extent of compartment switching observed in our model (22%) is similar to what is seen during B-cell differentiation (28.1%) (Vilarrasa-Blasi et al. 2021), although the latter includes an additional intermediate compartment, inflating the percentage. In contrast, only 7.5% of A and B compartments switch during differentiation of mouse embryonic stem cells (ESCs) into neuronal progenitors (Chakraborty et al. 2022). A/B-compartment switching varies between 8-36% comparing various ESC-derived cell types (Dixon et al. 2015; Liu et al. 2021). Comparatively, this indicates that the level of switching observed in our oncogenic model is substantial.

Our study revealed no significant role of TAD border dynamics in the in vitro progression of breast cancer. However, we cannot entirely rule out its subtle involvement due to minor differences in consistency across replicates. Additionally, the potential role of cell heterogeneity in cancer cells cannot be dismissed, given that our data represent an overall behavior in bulk cell populations (Zhu et al. 2023).

A new observation from our data is a previously unknown insertion of the *MYC* locus (normally localized on chromosome 8) into highly active A3-subcompartments on chromosome 10. This is accompanied by *de novo* enhancer-promoter interactions correlating with not only elevated *MYC* expression in T1 and C1 cells, but also with enhanced expression of other breast cancer related genes on both chromosomes. Importantly, our FISH analysis strongly supports our Hi-C findings of *de novo* interactions between *MYC* and an enhancer element on chromosome 10 in C1 cells, but not in 10A cells. This also supports a *MYC* locus insertion event. Notably, in murine B-cells, *MYC* has been shown to relocate to active transcription factories in proximity to *IGH,* a frequent translocation partner (Osborne et al. 2007). However, our FISH results indicate that in 10A cells, *MYC* is not proximal to the chromosome 10 locus, suggesting *MYC* is not part of a transcription factory with this locus prior to the insertion event.

The detailed spatial configuration of this insertion remains currently unclear, but there are several possibilities. The mirror-symmetry of contacts in the chromosome 10 segment with specific regions of the chromosome 8 segment (see Fig. 4D), could suggest that the region exists as circular, extrachromosomal DNA (ecDNA). The formation of ecDNA structures is a common form of oncogenic amplification, and often occurs around *MYC (Hung et al. 2022; Hoff et al. 1988)*. Our observation that the region exists essentially as an isolated contact map in the interchromosomal Hi-C data (see Fig 4D and Supplementary Fig. S44) further supports the possibility of an ecDNA containing *MYC* and interacting regulatory elements. Interestingly, ecDNA has been shown to promote functional cis-regulatory contacts (Morton et al. 2019), possibly through formation of ecDNA hubs (Hung et al. 2021), to drive oncogenesis. Our FISH data, however, cannot currently resolve whether *MYC* is inserted into chromosome 10 as an integral segment of the host chromosome or whether ecDNA structures also exist.

Beyond our identified interchromosomal insertion of the *MYC* locus, the intricate aneuploidy patterns commonly observed in cancer pose a challenge that existing standard software and pipelines struggle to effectively address. Promising strides have been made using long-read Hi-C technology (Garg 2023) or statistical approaches (Brunette et al. 2024) to alleviate this issue. Nevertheless, a significant gap remains in the availability of computational pipelines tailored for comprehensive downstream analysis of reordered cancer genome data relative to a karyotypically normal reference genome. This needs to be addressed if further progress in understanding the 3D cancer genome is to be achieved.

In conclusion, our findings provide new insights into the complex genomic structural changes that underlie breast cancer metastatic progression. The interplay between subcompartment switching, LAD-linked subcompartment spatial nuclear reorganization, and TAD clique dynamics represents a multifaceted mechanism by which the 3D genome can be abnormally and dynamically reconfigured during breast cancer development. Targeting these 3D genome alterations could potentially lead to improved or new therapeutic strategies to better treat breast cancer in the future (Park et al. 2023).

## Methods

### Cells

MCF10A, MCF10AT1, and MCF10Ca1a cell lines were grown in DMEM/Nutrient F12 (DMEM/F12) media supplemented with 5% horse serum (MCF10A and MCF10Ca1a) or 2.5% horse serum (MCF10AT1), 14 mM NaHCO3, 10 µg/mL insulin, 2 mM L-glutamine, 20 ng/mL human epidermal growth factor, 500 ng/mL Hydrocortisone and 100 ng/mL cholera Toxin. MCF10A cells were obtained from the American Type Culture Collection (CRL-10317). MCF10AT1 and MCF10Ca1a cells were obtained from the Barbara Ann Karmanos Cancer Institute (Detroit, Michigan).

### Fluorescence in situ hybridization (FISH)

FISH was done as exactly described by us previously (Paulsen et al. 2017). In short, MCF10A and MCF10Ca1a cells were incubated in hypotonic buffer, fixed in ice-cold methanol:acetic acid and dropped on glass slides. BAC FISH probe DNA (BacPac Resource Center) was labeled using a Nick Translation Kit (Roche). The following probes were used:

● ‘MYC gene’ probe: clone ID RP11-1136L8; chr.8; position 127543567-127702694
● ‘Chr.10 enhancer’ probe: clone ID RP11-152N13; chr.10; position 73117711-73267436
● ‘Chr.10 distal’ probe: clone ID RP11-390A15; chro.10; position 73794997-73992612

The MYC gene probe was labeled with Digoxigenin-11-dUTP (Roche). The chr.10 enhancer and chr.10 distal probes, both positioned to mark sites expected to be proximal to the MYC gene on chromosome 10) were labeled with Biotin-16-dUTP (Roche). For each slide, 200 ng of each ‘MYC gene’ + ‘chr.10 enhancer’ probe, or ‘MYC gene’ + ‘chr.10 distal’ probe were mixed with 30 µg of Cot-1 DNA and 150 µg salmon sperm DNA and precipitated. DNA was dissolved in hybridization mix and pre-annealed for 1 h. Slides were RNase-treated, washed, dehydrated in ethanol, denatured, and dehydrated again.

Probes were denatured, pre-annealed and applied onto cells for overnight hybridization at 37°C. Slides were then washed in 2x SSC and in 0.1x SSC, blocked in 5% skim milk and incubated with Anti-Digoxigenin (Roche; mouse; 0.4 µg/ml). Slides were washed, incubated with Avidin Alexa Fluor 488 conjugate (Invitrogen; 1.7 µg/ml) and Alexa Fluor® 594-conjugated Anti-Mouse (Jackson ImmunoResearch; rabbit; 2.5 µg/ml), washed and incubated with Biotinylated Anti-Avidin D conjugate (Vector; goat; 1.0 µg/ml) and Alexa Fluor® 594-conjugated Anti-Rabbit (Jackson ImmunoResearch; donkey; 2.5 µg/ml). Slides were washed and incubated 30 min with Avidin Alexa Fluor 488 conjugate (Invitrogen) (1.7 µg/ml). Slides were mounted with 0.2 µg/ml DAPI in Dako Fluorescent Mounting Medium. Images taken under a 100x objective (numerical aperture 1.4) mounted on an IX71 inverted microscope (Olympus) fitted with the DeltaVision wide-field imaging station (GE Healthcare).

### Hi-C

HiC Libraries were prepared in duplicates for each cell type using the Arima-HiC+ kit (Arima Genomics, USA) according to manufacturer instructions. For each cell line (10A, T1, C1) 1 million cells were used per HiC reaction. Cells were fixed with 1.2% formaldehyde for 12 min. All subsequent steps were carried out according to the Arima protocol. Resulting libraries were amplified with 5 PCR cycles and sequenced on an Illumina NovaSeq 6000 instrument in paired-end run with 2 x101 bp.

### RNA-seq

All mRNA-Seq experiments were performed in triplicate. Total RNA was isolated using the Qiagen RNeasy kit following manufacturer’s instructions. Stranded mRNAseq libraries were constructed using Illumina mRNA Prep kit, following vendor protocol with poly-A enrichment (Illumina). Libraries were sequenced with 2x75bp paired-end on an Illumina NextSeq 500.

### ChIP-seq of lamin B1 and mapping of lamina-associated domains (LADs)

ChIP of lamin B1 was done as described by us (Rønningen et al. 2015). In short, cells were cross-linked with 1% formaldehyde, lysed in 50 mM Tris-HCl pH 7.5, 10 mM EDTA, 1% SDS and protease inhibitors, and sonicated in a Bioruptor (Diagenode) into ∼200-bp fragments. After sedimentation, the supernatant was diluted tenfold in RIPA buffer. Chromatin was incubated overnight at 4°C with antibodies to lamin B1 (10 μg per 10 million cells; Abcam, ab16048), coupled to Invitrogen Dynabeads protein A/G (Thermo-Fisher). ChIP samples were washed four times in ice-cold RIPA buffer, after which cross-links were reversed and the DNA was eluted for 6 h at 68°C. DNA was purified using phenol–chloroform isoamylalcohol and dissolved in H2O. ChIP-seq libraries were prepared using the Diagenode Microplex library preparation kit v2 and TruSeq LT indexes, and samples were sequenced (single-end) on an Illumina HiSeq4000.

ChIP sequence reads were mapped to hg38 with Bowtie2 v2.4.1 (https://github.com/BenLangmead/bowtie2) after removing duplicates using Picard MarkDuplicates (http://broadinstitute.github.io/picard/). Reads from both input samples were merged, mapped to hg38 and duplicates removed as above. To alleviate normalization bias, each pair of mapped ChIP and input read files contained the same read depth after down-sampling reads for each chromosome. Mapped reads were used to call LADs from ten consecutive runs of Enriched Domain Detector (EDD) (http://github.com/CollasLab/edd) (Lund et al. 2014) with auto-estimation of GapPenalty and BinSize, and mean GapPenalty and BinSize values were used for a final EDD run (Forsberg et al. 2019). Final LADs for each cell type were the union of the three replicates.

### Structure of code used for data analyses

Source code used to produce all results presented in this paper (except LADs analysis and downstream analysis of Chrom3D models) is hosted on GitHub at github.com/paulsengroup/2022-mcf10a-cancer-progression and is archived on Zenodo at doi.org/10.5281/zenodo.13255896.

Most of the computation is organized into Nextflow workflows, with each workflow performing a subset of the data analysis (e.g. the differential expression and comparative analyses are performed by two different workflows).

All workflows mentioned in the method sections are hosted on the GitHub repository inside folder workflows/. Scripts and Jupyter notebooks are found under folders bin/ and notebooks/ respectively.

Some of the analysis steps required us patching third party tools. The patches are described in section Software patches and patch files are available on the GitHub repository under container/patches/. All our patches were contributed upstream. Most of our patches were accepted by upstream and are already part of a stable release. Figures were produced directly by workflows or using scripts and notebooks under bin/plotting and notebooks/, respectively. The final version of figures were produced by assembling draft images with Inkscape and Omnigraffle.

Data analysis workflows were run on a HPC cluster using Apptainer (Kurtzer et al. 2017).

### Hi-C preprocessing

We used a modified version of nf-core/hic v2.0.0 (https://zenodo.org/records/2669513) (Ewels et al. 2020; Servant et al. 2015; Ewels et al. 2016; da Veiga Leprevost et al. 2017) for quality control, sequence mapping and filtering using hg38.p14 (Lander et al. 2001). The workflow was run using restriction enzymes for the 2-enzyme Arima Kit (--restriction_site=”^GATC,G^ANTC” and -- ligation_site=”GATCGATC,GANTGATC,GANTANTC,GATCANTC”). The following optional flags were specified: --skip_maps, --skip_dist_decay, --skip_tads, --skip_compartments, -- skip_balancing, --skip_mcool, --split_fastq=false. The .chrom.sizes file given to nf-core/hic was filtered to only retain chromosome sequences using grep ‘^chr[[:digit:]XY]\+[[:space:]]’. Chromosomes were sorted by name using gnu sort -V. The output of nf-core/hic was compressed using workflow robomics/compress-nfcore-hic-output v0.0.1 (https://zenodo.org/records/7949266). Hi-C contact matrices in multi-resolution Cooler format were generated using a patched version of cooler v0.9.1 (Abdennur and Mirny 2020). We used cooler cload to ingest interactions in .validPair format into a .cool file at 1 kbp resolution. We then converted the .cool file to .mcool file format using cooler zoomify. Finally, matrices were balanced with several methods using juicer_tools v2.20.00 (Durand et al. 2016) (VC, KR, SCALE methods using intra-chromosomal, inter-chromosomal and genome-wide interactions), and cooler (ICE method using intra-chromosomal, inter-chromosomal and genome-wide interactions) (Abdennur and Mirny 2020). Regions overlapping centromeres and assembly gaps for hg38 were masked before balancing (files retrieved from UCSC FTP server on 2023/02/24). Downstream analyses were performed using cis-only, ICE balanced matrices unless otherwise specified. For some analyses, replicates for the same condition were merged to generate deeper Hi-C matrices using cooler merge (Abdennur and Mirny 2020). The resulting matrices were then coarsened and balanced cooler and juicer_tools as outlined above (Abdennur and Mirny 2020; Durand et al. 2016). All the above steps were performed by running script run_nfcore_hic.sh, which runs workflow postprocess_nfcore_hic.nf after running nf-core/hic. Fig. S1 was generated by workflow postprocess_nfcore_hic.nf based on the output of nf-core/hic. Fig. S2 was generated by workflow comparative_analysis_hic.nf by running the Python implementation of hicrep v0.2.6 (Lin et al. 2021; Yang et al. 2017). Correlation values shown in the plot were computed as the weighted average of the correlation coefficient of each individual chromosome using chromosome sizes as weights.

In addition, interactions involving chromosome 8 and chromosome 10 were balanced with LOIC using the iced package (v0.5.10) (Servant et al. 2018). LOIC balancing is performed in one of the steps of the comparative_analysis_hic.nf workflow.

### RNA-seq preprocessing

We used nf-core/rnaseq v3.12.0 (https://zenodo.org/records/7998767) to perform quality control, trimming, alignment and generating the gene expression matrix (Ewels et al. 2020; Di Tommaso et al. 2017) (Ewels et al. 2020; Di Tommaso et al. 2017); This includes the following tools: featureCount (Liao et al. 2014); GffRead (Pertea and Pertea 2020); MultiQC (Ewels et al. 2016); preseq (Daley and Smith 2013); Qualimap 2 (Okonechnikov et al. 2016) ; RSeQC (Wang et al. 2012); Salmon (Patro et al. 2017); SortMeRNA (Kopylova et al. 2012); STAR (Dobin et al. 2013); StringTie2 (Kovaka et al. 2019); UCSC Tools (Kent et al. 2010); DESeq2 (Love et al. 2014); dupRadar (Sayols et al. 2016); Tximeta (Love et al. 2020); Bioconda (Grüning et al. 2018) ; Biocontainers (da Veiga Leprevost et al. 2017) and Apptainer (Kurtzer et al. 2017). nf-core/rnaseq was run using hg38.p14 as reference genome (Lander et al. 2001). The FASTA file used as reference was first filtered using SeqKit v2.5.1 (Shen et al. 2016) to remove mitochondrial chromosomes and unplaced scaffolds by using the following pattern ’^chr[XY\d]+$’. We used the genome annotation for hg38 from Gencode v43 (Frankish et al. 2019). The pipeline was launched using the following optional flags: --aligner=star_salmon, -- pseudo_aligner=salmon, --extra_salmon_quant_args=’--seqBias --gcBias’, --gencode=true. The workflow was run by launching script run_nfcore_rnaseq.sh.

### ChIP-seq analyses of public datasets

We used nf-core/chipseq v2.0.0 to perform quality control, mapping, filtering and peak calling of publicly available ChIP-seq datasets from MCF10A cells (Ewels et al. 2020; Di Tommaso et al. 2017). This includes the following tools: BWA (Li and Durbin 2009); BEDTools (Quinlan and Hall 2010); BamTools (Barnett et al. 2011); deepTools (Ramírez et al. 2016); featureCounts (Liao et al. 2014); HOMER (Heinz et al. 2010); MultiQC (Ewels et al. 2016); phamtompeakqualtools (Landt et al. 2012); preseq (Daley and Smith 2013); SAMtools (Li et al. 2009); UCSC Tools (Kent et al. 2010); DESeq2 (Love et al. 2014); Bioconda (Grüning et al. 2018); BioContainers (da Veiga Leprevost et al. 2017) and Apptainer (Kurtzer et al. 2017). The pipeline was run using hg38.p14 as reference genome from UCSC (Lander et al. 2001) and the gene annotation from Gencode v43 (Frankish et al. 2019). BWA was used for mapping and effective genome size for MACS2 was computed using unique-kmers.py from khmer v2.1.1 (Li and Durbin 2009; Zhang et al. 2008; Crusoe et al. 2015; Döring et al. 2008). Option--narrow_peak was specified when processing ChIP-seq datasets for CTCF and p53. The workflow was launched by executing script run_nfcore_chip.sh.

### Calling of copy number variations

Copy number variations (CNVs) were called from Hi-C data using HiNT-CNV v2.2.8 (Wang et al. 2020).

The list of restriction sites for the Arima 2-enzyme kit was generated by running script bin/compute_restriction_sites_for_hint.py on hg38.14 (Lander et al. 2001).

Reference data and background matrices used by HiNT were downloaded from compbio.med.harvard.edu/hint and have been archived on Zenodo at the following DOI: 10.5281/zenodo.13255302.

Data analysis steps are defined in workflow detect_structural_variants which was run by launching script run_detect_structural_variants.sh.

### Annotation of structural variations

Large structural variations such as translocations were manually annotated by us by looking for square regions of enriched interactions and sharp transitions in the trans portion of the Hi-C matrix (see Supplementary Tables S15-17).

### TAD analyses

We determined genomic positions of topologically associated domains (TADs) in all samples with HiCExplorer v3.7.2 using default parameters on matrices at 10, 20, 50 and 100 kbp resolutions (Ramírez et al. 2018; Wolff et al. 2018, 2020). Workflow tad_analysis.nf was used to generate the draft figures that went into Supplementary figures S3-6. The same workflow was also used to run HiCExplorer. Draft figures for Fig. 1A was generated using notebook bin/plotting/plot_tads_higlass.ipynb. Supplementary Fig. S5 was generated by fetching interactions at 50 kbp resolution for each 10A TAD across cell types, masking values overlapping the first 150 kbp around the diagonal. Finally interactions were summed and normalized by TAD size (as in, the number of pixels overlapping a TAD). To generate Supplementary Fig. S6, TADs from two cell types were paired based on the highest overlap (Jaccard Index of genomic coordinates), and the distribution of pairwise overlaps were plotted. TADs overlapping or around regions masked by matrix balancing were not considered, as the insulation score can be unreliable in these regions.

The analysis is encapsulated by workflow tad_analysis.nf which was run by launching script run_tad_analysis.sh.

### TAD clique analyses

The TAD annotation generated by HiCExplorer was given as input to workflow github.com/robomics/call_tad_cliques v0.5.1 (doi.org/10.5281/zenodo.12689353) to identify cliques of TADs as outlined in Paulsen et al. (Paulsen et al. 2019). TAD cliques were called across all three cell types using TADs from 10A. This is a necessary simplification required to allow comparing cliques across cell types in downstream analyses. The output of github.com/robomics/call_tad_cliques was used as input for workflow postprocess_call_tad_cliques.nf, which generated the draft version of Fig. 3B, 3C, 3D and supplementary figures S31,S33-39.

Workflows were launched by running script run_call_tad_cliques_workflow.sh.

### Subcompartment analyses

Subcompartments were determined using a patched version of dcHiC d4eb244 (Chakraborty et al. 2022). A and B compartment annotation was generated from subcompartments by aggregating A-subcompartments and B-subcompartments. The dcHiC processing pipeline was wrapped in workflow compartment_analysis.nf, which converts matrices in. mcool file to a format understood by dcHiC and runs all analysis steps except those for --pcatype=trans, --pcatype=fithic, --pcatype=dloop, and --pcatype=enrich. When appropriate, the reference genome assembly, annotation and .chrom.sizes for hg38.p14 were provided to dcHiC through the --gfolder option. We fixed the seed used by dcHiC to ensure our results are reproducible by others. Subcompartments were called at the following resolutions: 10 kbp, 20 kbp, 50 kbp, 100 kbp, 200 kbp and 500kbp. However, only subcompartments at 10 kbp resolution were used for further data analyses. The same workflow was used to generate the draft figures for Fig. 1B and 1D as well as supplementary figures S16-19. Fig. 1C was generated using the output of the viz step of dcHiC as the starting point.

Figure S9-11 were generated by tracking subcompartment assignments across cell types. Bins were paired based on their genomic coordinates, then subcompartment switches were classified into 5 classes: neutral (no switch), reverted (e.g. switch occurred in T1 but was reverted in C1), transition to open/close and other switches (e.g. partial reversion: A3→A1→A2). Subcompartment switches were further classified using a delta score *Δ_ij_* defined as follows. We quantify the degree of switching between two stages, i and j (*Δ_ij_*), by assigning ranks to the subcompartments (B3=0…A3=7) and calculating the rank difference. A negative rank difference (*Δ_ij_*) thus indicates a transition towards B-type subcompartments. Conversely, a positive value signifies a shift towards A-type compartments.

Data analysis steps are defined in workflow compartment_analysis which was run by executing script run_compartment_analysis.sh.

### Differential expression analyses

Differential expression analysis was performed using DESeq2 v1.38.0 and apeglm v1.20.0 (Love et al. 2014; Zhu et al. 2019).

DESeq2 was called with default parameters using the raw count table produced by nf-core/rnaseq as input. Log2-fold-change shrinkage estimation was computed with the lfcShrink function from DESeq2 using apeglm as shrinkage estimator and the following log2FoldChange cutoffs: 0.0, 0.1, 0.25, 0.5, 1.0, 1.5, 2.0, 2.5, 3.0, 3.5, 4.0, 4.5, 5.0.

The analysis was automated using workflow diff_expression_analysis.nf, which was launched with run_diff_expression_analysis.sh.

Supplementary figure S28 was generated by running script run_cluster_profiler_do.py. The script uses clusterProfiler v4.8.1 (Wu et al. 2021; Yu et al. 2012) and the disease ontology (DO) database from the DOSE v3.26.1 package (Yu et al. 2015) to perform over-representation analysis of DO terms (Schriml et al. 2023). Package enrichplot v1.20.0 (Wu et al. 2021) was used for visualization. The IDs of differentially expressed genes were provided as input to clusterProfiler. Genes were considered as differentially expressed based if they exhibit an absolute log2FoldChange value of 0.5 or greater and a p-value of 0.01 or smaller (correcting for multiple testing). Furthermore, clusterProfiler was run using a q-value of 0.05.

### Comparative analyses

All comparative analyses were performed using workflow comparative_analysis.nf.

Fig. S15 was generated by overlapping subcompartment labels with several different epigenetic markers. First, each ChIP-seq peak was assigned a score by summing the raw ChIP-seq signal over each peak, then each peak was assigned to a subcompartment. Finally peaks were grouped by subcompartment label and the mean signal was computed for each subcompartment.

Fig. 2A and supplementary Fig. S29 were generated by overlapping expression levels in TPMs with subcompartment labels overlapping genes from Gencode v43 for hg38. In case a gene was tagged with multiple subcompartment labels, the gene was assigned the label with the largest coverage. In case of coverage tie, the gene was discarded (note that this is an extremely rare occurrence).

Figure 2B was generated using pyGenomeTracks (Ramírez et al. 2018; Lopez-Delisle et al. 2021) to visualize up/down regulated genes overlapping a region involved in subcompartment switching.

Figures 2C-F were generated by overlapping subcompartments with differentially expressed genes across all three cell types. First genes were assigned a subcompartment label by overlapping their TSS with the subcompartment annotation generated by dcHiC. Next genes were grouped based on log2FoldChange and p-value in down-regulated, non-differentially expressed and up-regulated genes (lfc=2.0; pvalue=0.01). Finally each group of genes was plotted as heatmaps showing subcompartment switches across pairs of cell types (genes not involved in subcompartment switches are not shown). Figures 2D and 2F were generated from the table underlying heatmaps 2C and 2E as follows: compute the sum of genes for each diagonal *i*; starting from diagonal *i* = *1*, pair diagonal i with diagonal −*i*; finally compute the log2 ratio between the sum of diagonal *i* and the sum of diagonal −*i*. Positive values indicate a positive correlation between one of the classes of differentially expressed genes and switches towards open chromatin while negative values indicate a positive correlation with subcompartment switches towards close chromatin.

The draft figure for Fig. 3A was generated with bin/plotting/plot_tad_cliques.py using TAD cliques, TADs, compartment PCA and gencode v43 gene annotation as input.

The draft figure for supplementary Fig. S32 was generated by overlapping subcompartment states with TADs annotated with the size of the largest clique to which they belonged to.

The draft figures for Fig. 3F-G and supplementary Fig. S40 were generated by clustering TADs and TAD cliques using HDBSCAN (McInnes et al. 2017; McInnes and Healy 2017) based on their subcompartment composition.

First, domains (that is TADs or TAD cliques) were annotated with their subcompartment state composition by overlapping subcompartment states with the domains using annotate_domains_with_subcompartments.py. This resulted in a count matrix with one row per domain, where each row counts the number of bins labeled for each subcompartment state. For example, given a domain overlapping 3 A1 bins, 10 A2 bins and 2 A3 bins, the entry in the count matrix corresponding to this domain would be 0, 0, 0, 0, 0, 3, 10, 2.

Next, this count matrix was given as input to cluster_domains_by_subcompartment_state.py, which clustered domains using HDBSCAN.flat based on their similarity in subcompartment composition. Clustering was performed using the following settings: n_clusters=9; metric=euclidean; min_cluster_size=200; min_samples=5; cluster_selection_method=leaf. Finally, clusters were visualized using plot_domain_subcompartment_clusters.py.

All comparative analyses are defined in workflow comparative_analysis.nf which was launched with script run_comparative_analysis.sh.

### 3D genome modeling

A patched version of Chrom3D v1.0.2 was used to generate 3D genome models for 10A, T1 and C1 (Paulsen et al. 2017). Briefly, this Nextflow workflow runs NCHG (doi.org/10.5281/zenodo.12680450) to identify statistically significant cis and trans interactions. Translocated regions were excluded when identifying statistically significant trans interactions. For intra-chromosomal interactions we used a log-ratio of 1.2 and adjusted-pvalue of 0.01 as cutoffs, while for inter-chromosomal interactions a log-ratio of 0.25 and adjusted-pvalue of 0.01 were used as cutoffs. The workflow then uses statistically significant interactions in Chrom3D simulations as spatially proximal interaction constraints. Furthermore, the workflow matches LADs to the corresponding TAD domains such that these domains can be used as peripheral sub-nuclear constraints in the simulations. A total of 100 Chrom3D simulations were run for each condition with a nuclear occupancy of 0.15, at TAD resolution and 2 million simulation steps (Paulsen et al. 2017).

ChimeraX was used to visualize the simulated Chrom3D models (Pettersen et al. 2021). The median distance of each chromosome from the nuclear center was calculated for each Chrom3D simulation and visualized using ggplot2 (Villanueva and Chen 2019). The median distance and the standard deviation of each sub-compartment were likewise evaluated for each Chrom3D simulation and visualized using ggplot2. Sub-compartment distances from the nuclear center were statistically evaluated and compared within conditions using the Wilcoxon rank sum test in R with the function wilcox.test.

### FISH analysis

Analysis of FISH data is automated using workflow fish.nf. Draft plots have been generated using notebooks plot_fish_blob_distance_stats.ipynb and plot_fish_blob_radial_positioning.ipynb.

The FISH data analysis consists of three steps:

- Nuclei segmentation
- Probe localization
- Plot generation and statistical tests

#### Nuclei segmentation

Nuclei segmentation is performed using DAPI signal (i.e. the blue channel of the RGB image produced by the microscope).

First, RGB images containing one or more nuclei are converted to grayscale by extracting the blue channel from the images. Grayscale images are then sharpened using unsharp masking. This is done to improve the separation of the profiles of nuclei that are located very close to one another. Next, foreground objects (i.e. nuclei profiles) are highlighted using Otsu thresholding (Otsu 1979). Objects that are significantly smaller than the plausible nuclei profiles are masked out. Next, we compute the contour of each individual object using the “marching squares” method. Finally, object contours are used to crop out individual nuclei.

To properly deal with various edge cases (such as mitotic nuclei and overlapping nuclei), nuclei whose contour significantly deviates from an ellipsoid are flagged so that they can be skipped in subsequent analysis steps. This is achieved by first fitting an ellipse to the contour detected using the first algorithm described in (Fitzgibbon and Fisher). Next, we compute the overlap coefficient (OC) between the fitted ellipse and the nucleus contour, and mask nuclei with an OC of 0.95 or lower. Finally, nuclei that are only partially visible in an image are also flagged.

#### Probe localization

For each nucleus, we extract the red and green channels from the RGB image produced by the segmentation step. The resulting grayscale images are processed independently as follows.

Given a probe and a cell line, we can estimate the number of probe signals we expect to find in each nucleus. This information is used to guide the blob (i.e. probe signal) detection using the Laplacian of Gaussian (LoG) method. We perform a sweep of LoG parameters, starting from parameters that are suitable to detect large blobs in an image. If the first run of the LoG algorithm detects *N* or more blobs, where *N* is the number of expected probe signals, then the parameter sweep is halted. Otherwise, the coordinates of detected blobs (if any) are stored, and the LoG algorithm is re-run using parameters to detect progressively smaller blobs. Every iteration, the coordinates of newly detected blobs are merged with the coordinates of blobs detected in previous runs of the algorithm. The parameter sweep stops as soon as *N* or more blobs are detected, or when the parameter space is exhausted.

#### Plot generation and statistical tests

The coordinates of blobs identified in the previous step are finally used to generate Fig. 4F and Supplementary Fig. S46.

The number of nuclei and observations for each condition is shown in Suppl. Table S18.

### Software used throughout data analysis code

The following software packages were used throughout the code used for data analysis:

● bedtools (Quinlan and Hall 2010) - to perform common set operations on genomic intervals.
● bedGraphToBigWig (Kent et al. 2010) - to convert bedgraph files to bigWig format.
● bioframe (Open2C et al. 2022) - to perform common set operations on genomic intervals.
● hictkpy (Rossini and Paulsen 2024) - to perform certain input-output operations on .cool files.
● matplotlib (Hunter May-June 2007) - to generate plots.
● NumPy (Harris et al. 2020) - to efficiently perform arithmetic operations on vector data.
● OpenCV - to manipulate and process microscopy images.
● pandas (McKinney 2010) - to read, write and manipulate tabular data using dataframes.
● pyBigWig - to read and write bigWig files.
● samtools (Danecek et al. 2021) - to index FASTA files and perform IO operations on SAM, BAM and CRAM files.
● scikit-image (van der Walt et al. 2014) - to manipulate and process microscopy images.
● Scipy (Virtanen et al. 2020) - to compute Pearson correlation values and perform statistical tests.
● Seaborn (Waskom 2021) - to generate plots.

### Software patches

This section describes the purpose of the patches we applied to Chrom3D, cooler, dcHiC, and HiNT. Patch files are available under containers/patches.

Patch for nf-core/hic v2.0.0 (https://zenodo.org/records/2669513)

Our patch involved updating the pipeline to correctly handle the --restriction_site and -- ligation_site parameters.

Our patch was accepted by upstream and is now part of v2.1.0 of the pipeline. See nf-core/hic/pull/153 for more details.

Patches for dcHiC d4eb244 (Chakraborty et al. 2022)

We patched dcHiC as follows:

● Define a default seed and introduce a --seed CLI option to ensure results are reproducible across runs
● Wrap calls to depmixS4::fit into a try-catch block to attempt model fitting up to 5 times in case of spurious failures
● Properly handle the possibility that a chromosome does not have any bin overlapping one or more subcompartment type.

Our patches were accepted by upstream but are not yet part of a stable release. See ay-lab/dcHiC/pull/59, ay-lab/dcHiC/pull/60, ay-lab/dcHiC/pull/62 and containers/patches/dchic.patch for more details.

Patch for cooler v0.9.1 (Abdennur and Mirny 2020)

We patched cooler to ensure that convergence of cooler balance using cis-only interactions was correctly reported for all chromosomes instead of just the last one.

When balancing interactions using the --cis-only flag, cooler balances interactions for each chromosome independently. Given that there is no guarantee that ICE will converge within the given number of iterations, cooler balance stores an attribute inside balanced cooler files to report whether balancing was successful (i.e. convergence was achieved).

In cooler v0.9.1 there is a bug in the logic that writes this attribute that causes the convergence for the last chromosome balanced to be reported instead of the convergence status for each chromosome. Our patch addresses this bug such that the convergence status of matrices balanced with --cis-only can be assessed correctly.

The patch also addressed some minor issues related to detecting pandas’ version at runtime and handling of bin sizes represented using unsigned integers.

Our patches were accepted by upstream and are now part of cooler v0.9.2.

See open2c/cooler/pull/313, open2c/cooler/pull/323, open2c/cooler/pull/324 and containers/patches/cooler.patch for more details.

Patch for Chrom3D v1.0.2 (Paulsen et al. 2017)

We patched Chrom3D to support building the project using CMake and addressing a few compiler warnings and errors that were preventing us from compiling Chrom3D with a modern C++ compiler toolchain. For more details, refer to the following two patch files available at robomics/chrom3d-nf (doi.org/10.5281/zenodo.12687842) chrom3d-cmake.patch, chrom3d-fix-warnings.patch.

Patch for HiNT v2.2.8 (Wang et al. 2020)

We patched HiNT to support processing Hi-C matrices using Arima restriction enzymes. See containers/patches/hint.patch for more details.

## Data Access

All raw and processed sequencing data generated in this study have been submitted to the NCBI Gene Expression Omnibus (GEO; https://www.ncbi.nlm.nih.gov/geo/) under the following accession numbers: GSE246689, GSE246599, GSE247171, and GSE246947.

All raw and processed microscopy data generated in this study have been submitted to Zenodo (https://zenodo.org/) and are available for download at the following DOI: 10.5281/zenodo.13255558.

## Supporting information

Supplemental Table with SCC correlation values

## Acknowledgments

Molecular graphics and analyses performed with UCSF ChimeraX, developed by the Resource for Biocomputing, Visualization, and Informatics at the University of California, San Francisco, with support from National Institutes of Health R01-GM129325 and the Office of Cyber Infrastructure and Computational Biology, National Institute of Allergy and Infectious Diseases. HiC and RNA-Seq library constructions and Illumina sequencing were done at the Biomolecular Resource Facility (BRF), JCSMR, Australian National University. Some of the analyses were performed on resources provided by Sigma2 - the National Infrastructure for High Performance Computing and Data Storage in Norway, with account number NN8041K. We thank Anita L. Sørensen for performing FISH experiments and microscopy imaging. This work was funded by the Norwegian Research Council project. no. 324137 (JP), the Norwegian Cancer Society (PC), UNIFOR (PC and AB), and National Health and Medical Research Council ID#1182759 (DT)

## Authors’ contributions

JP, PC, DT conceived and designed the study; JP, RR and MO designed data analyses; RR and MO analyzed data; RR processed Hi-C, RNA-seq, ChIP-seq, and FISH data and did downstream analyses; MO performed 3D genome modeling; MN did Hi-C experiments; RD and YD did cell culture work and RNA-seq; AB and MA analyzed LADs; JP, PC, DT supervised the work. All authors read and approved the final manuscript.

