## Supplemental Figures for "Loss of multi-level 3D genome organization during breast cancer progression"

### Supplementary figures

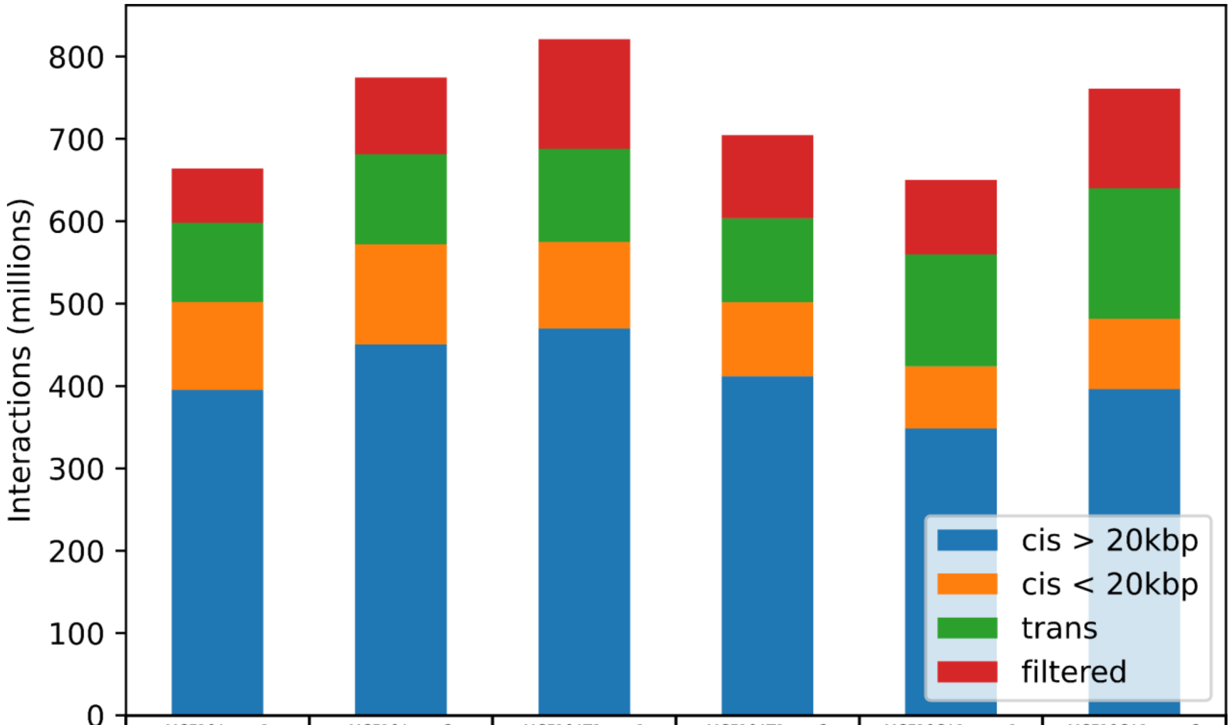


| | MCF10A rep.1 | MCF10A rep.2 | MCF10AT1 rep.1 | MCF10AT1 rep.2 | MCF10CA1a rep.1 | MCF10CA1a rep.2 |
|---|---|---|---|---|---|---|
| cis > 20kbp | 395.19 | 450.37 | 469.72 | 411.8 | 348.31 | 396.13 |
| cis < 20kbp | 106.91 | 121.64 | 105.22 | 89.85 | 75.64 | 85.27 |
| trans | 95.93 | 109.5 | 112.73 | 101.96 | 135.7 | 158.56 |
| filtered | 66.14 | 92.97 | 133.61 | 101.12 | 90.5 | 120.95 |
| total | 664.17 | 774.48 | 821.28 | 704.73 | 650.15 | 760.91 |

**Fig. S1**: Barplot showing the number of Hi-C interactions (in millions) classified by type: long-range cis (blue), short-range cis (orange), trans (green) and filtered out (red). The table under the graph shows the data underlying the barplot.

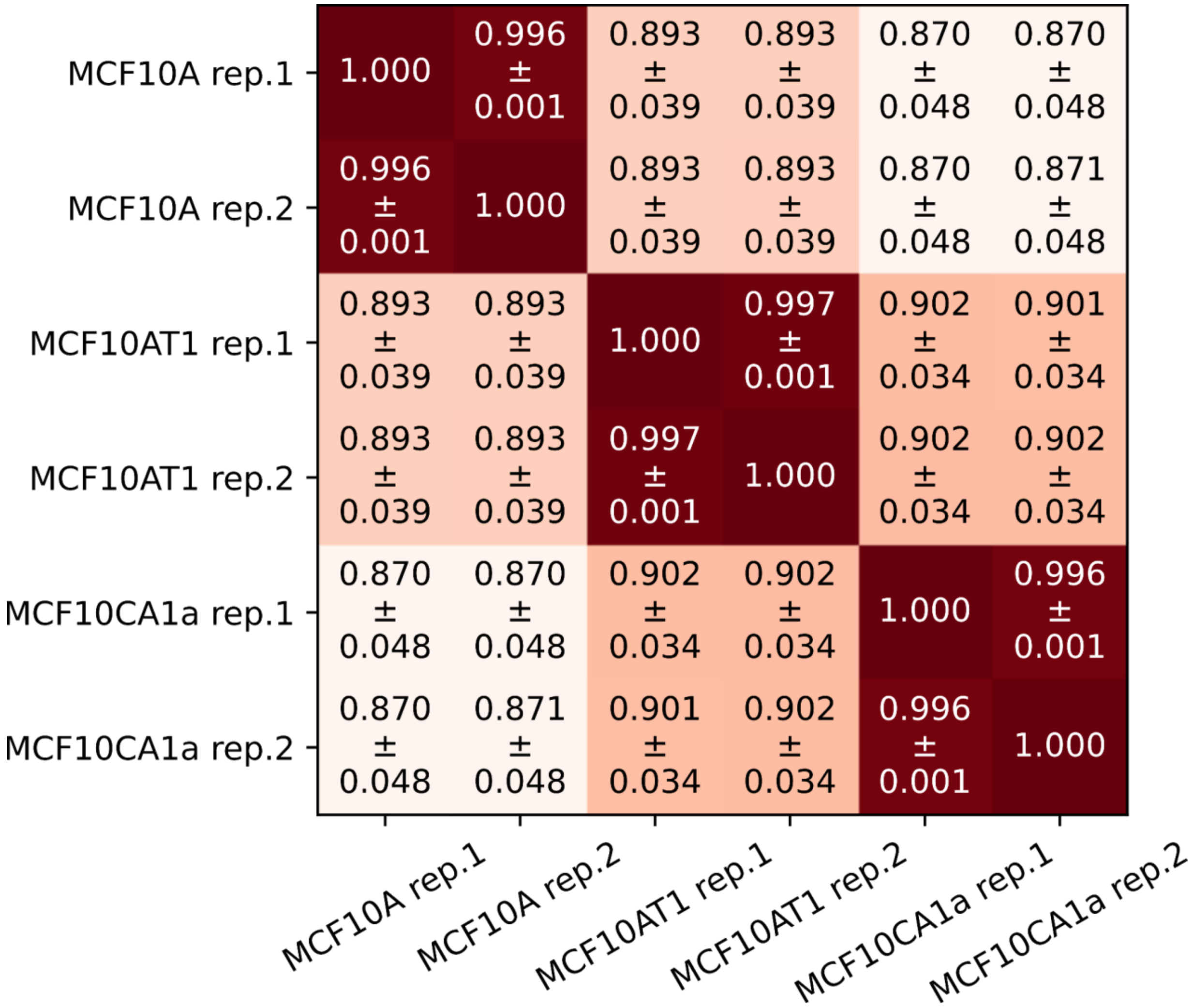


**Fig. S2:** Heatmap showing the weighted average of the stratified correlation coefficient (SCC) computed for each pair of samples using HiCRep, which avoids inflated correlations from distance-dependent decay in HiC data. Chromosome sizes are used as weights in the computation of the SCC weighted average. Overlaid numbers show the weighted average and standard deviation for each pair of samples.

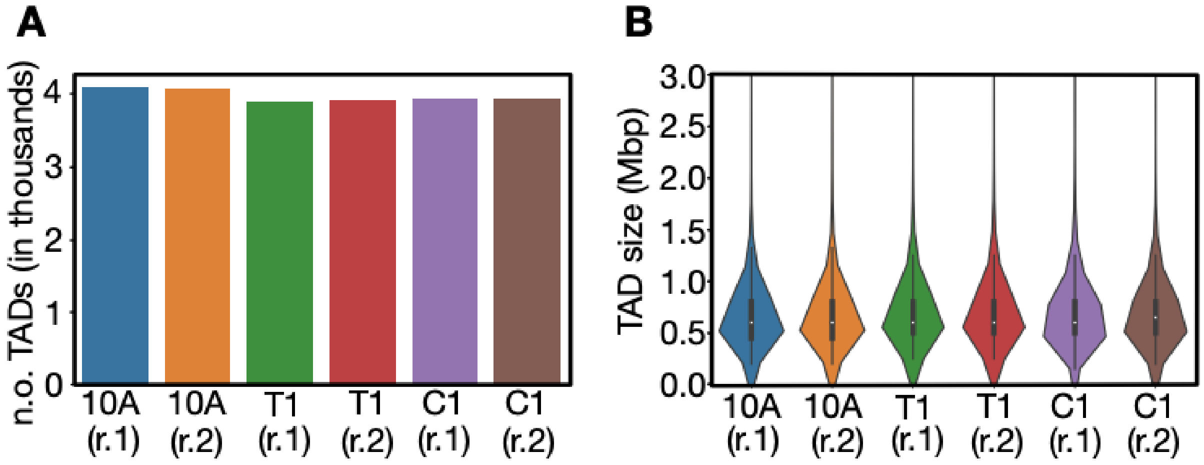


**Fig. S3:** **A**: Number of TADs (in thousands) for each of the six Hi-C samples generated. **B**: Comparison of genomic sizes of all TADs for the six samples.

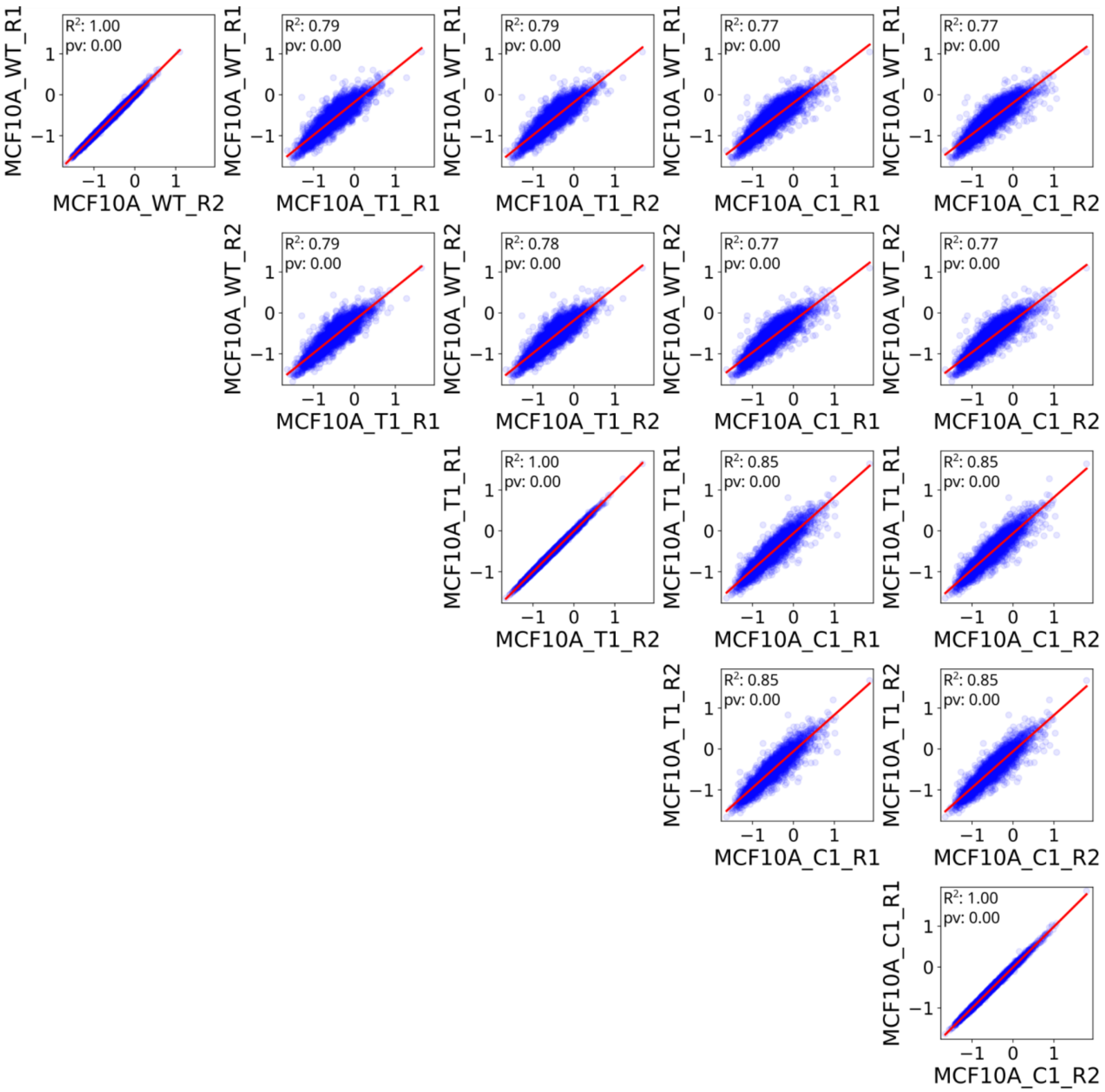


**Fig. S4**: Scatterplot contrasting TAD insulation scores at 50 kbp resolution for all possible pairs of samples. Trend line is depicted in red.

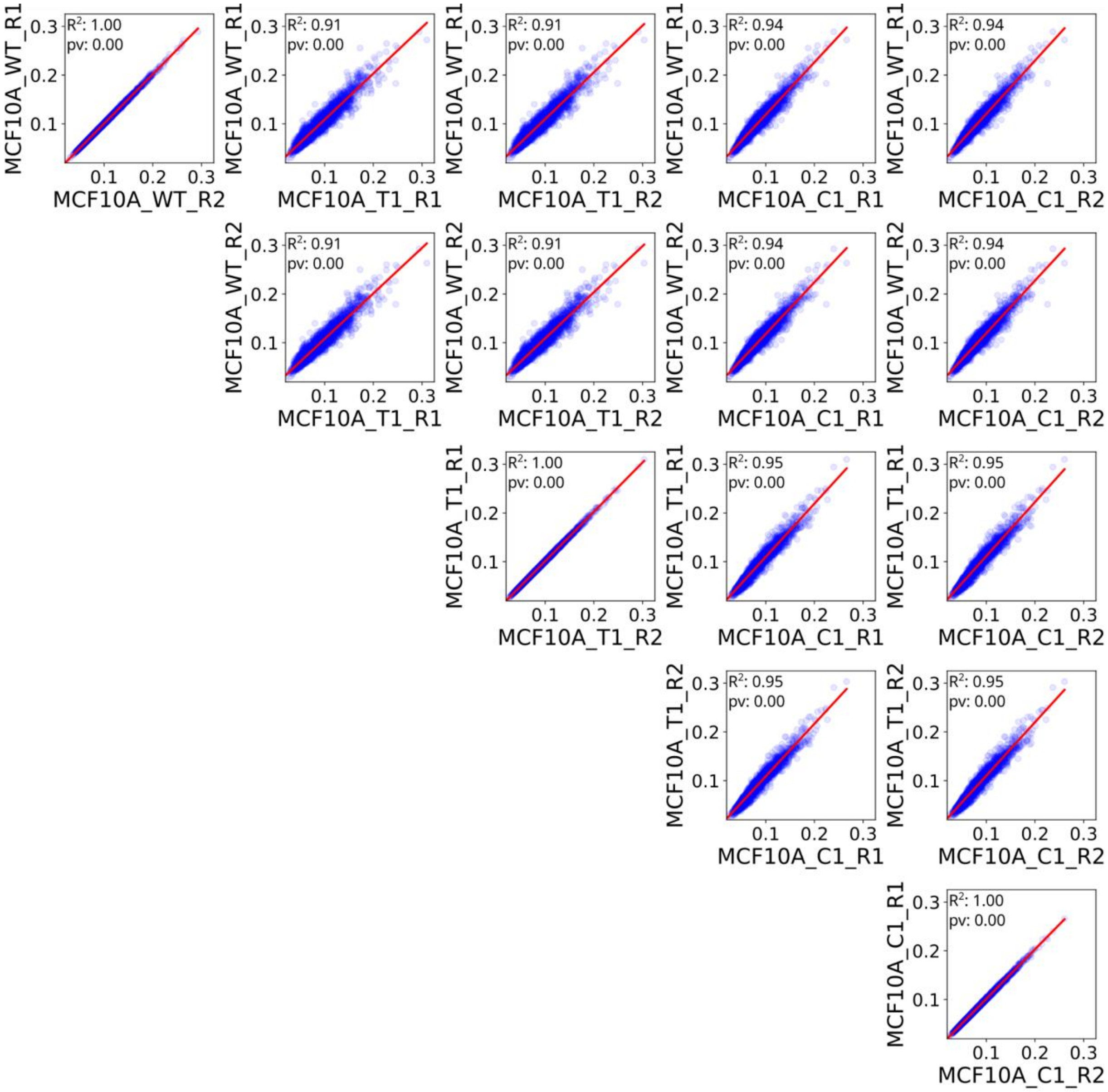


**Fig. S5**: Scatterplot contrasting the normalized number of interactions of MCF10A TADs at 50 kbp resolution across all possible pairs of samples. Trend line is shown in red. Scores are computed by aggregating interactions from the upper triangle of the Hi-C matrix overlapping TADs. Aggregated interactions are normalized based on the number of pixels belonging to a TAD.

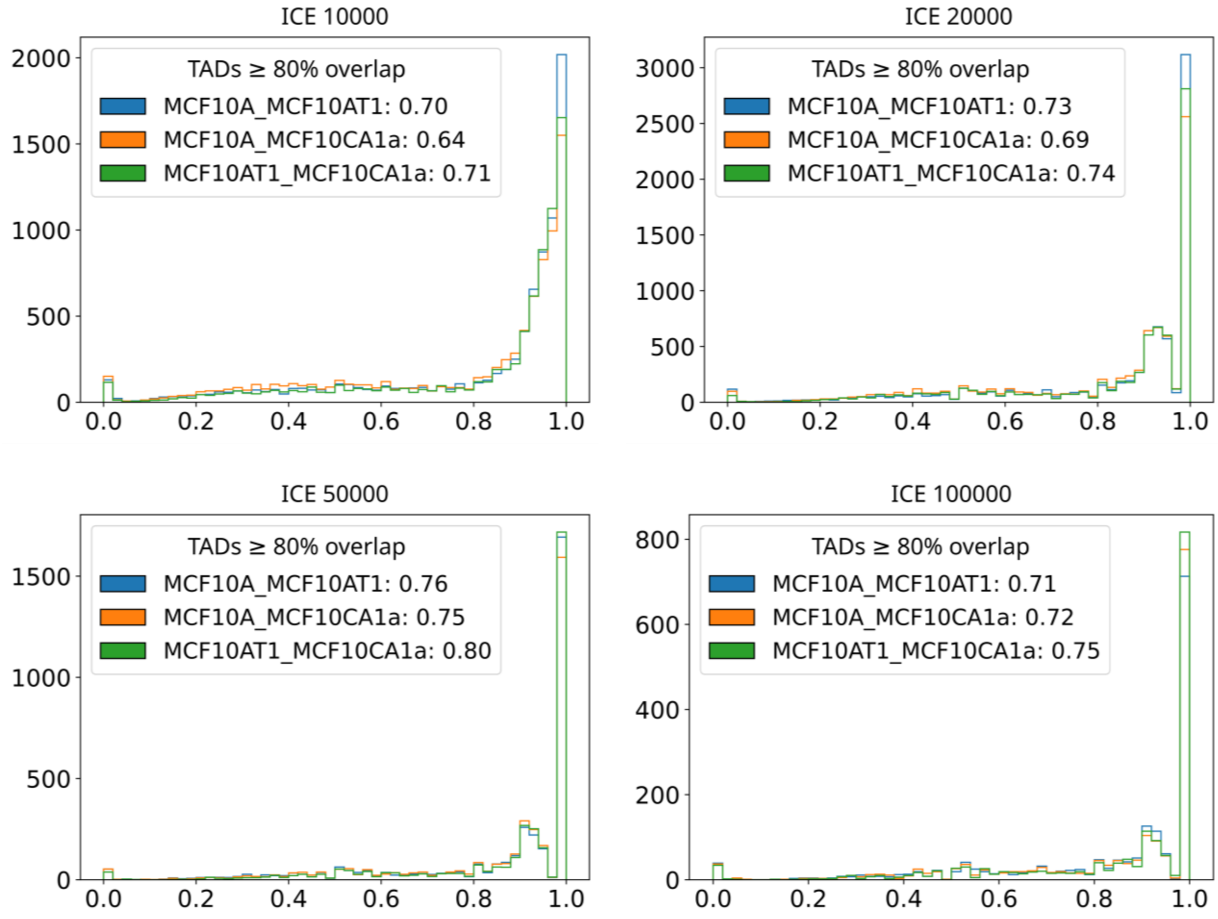


**Fig. S6**: Histogram of TAD Jaccard overlap coefficients. Coefficients are computed from Hi-C matrices after collapsing replicates. Numbers shown in plot legends show the fraction of TADs with at least 80% overlap. Panels show coefficients computed using TADs called at 10 kbp (top left), 20 kbp (top right), 50 kbp (bottom left) and 100 kbp (bottom right) resolutions.

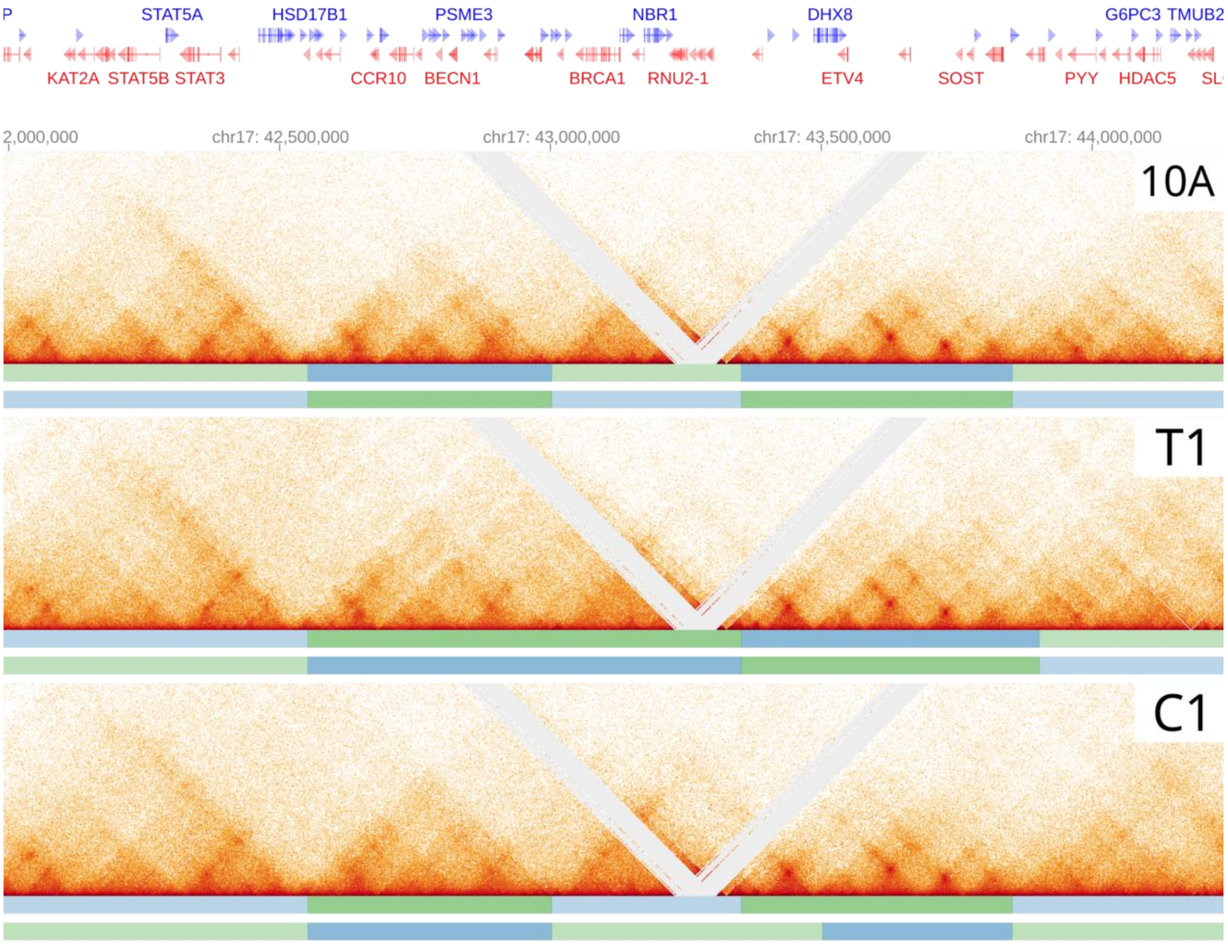


**Fig S7**: HiGlass view of the Hi-C matrices for 10A, T1, and C1 centered around the BRCA1 gene. Bars shown below the Hi-C matrices show the TADs called from repl. 1 and repl. 2 of the respective dataset.

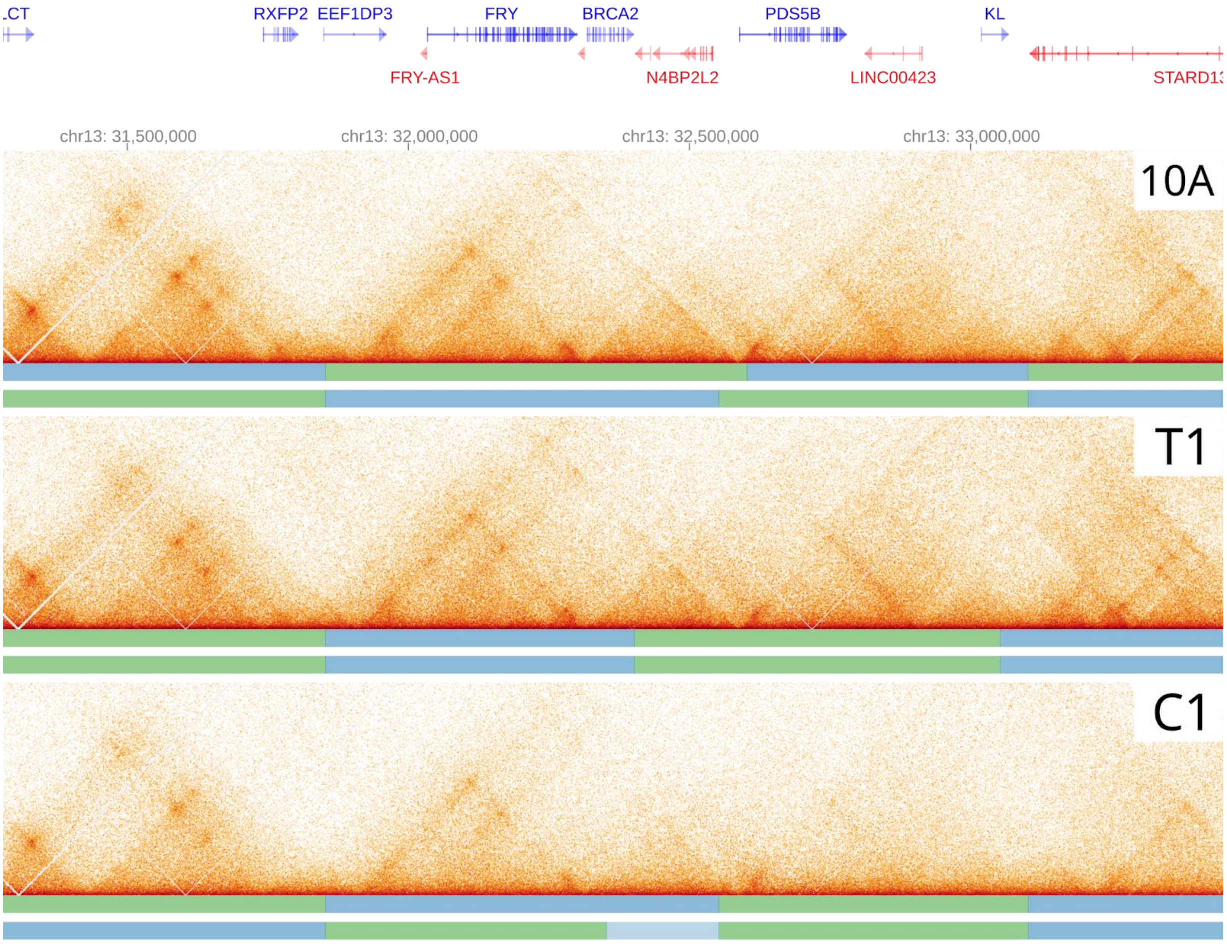


**Fig S8**: HiGlass view of the Hi-C matrices for 10A, T1, and C1 centered around the BRCA2 gene. Bars shown below the Hi-C matrices show the TADs called from repl. 1 and repl. 2 of the respective dataset.

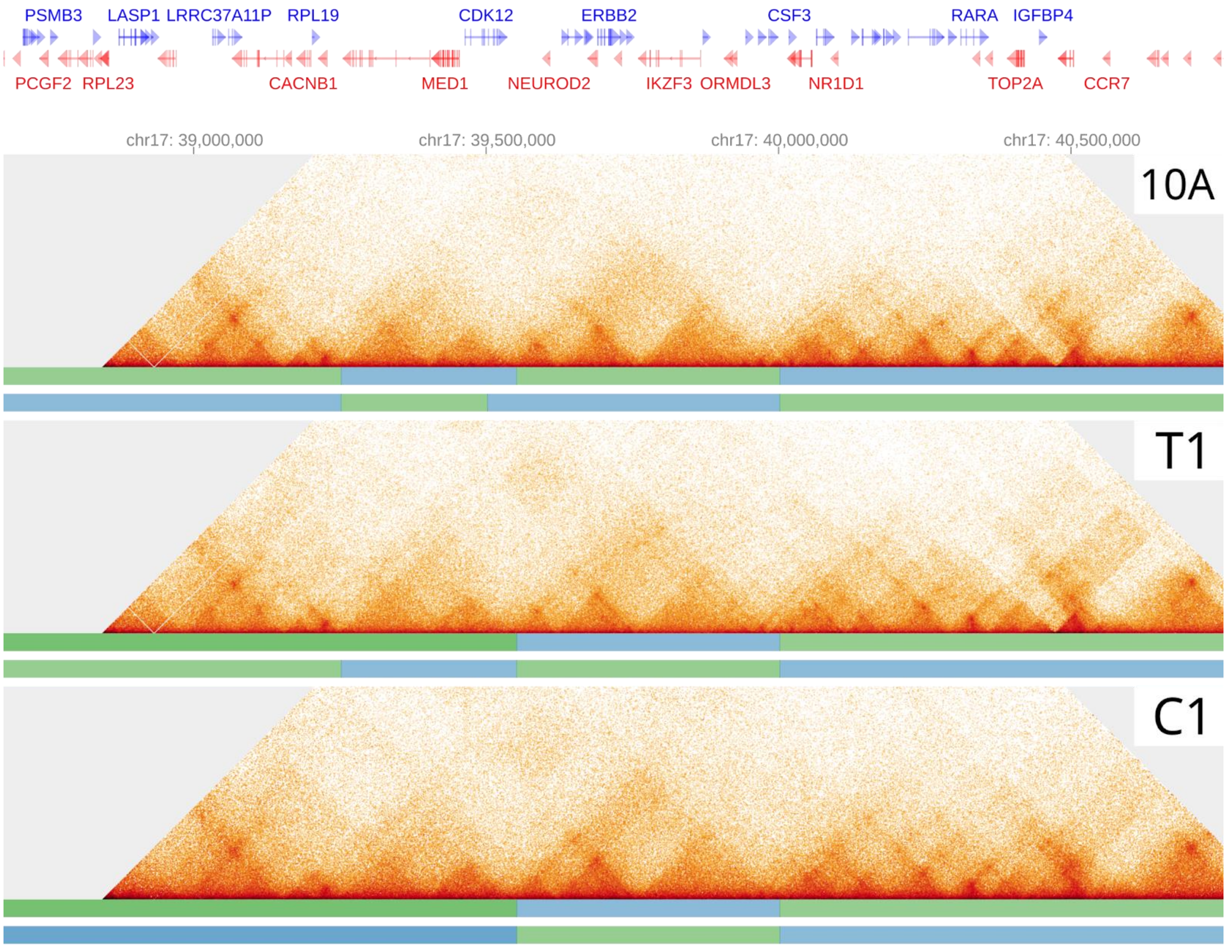


**Fig S9**: HiGlass view of the Hi-C matrices for 10A, T1, and C1 centered around the ERBB2 gene. Bars shown below the Hi-C matrices show the TADs called from repl. 1 and repl. 2 of the respective dataset.

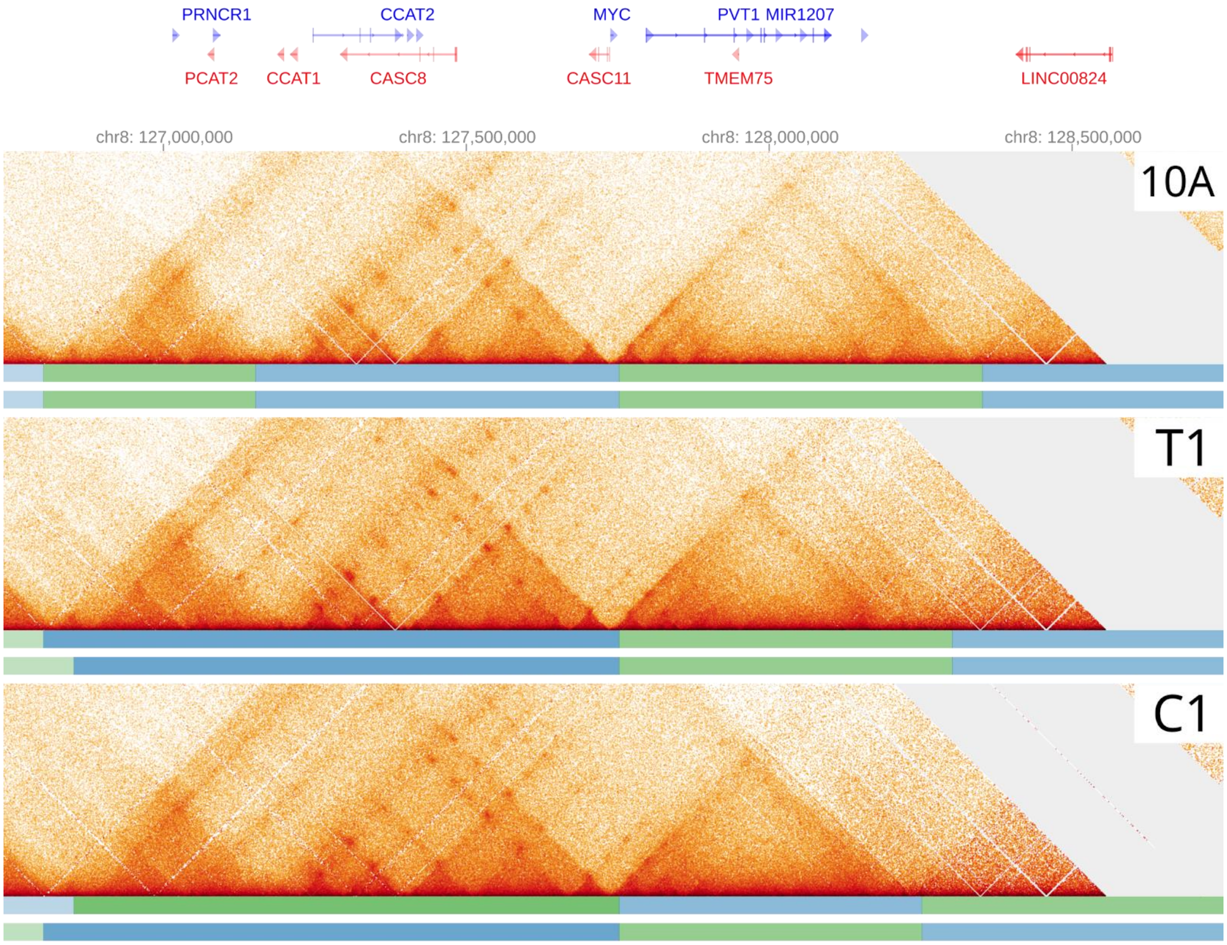


**Fig S10**: HiGlass view of the Hi-C matrices for 10A, T1, and C1 centered around the MYC gene. Bars shown below the Hi-C matrices show the TADs called from repl. 1 and repl. 2 of the respective dataset.

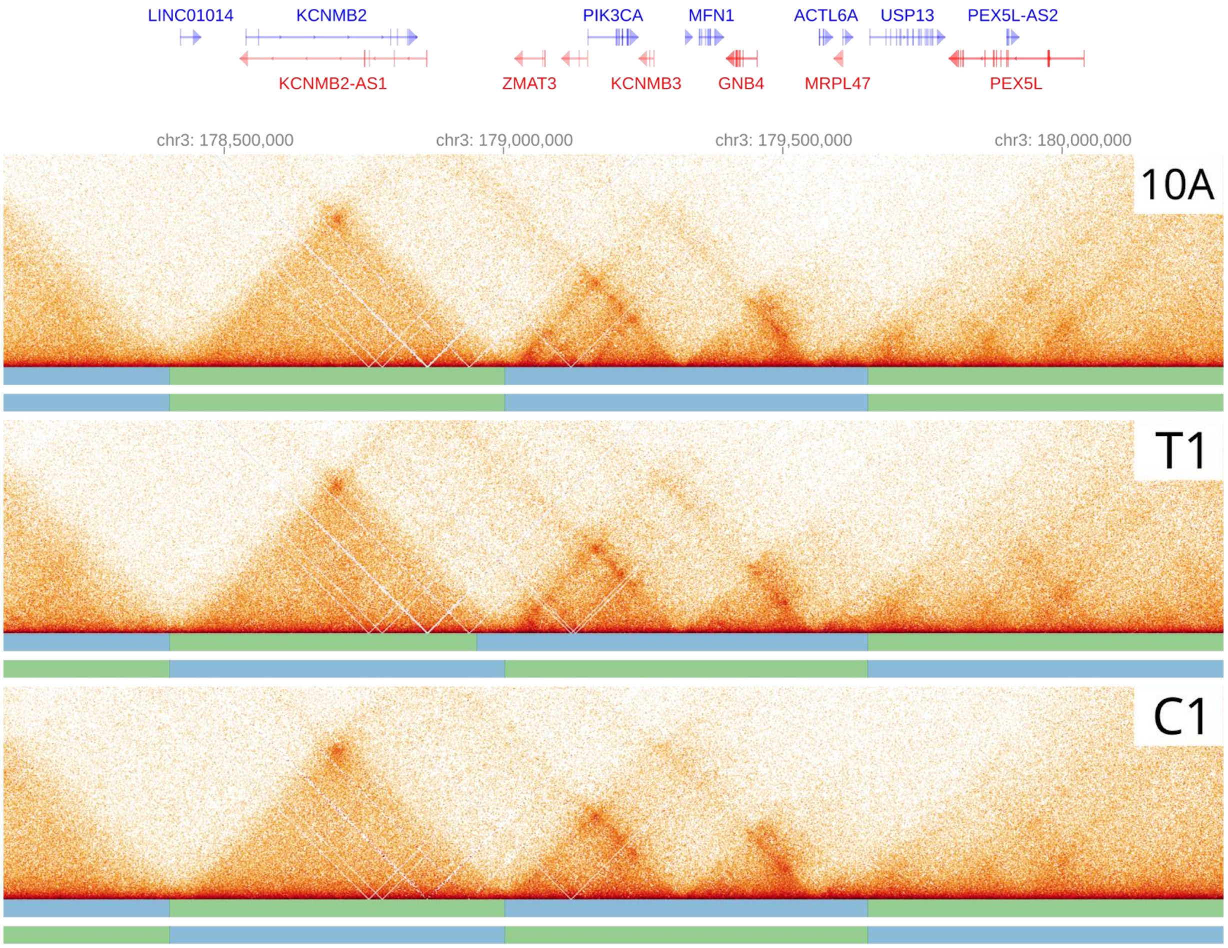


**Fig S11**: HiGlass view of the Hi-C matrices for 10A, T1, and C1 centered around the PIK3CA gene. Bars shown below the Hi-C matrices show the TADs called from repl. 1 and repl. 2 of the respective dataset.

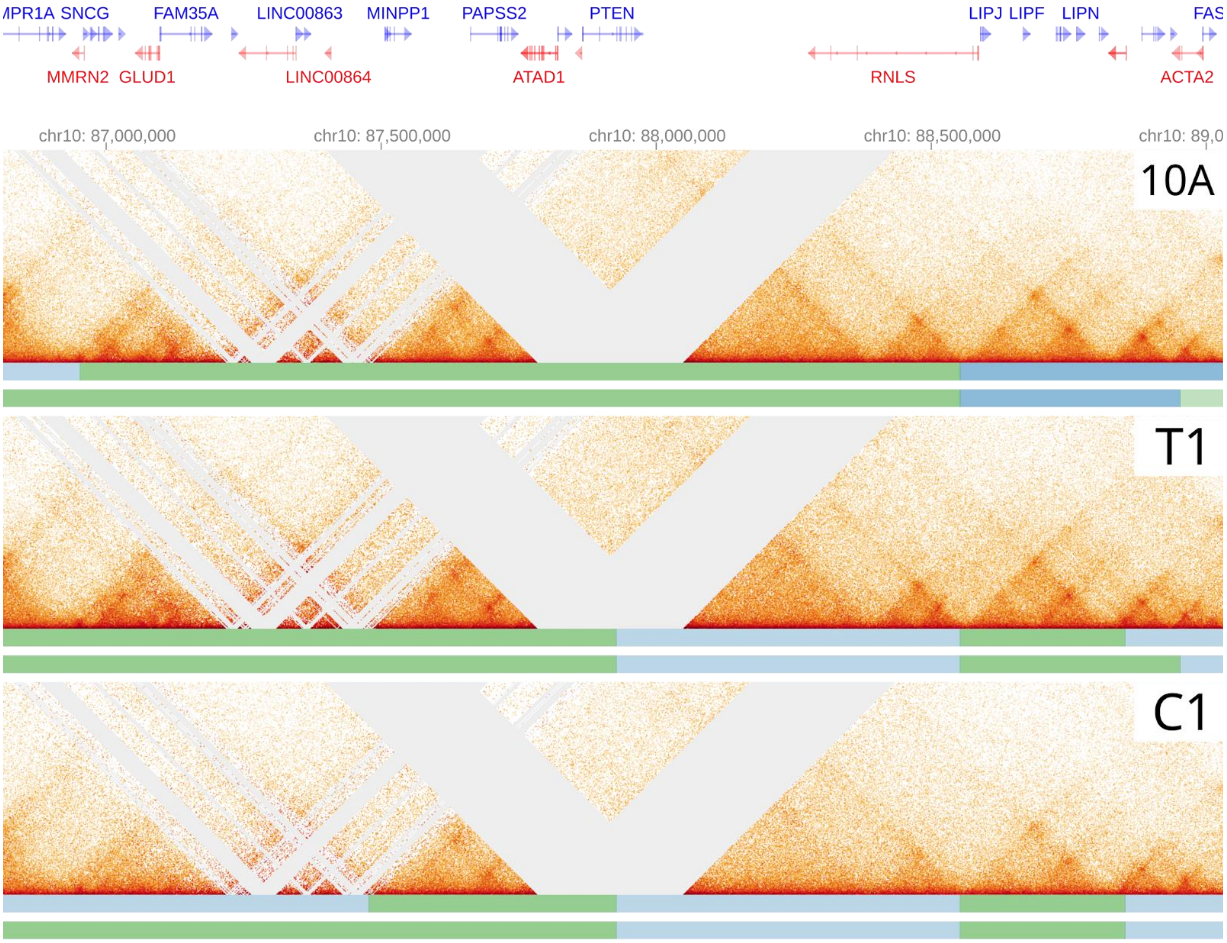


**Fig S12**: HiGlass view of the Hi-C matrices for 10A, T1, and C1 centered around the PTEN gene. Bars shown below the Hi-C matrices show the TADs called from repl. 1 and repl. 2 of the respective dataset.

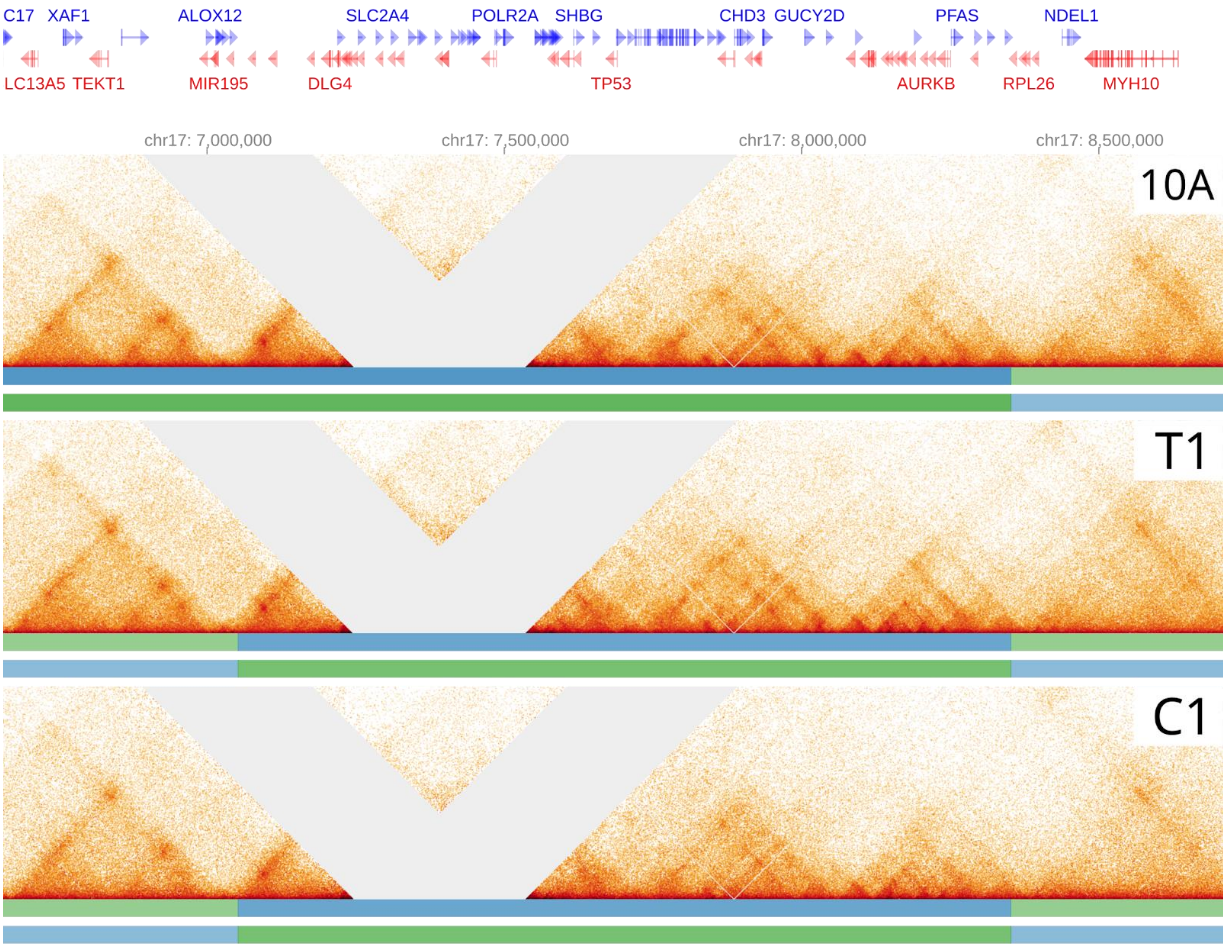


**Fig S13**: HiGlass view of the Hi-C matrices for 10A, T1, and C1 centered around the TP53 gene. Bars shown below the Hi-C matrices show the TADs called from repl. 1 and repl. 2 of the respective dataset.

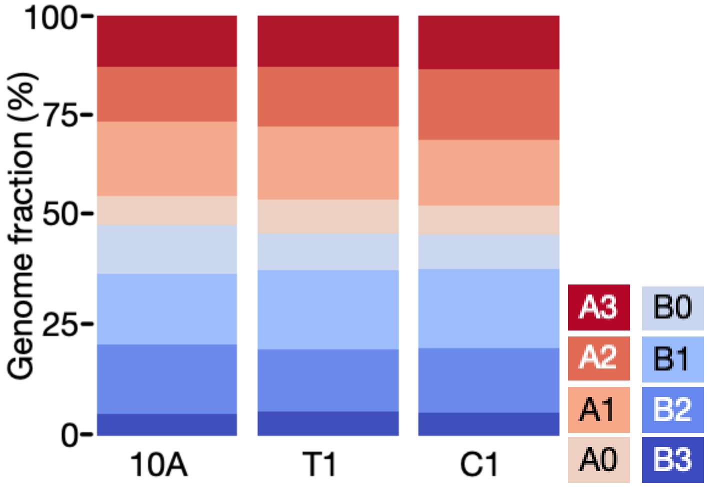


**Fig. S14**: Relative genome coverage of subcompartments (at 10kbp resolution) in the three samples.

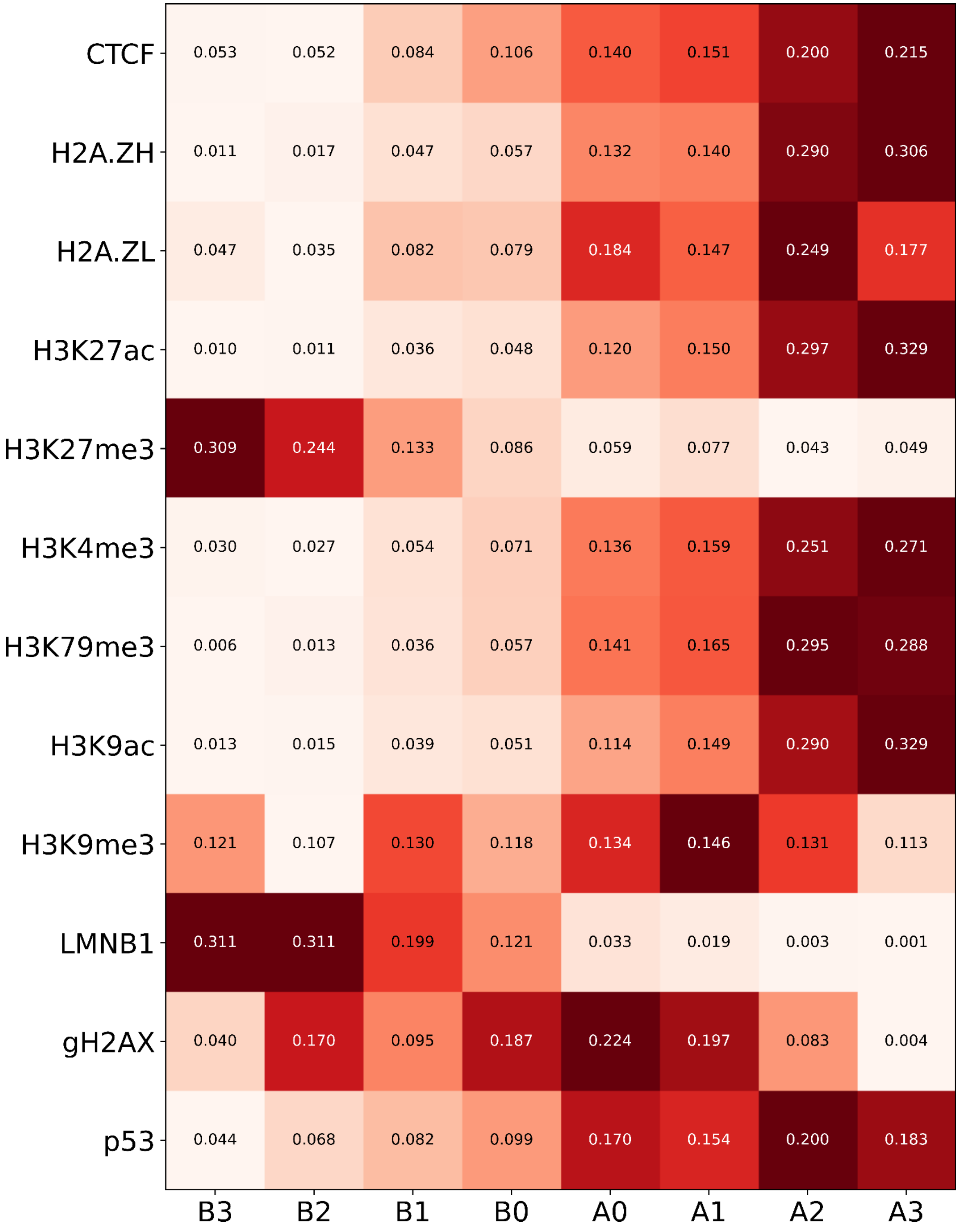


**Fig. S15**. Heatmap showing the average coverage of several epigenetic marks across subcompartments called on MCF10A matrices after collapsing replicates at 10 kbp resolution. Values overlaid on the heatmap show the average intensity of ChIP-seq signal over peaks overlapping each subcompartment. Color scale is normalized separately for each row to show the lowest and highest values of each row in white and dark red respectively.

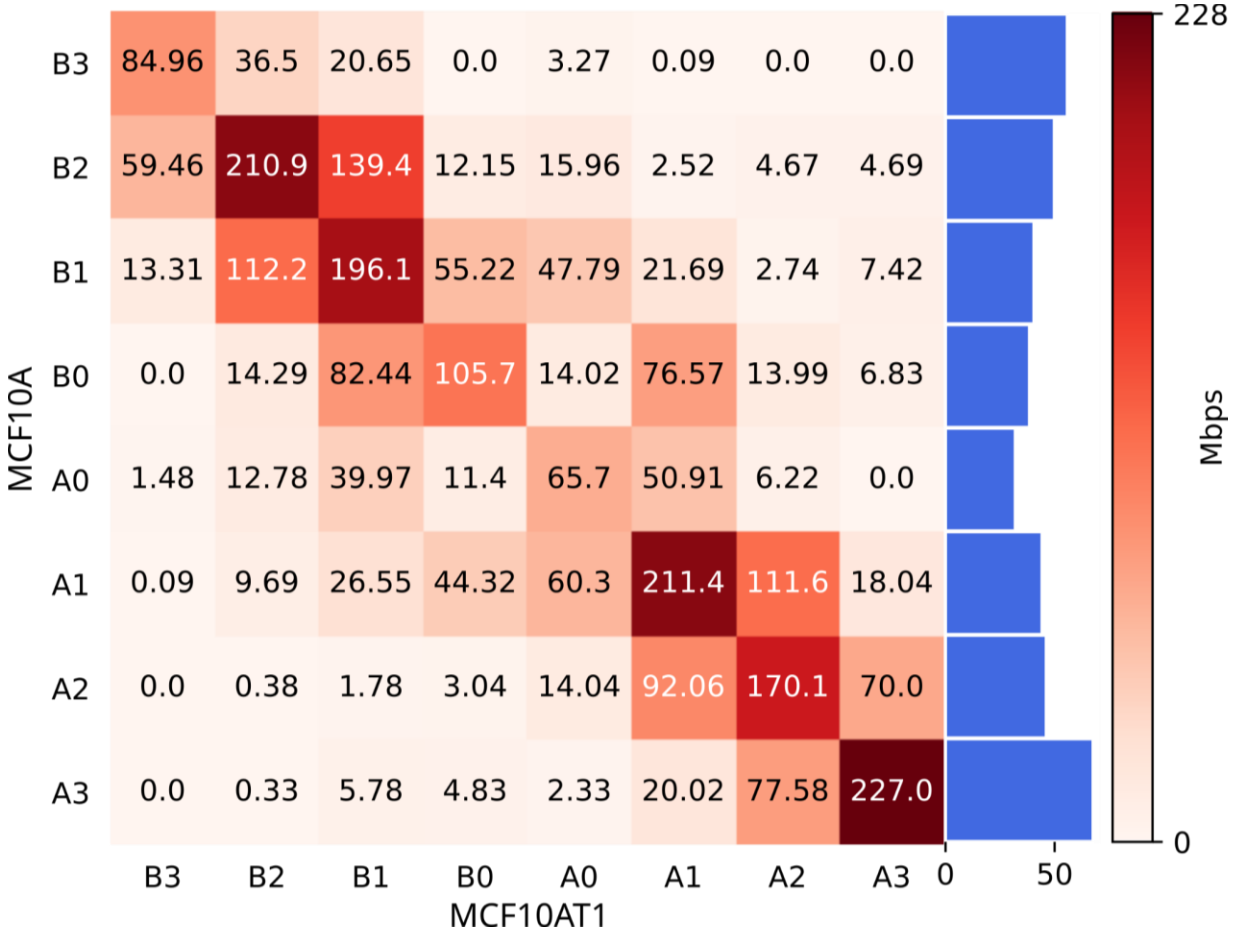


**Fig. S16:** Subcompartment switches (10kbp resolution) for MCF10A (WT) (vertical axis) vs MCF10AT1 (T1) (horizontal axis). Numbers are expressed in Mbp unless otherwise specified. Subcompartment switches are computed by comparing subcompartment labels across cell types. Comparison is done at the bin-level, comparing the same genomic regions.
The bar plot shows the fraction (%) of Mbps not involved in subcompartment switching. Bars are relative to the number of Mbps belonging to a given subcompartment in at least one cell type.

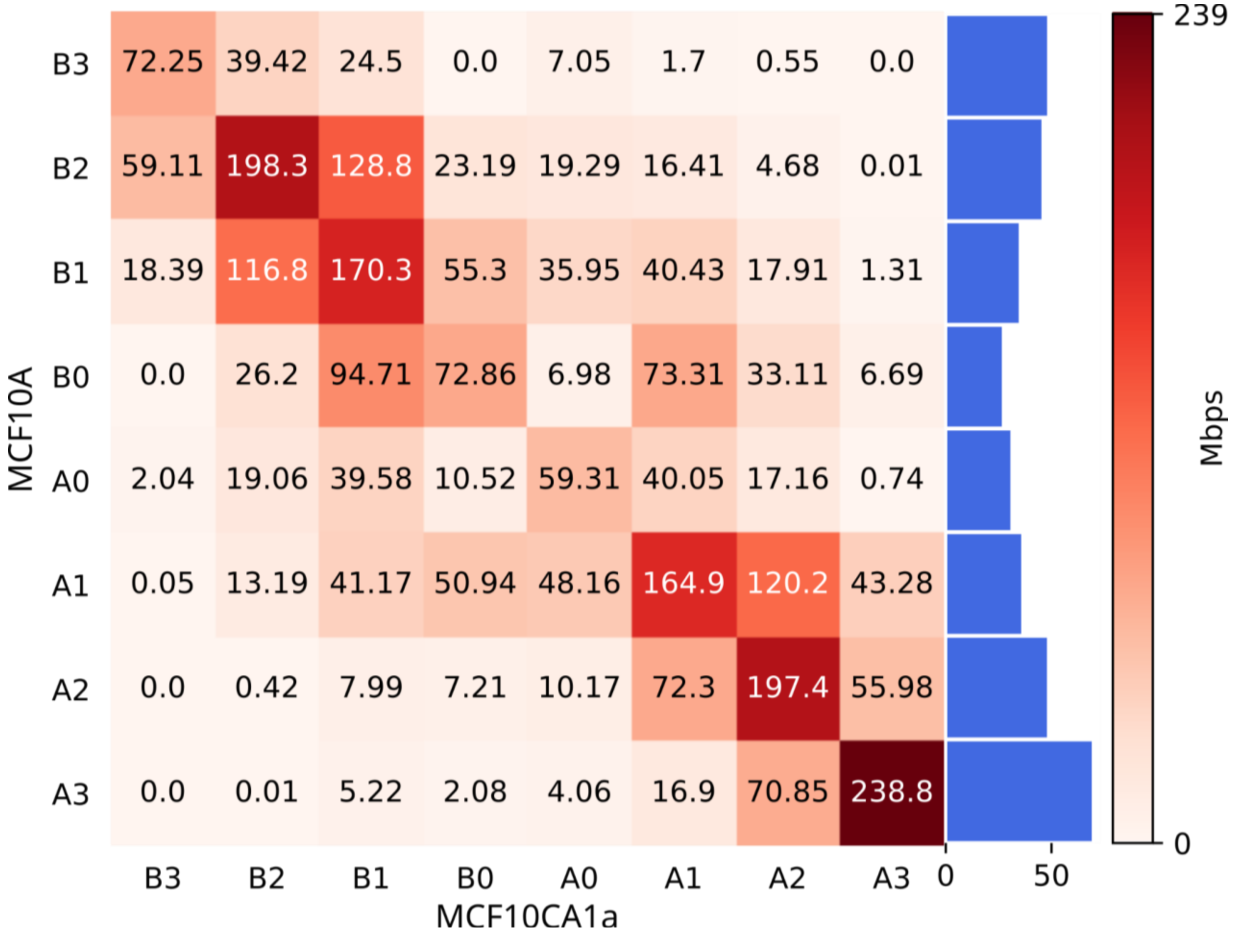


**Fig. S17:** Subcompartment switches (10kbp resolution) for MCF10A (WT) (vertical axis) vs MCF10CA1a (C1) (horizontal axis). Numbers are expressed in Mbp unless otherwise specified. Subcompartment switches are computed by comparing subcompartment labels across cell types. Comparison is done at the bin-level, comparing the same genomic regions. The bar plot shows the fraction (%) of Mbps not involved in subcompartment switching. Bars are relative to the number of Mbps belonging to a given subcompartment in at least one cell type.

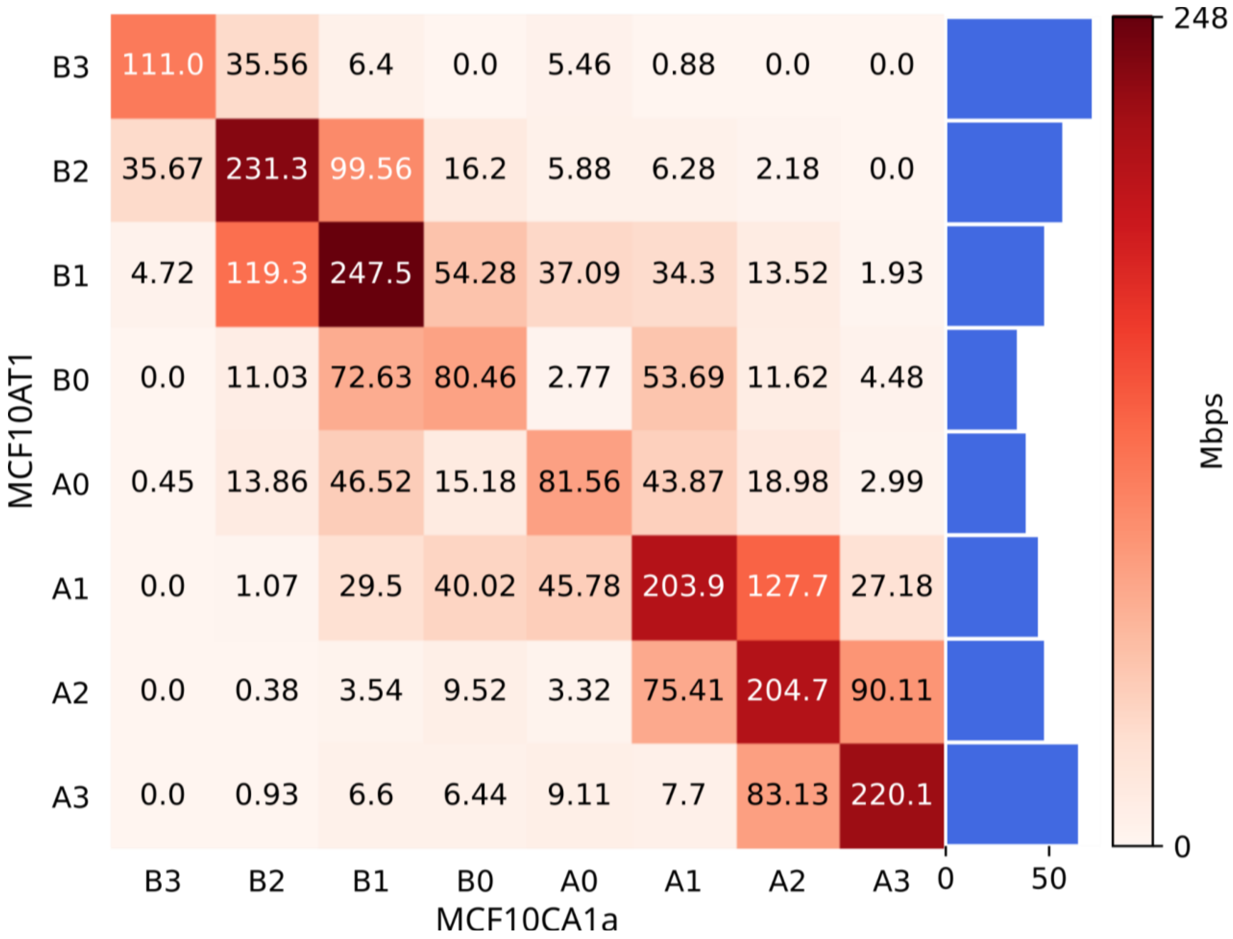


**Fig. S18:** Subcompartment switches (10kbp resolution) for MCF10AT1 (T1) (vertical axis) vs MCF10CA1a (C1) (horizontal axis). Numbers are expressed in Mbp unless otherwise specified. Subcompartment switches are computed by comparing subcompartment labels across cell types. Comparison is done at the bin-level, comparing the same genomic regions. The bar plot shows the fraction of Mbps not involved in subcompartment switching. Bars are relative to the number of Mbps belonging to a given subcompartment in at least one cell type.

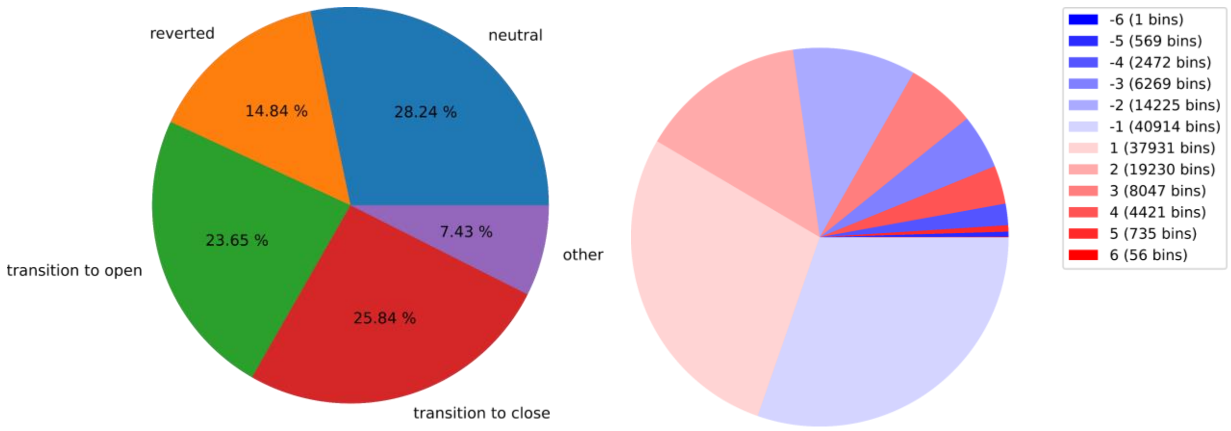


**Fig. S19**:
Left: Pie chart showing the fate of genomic regions involved in subcompartment switches from WT via T1 to C1. Right: Pie chart showing the magnitude and direction of subcompartment switching for genomic regions involved in subcompartment switches. Positive numbers indicate a switch towards A-like subcompartments while negative values represent a switch towards B-like subcompartments. Values represent the number of consecutive subcompartment steps that are switched (e.g. A3→A0 gives a value of -3).

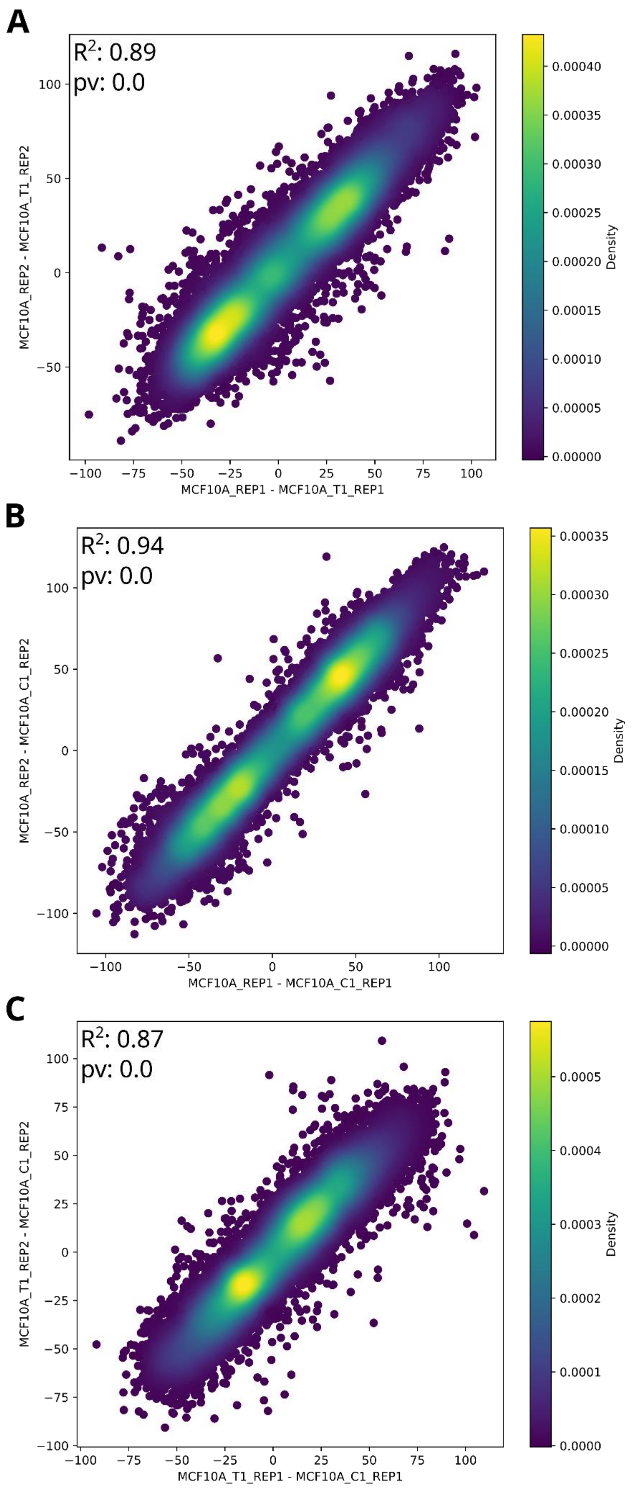


**Fig. S20**: Scatter plots comparing differences in compartment PCs. **A** Scatter plot comparing PC differences between 10A and T1 repl. 1 with PC differences between 10A and T1 repl. 2. **B** Scatter plot comparing PC differences between 10A and C1 repl. 1 with PC differences between 10A and C1 repl. 2. **C** Scatter plot comparing PC differences between T1 and C1 repl. 1 with PC differences between T1 and C1 repl. 2.

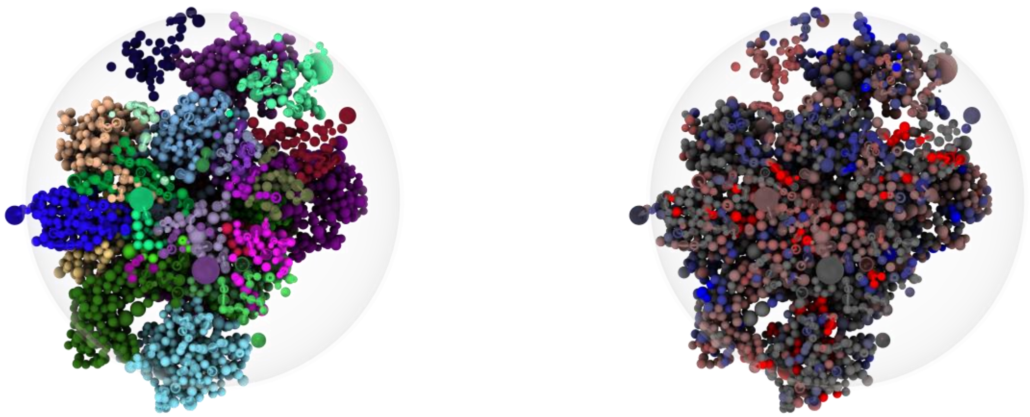

**Fig S21:** Left: Exemplary Chrom3D simulation model for 10A cells. Beads represent TADs sized according to genomic coverage. Chromosomes are colored distinctly. Right: The same model using sub-compartments for coloring each bead. The relative size of the model nucleus is indicated with a transparent sphere.

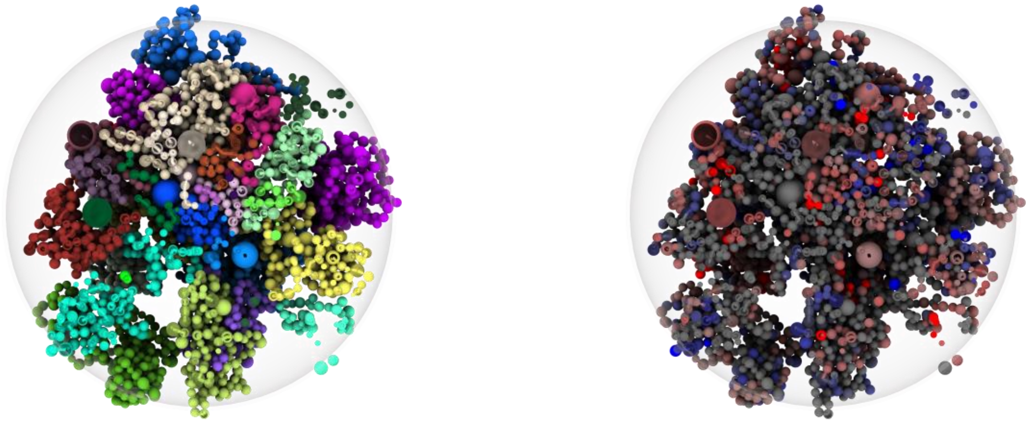

**Fig S22:** Left: Exemplary Chrom3D simulation model for T1 cells. Beads represent TADs sized according to genomic coverage. Chromosomes are colored distinctly. Right: The same model using sub-compartments for coloring each bead. The relative size of the model nucleus is indicated with a transparent sphere.

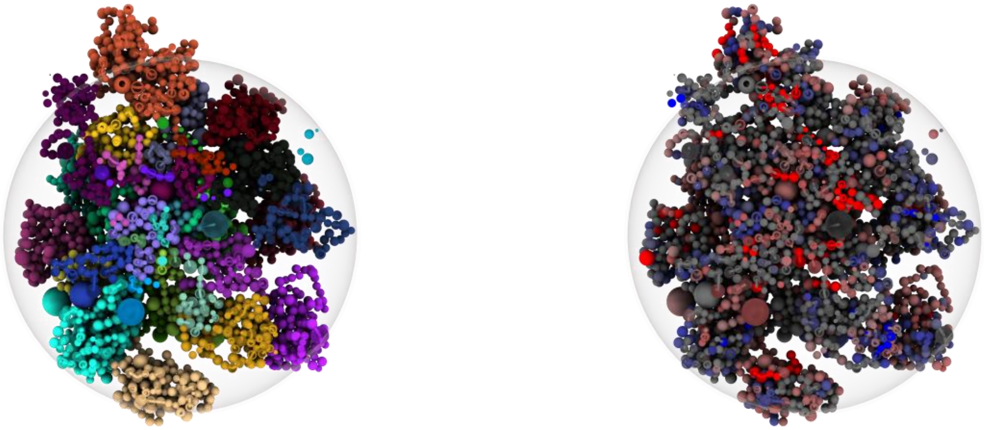

**Fig S23:** Left: Exemplary Chrom3D simulation model for C1 cells. Beads represent TADs sized according to genomic coverage. Chromosomes are colored distinctly. Right: The same model using sub-compartments for coloring each bead. The relative size of the model nucleus is indicated with a transparent sphere.

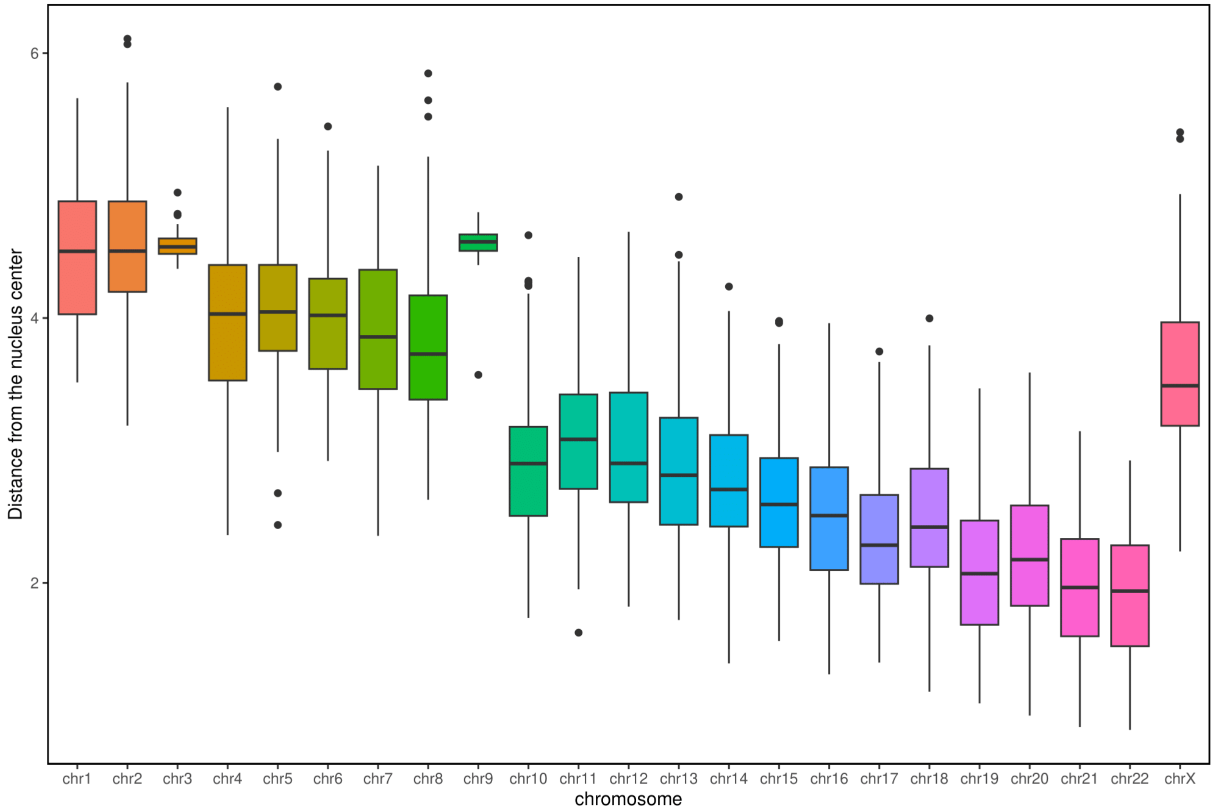


**Fig. S24:** Median chromosome distance from the nucleus center in 100 Chrom3D simulation models based on 10A.

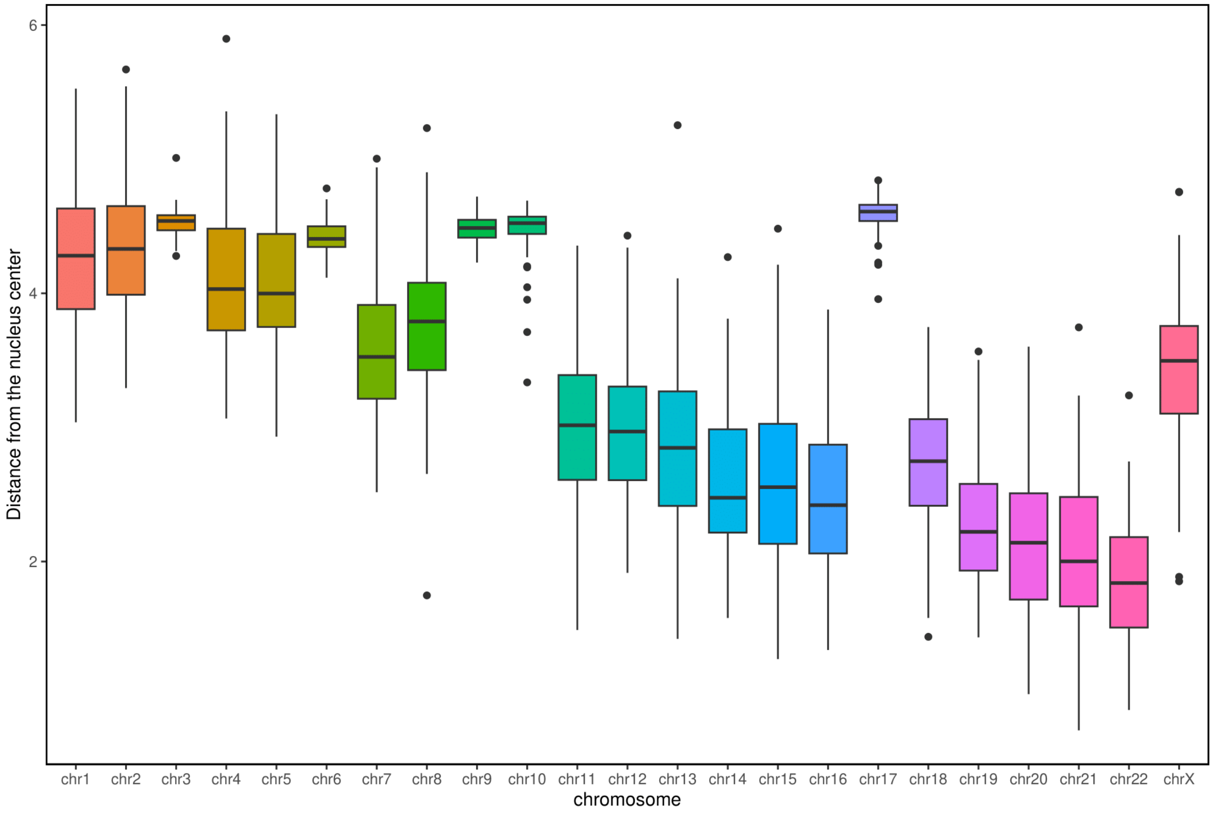


**Fig. S25:** Median chromosome distance from the nucleus center in 100 Chrom3D simulation models based on T1.

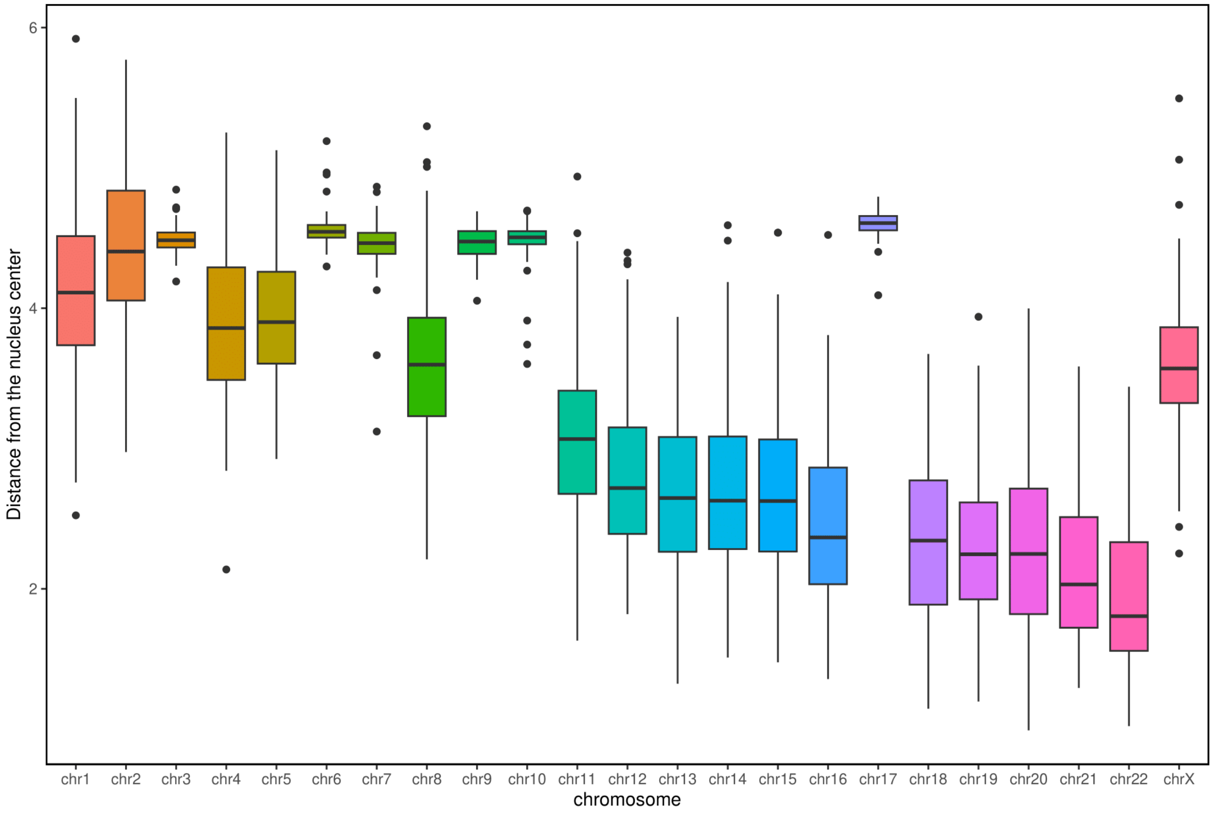


**Fig. S26:** Median chromosome distance from the nucleus center in 100 Chrom3D simulation models based on C1.

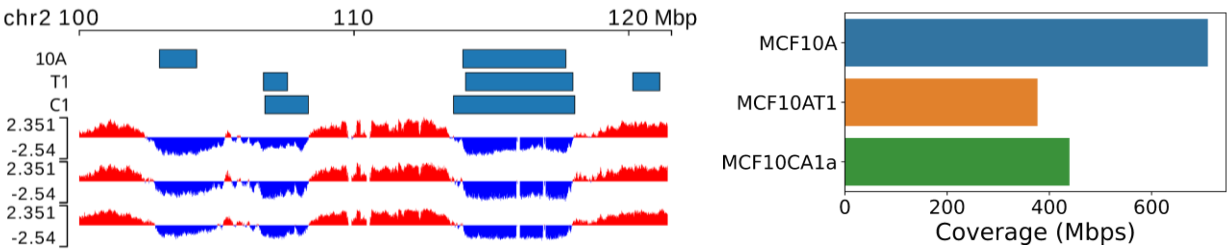


**Fig. S27**: Coverage of lamina associated domains (LADs).
Left: browser view of a region on chromosome 2 showing LMNB1 LADs for MCF10A (WT), MCF10AT1 (T1) and MCF10CA1a (C1) as blue bars. Principal component 1 for the same cell lines is shown below. Regions corresponding to A and B compartments are colored in red and blue respectively. Right: Genome-wide coverage of lamina associated domains.

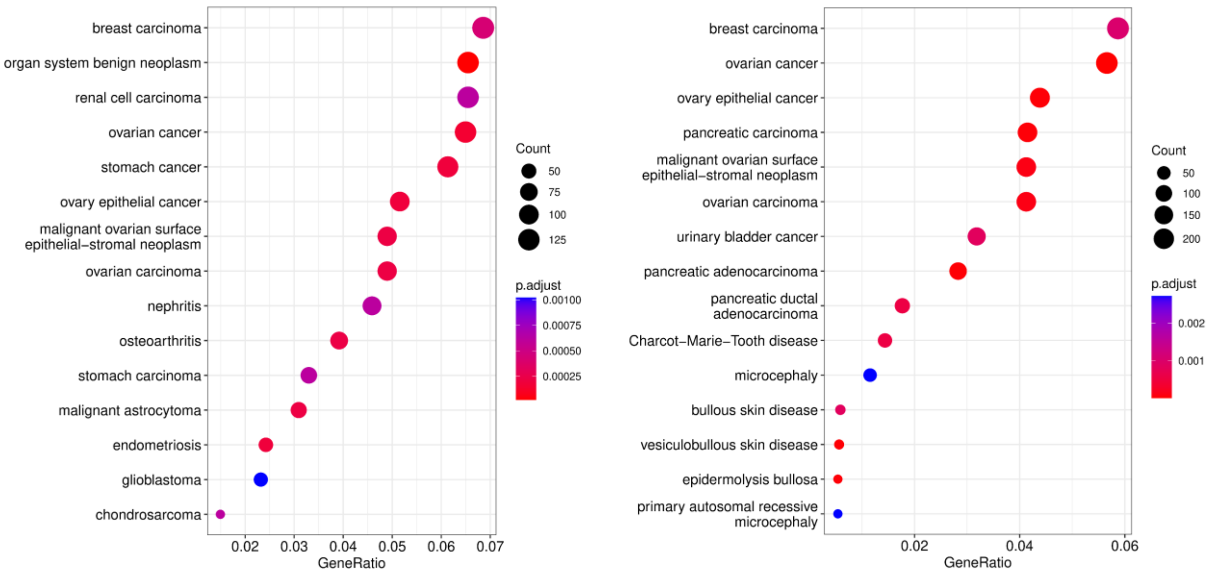


**Fig. S28**: Over-representation analysis for disease ontology (DO) terms for differentially expressed genes in MCF10AT1 (T1) (left) and MCF10CA1a (C1) (right) using MCF10A (10A) as contrast (lfc=0.5, p-value=0.01). Disease terms are sorted by the fraction of differentially expressed genes found in a given gene set. Dot size is proportional to the number of differentially expressed genes found in a given gene set. Dots are colored based on the statistical significance of the enrichment.

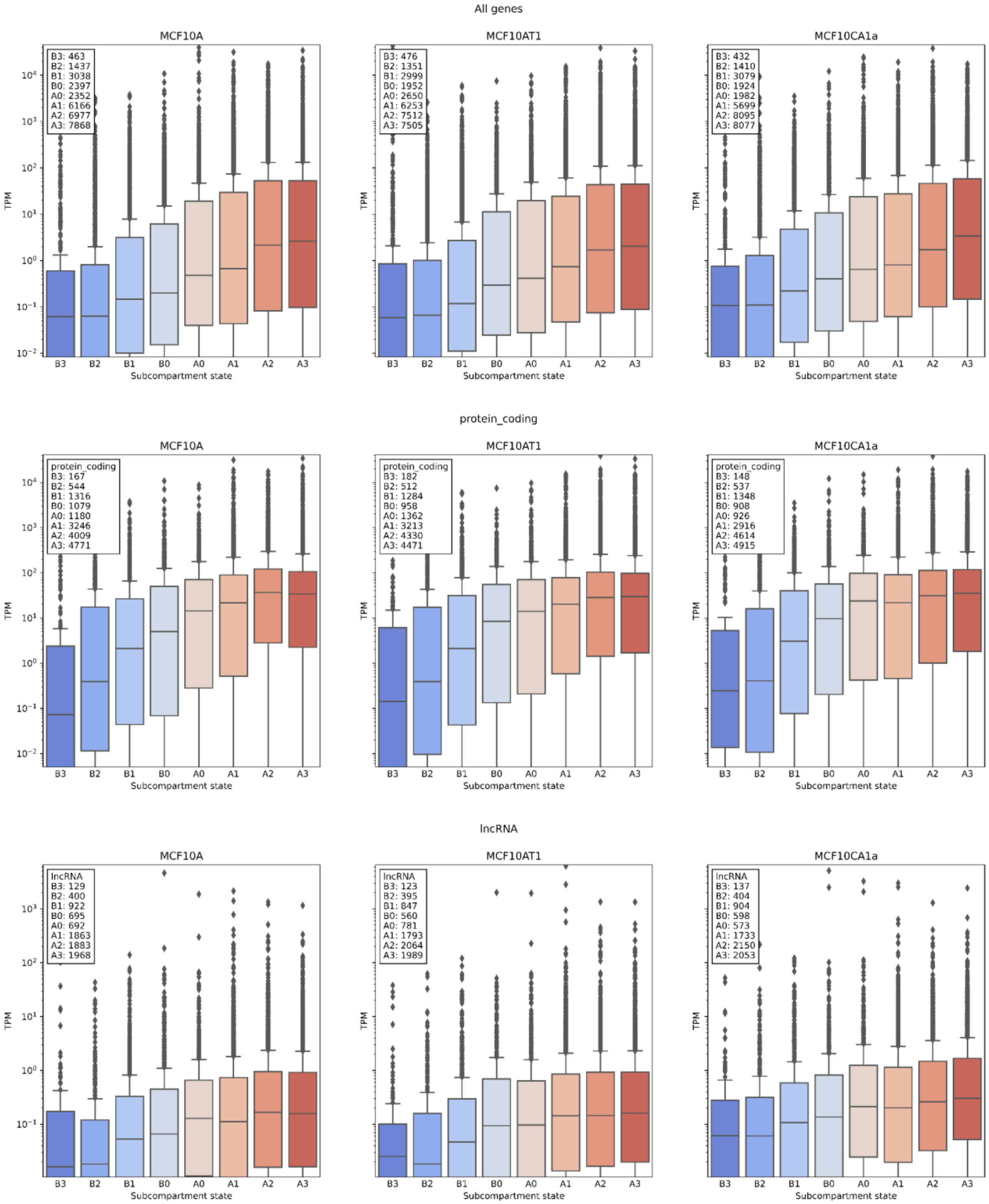


**Fig. S29** Distribution of expression levels (TPM) for each subcompartment across MCF10A (left panels), MCF10AT1 (middle panels) and MCF10CA1a (right panels). Rows 1 to 3 show the expression level for all genes, genes encoding for proteins, and long non-coding RNA, respectively. Legend shows the number of genes expressed in each subcompartment.

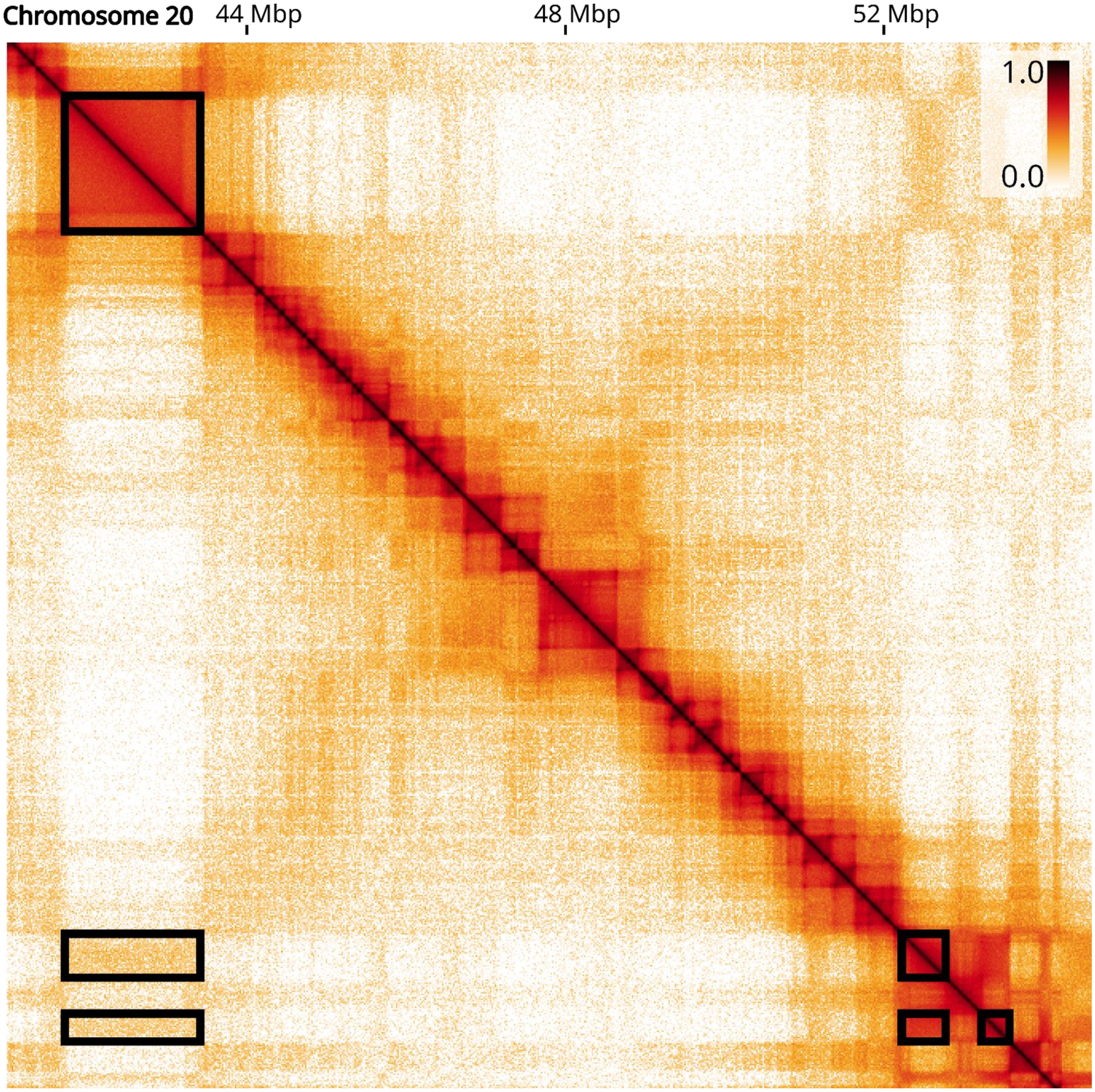


**Fig S30**: Hi-C matrix showing an example of a TAD clique of size 3. Domains involved in the TAD clique are enclosed in black rectangles.

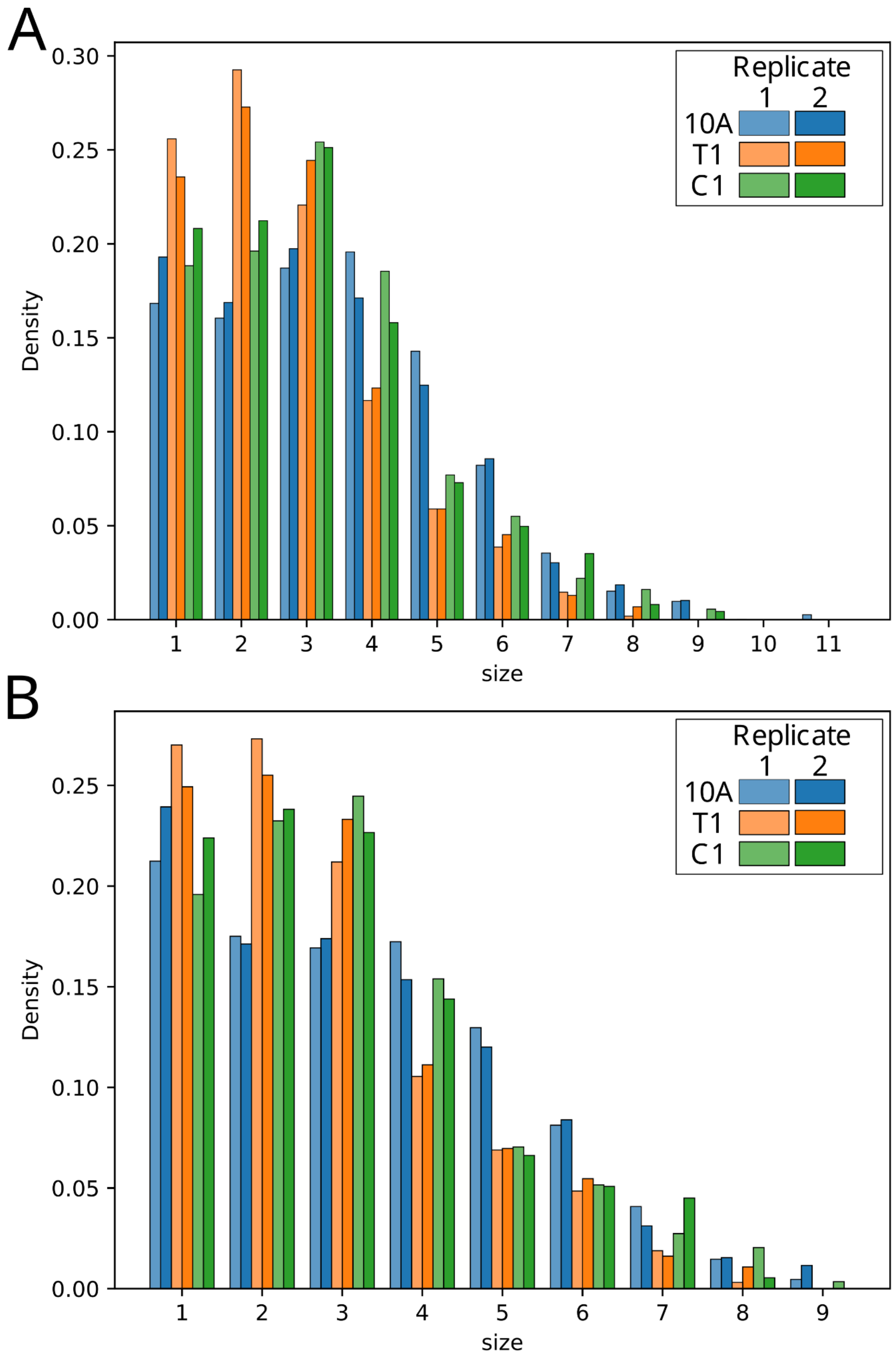


**Fig. S31**: **A** Bar plot showing the distribution of maximal clique sizes for MCF10A, MCF10AT1 and MCF10CA1a. Heights of bars are normalized such that bars for each replicate sum up to 1. **B** Bar plot showing the same data as panel A after masking cliques with one or more TADs overlapping with regions that are involved in translocation events in any of the 3 cell types studied.

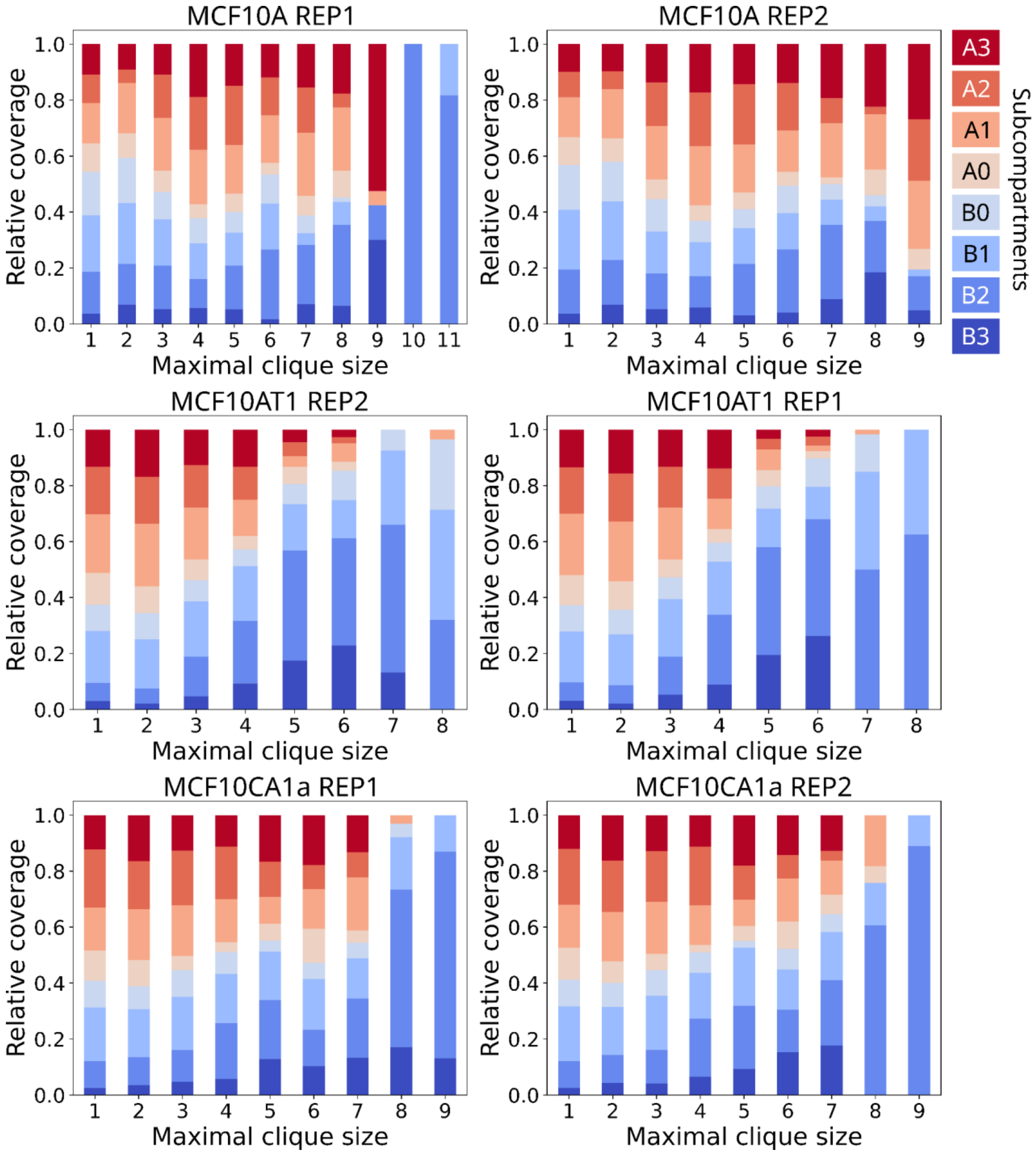


**Fig. S32**: Subcompartment composition of TAD cliques of size 1-11 in 10A (first row), T1 (second row) and C1 (third row). Cliques of size 1 and 2 are composed of singleton and binary TAD interactions not belonging to any clique.

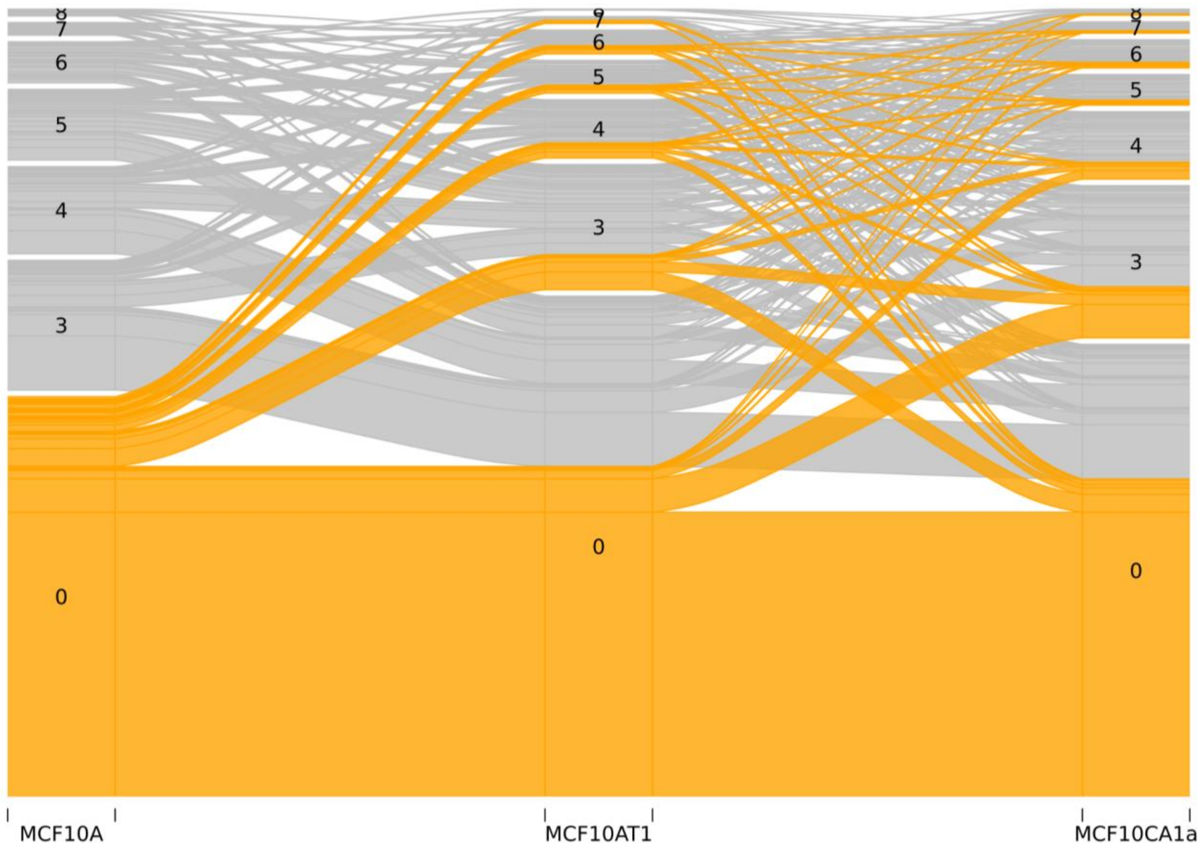


**Fig. S33**: Alluvial plot showing changes in TAD maximal clique size across the three cancer stages. The orange color highlights the alluvial path that starts with a non-clique (denoted as 0) in MCF10A (10A).

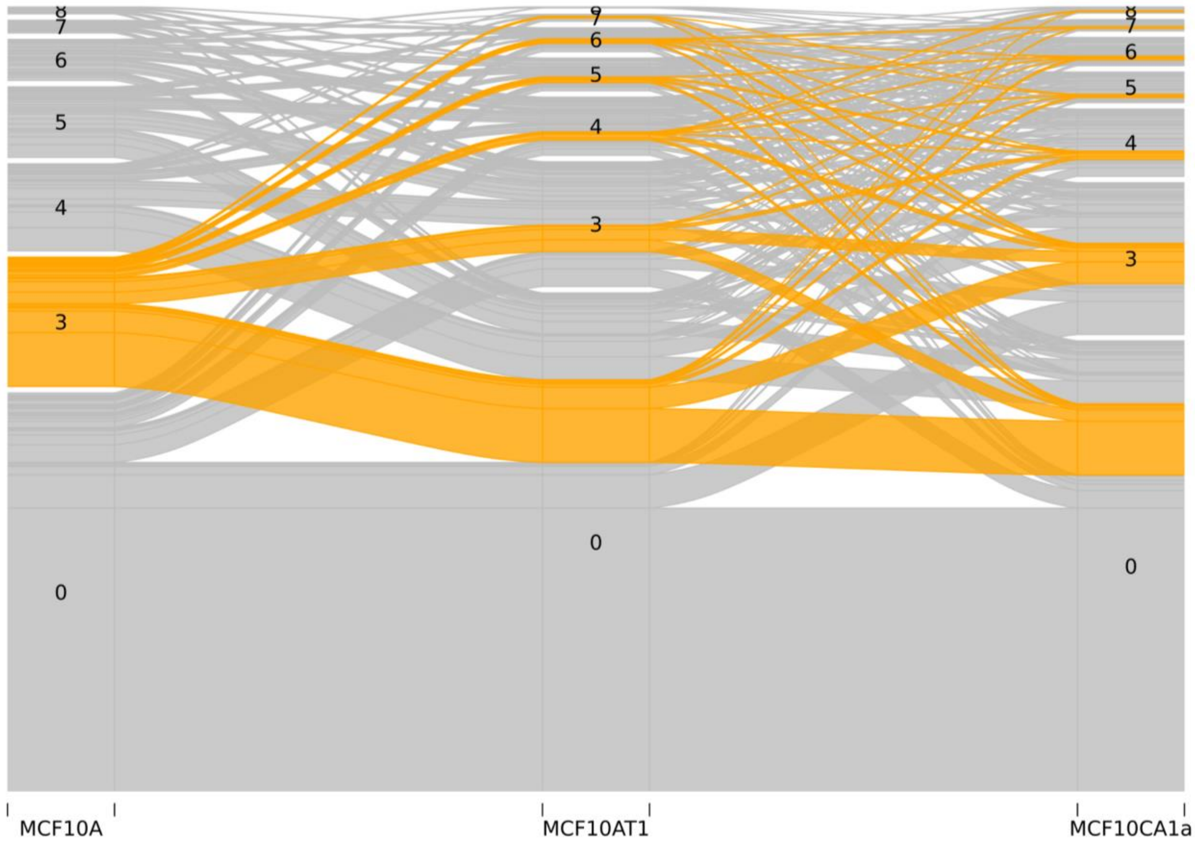


**Fig. S34**: Alluvial plot showing changes in TAD maximal clique size across the three cancer stages. The orange color highlights the alluvial path that starts with a clique of size 3 in MCF10A (10A).

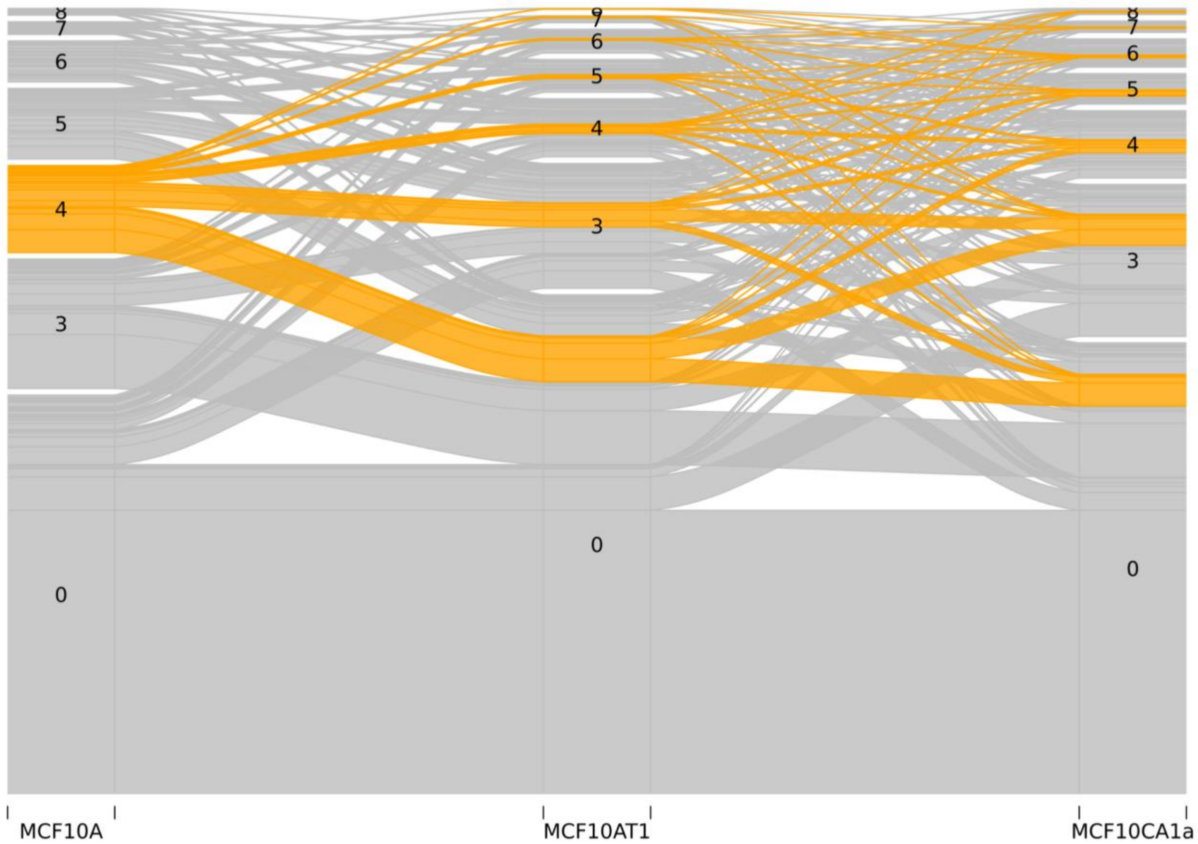


**Fig. S35**: Alluvial plot showing changes in TAD maximal clique size across the three cancer stages. The orange color highlights the alluvial path that starts with a clique of size 4 in MCF10A (10A).

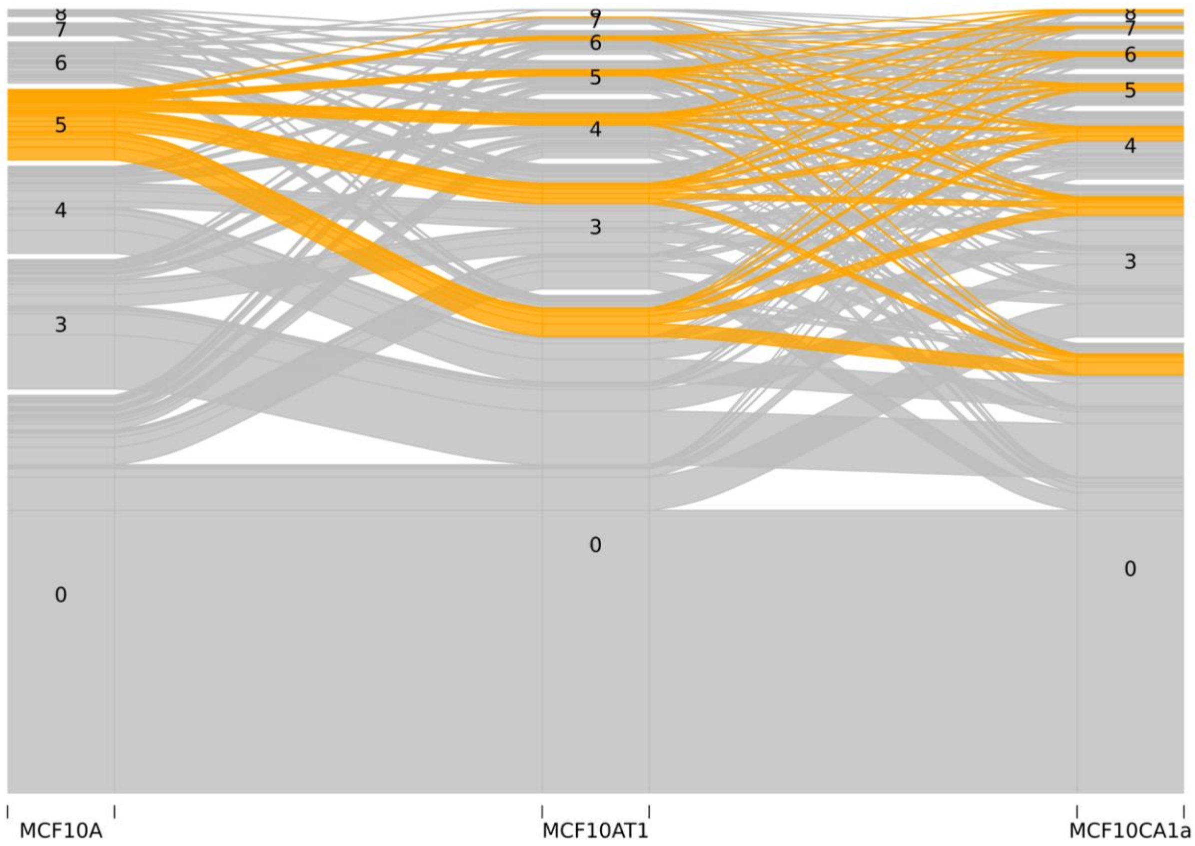


**Fig. S36**: Alluvial plot showing changes in TAD maximal clique size across the three cancer stages. The orange color highlights the alluvial path that starts with a clique of size 5 in MCF10A (10A).

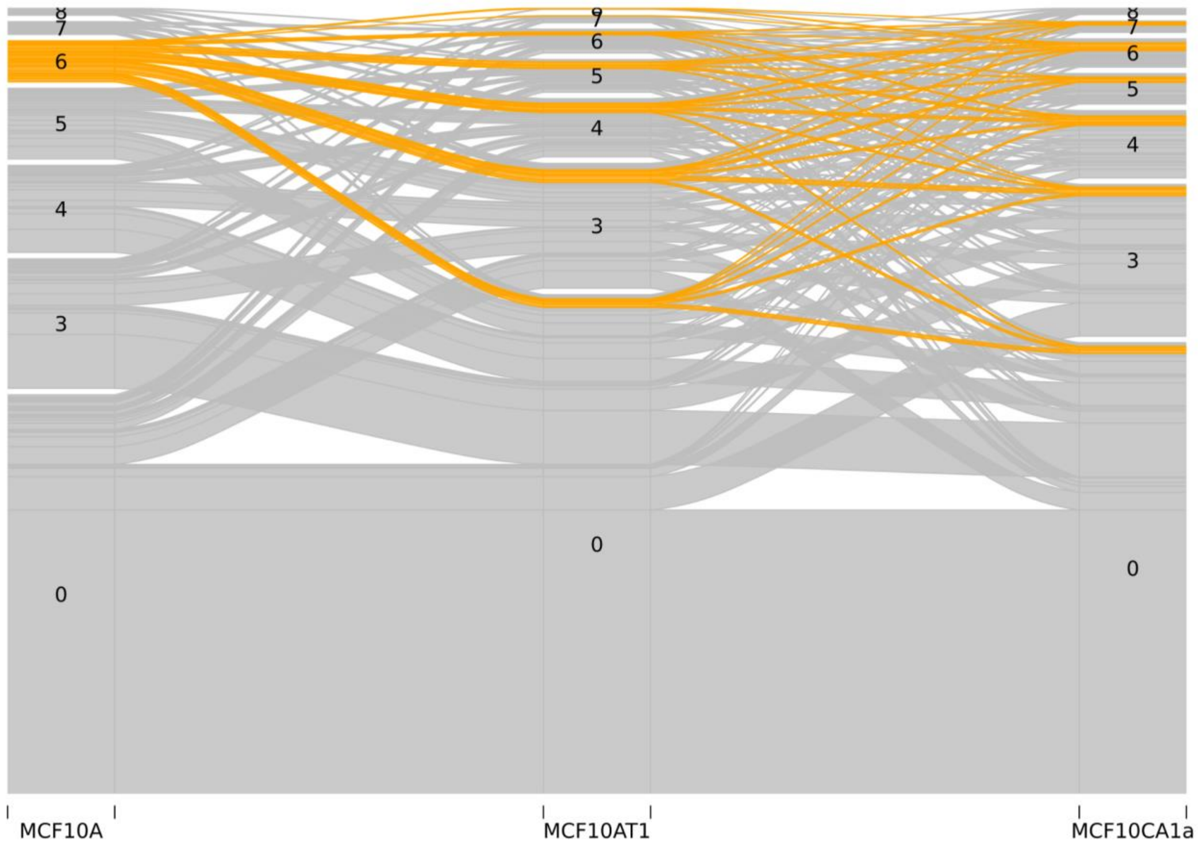


**Fig. S37**: Alluvial plot showing changes in TAD maximal clique size across the three cancer stages. The orange color highlights the alluvial path that starts with a clique of size 6 in MCF10A (10A).

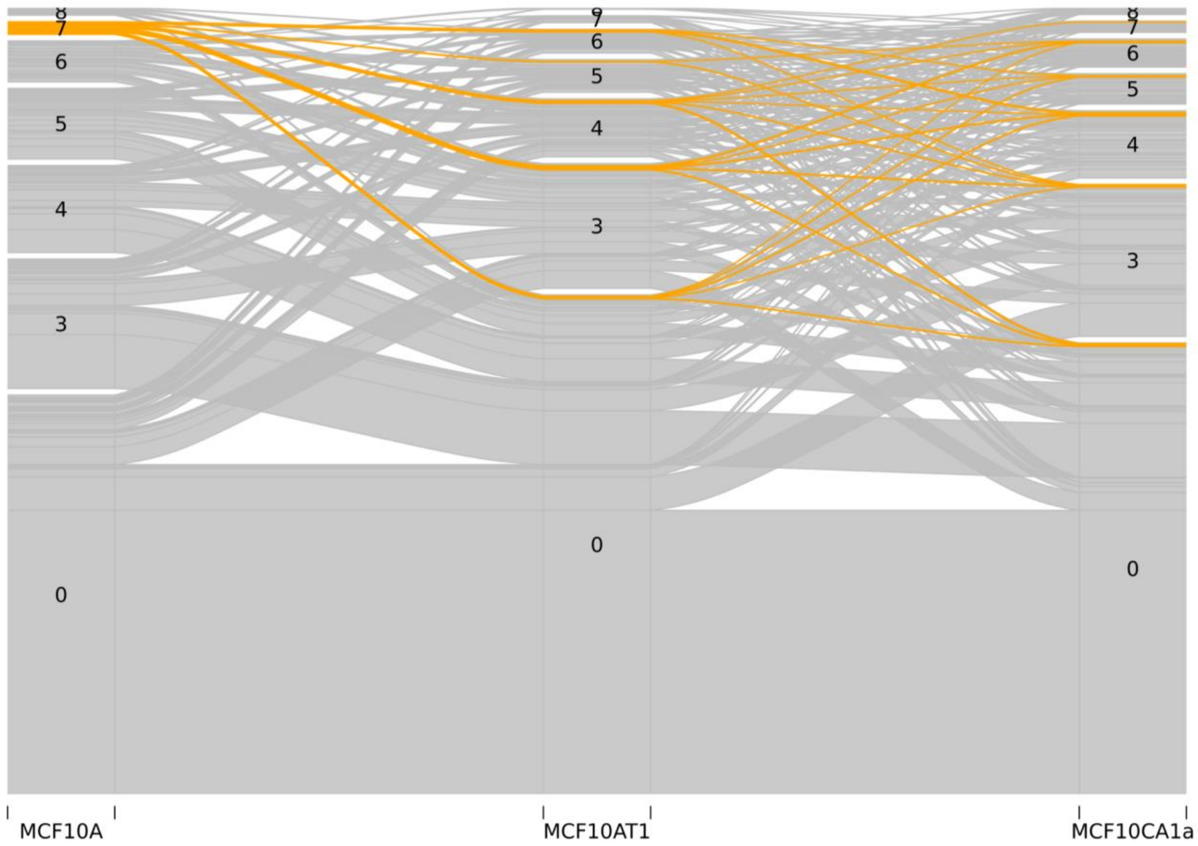


**Fig. S38**: Alluvial plot showing changes in TAD maximal clique size across the three cancer stages. The orange color highlights the alluvial path that starts with a clique of size 7 in MCF10A (10A).

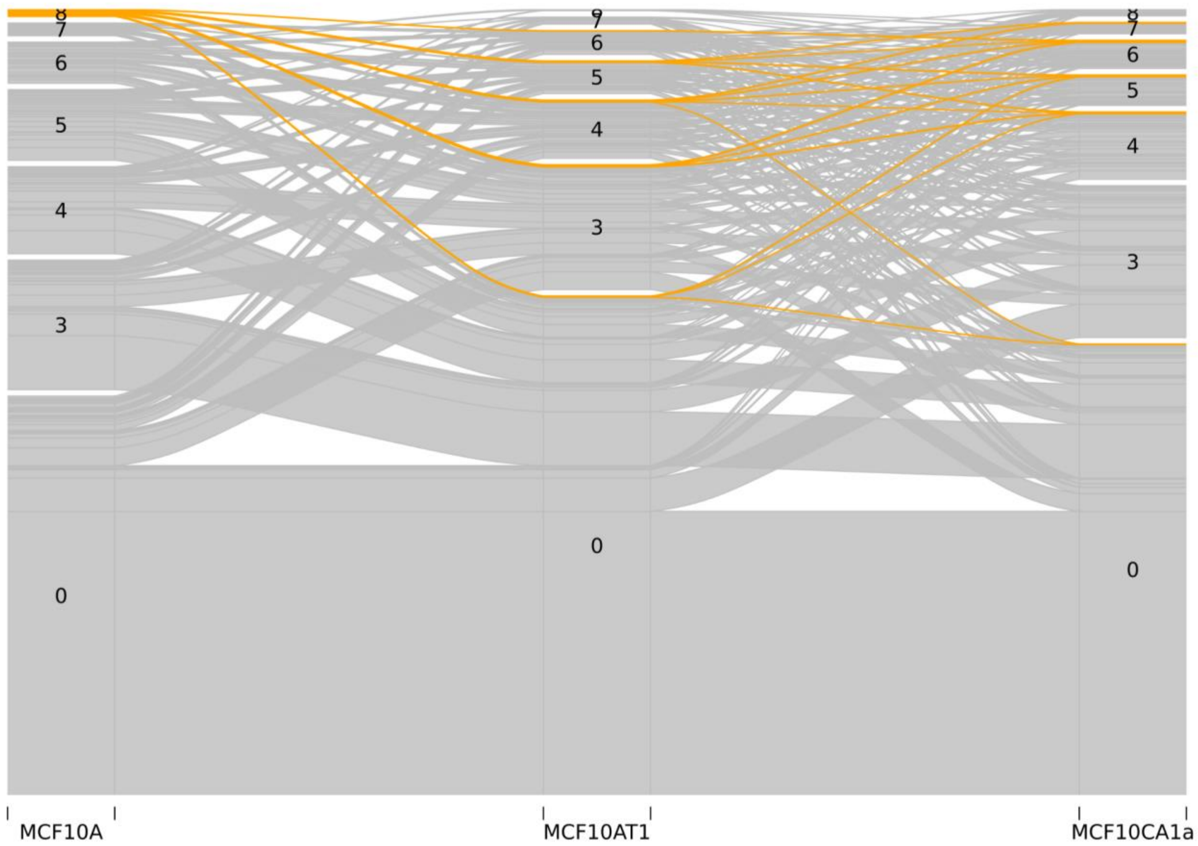


**Fig. S39**: Alluvial plot showing changes in TAD maximal clique size across the three cancer stages. The orange color highlights the alluvial path that starts with a clique of size 8 in MCF10A (10A).

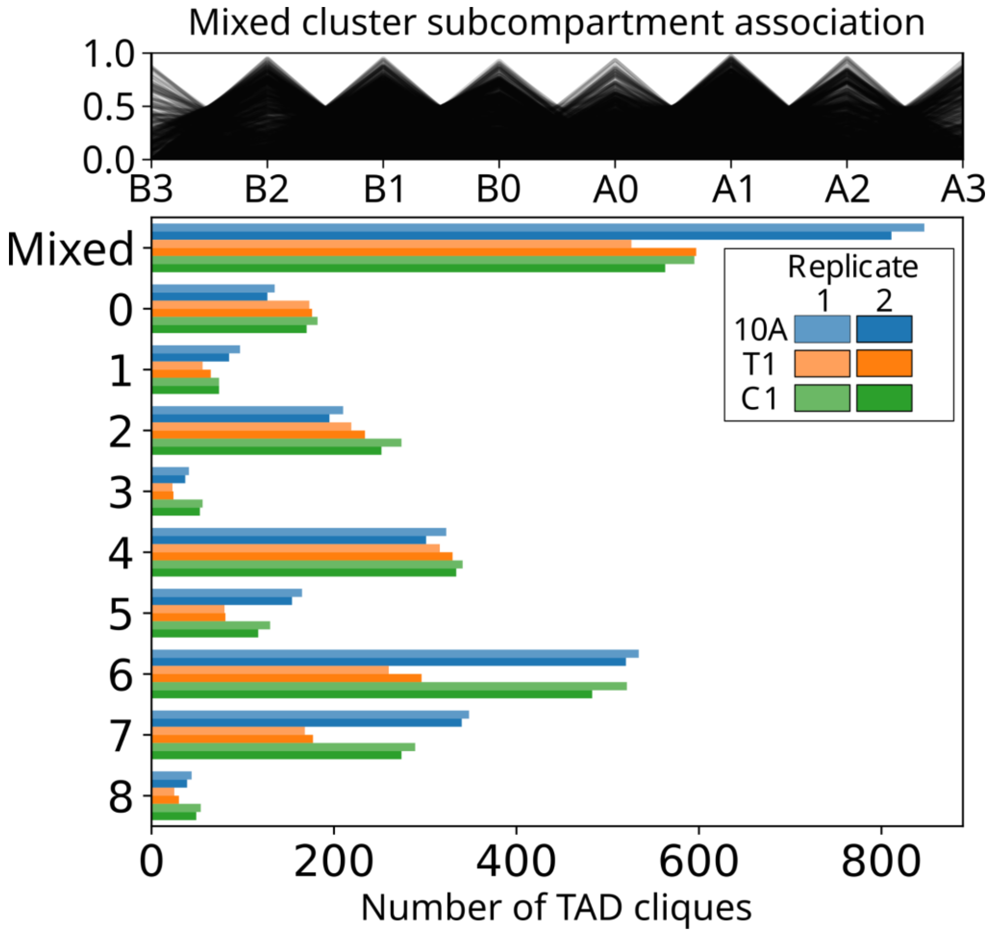


**Fig. S40: A**: Subcompartment enrichment for TAD cliques placed in the outlier ("mixed") cluster by HDBSCAN. **B**: Cluster sizes for TAD clique clusters shown in Fig. 3F (including the mixed cluster).

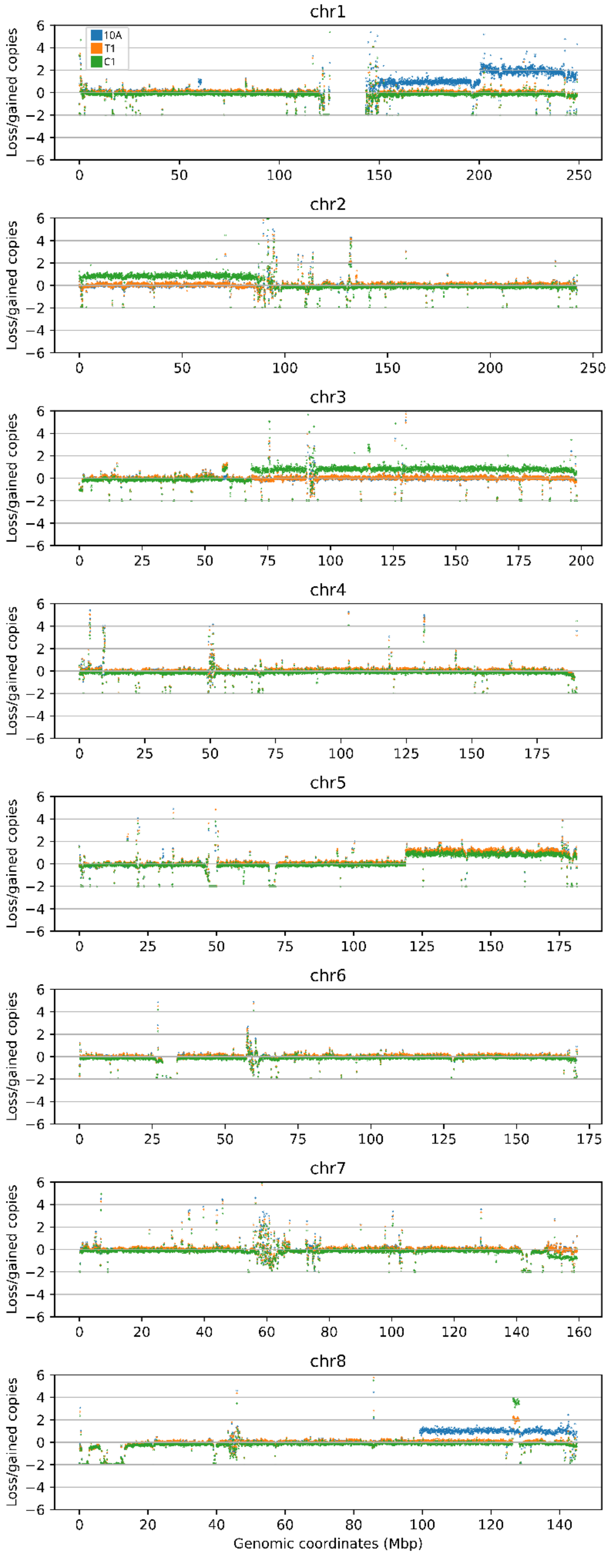


**Fig. S41**: Figure showing called CNVs across genome. Chrs 1-8

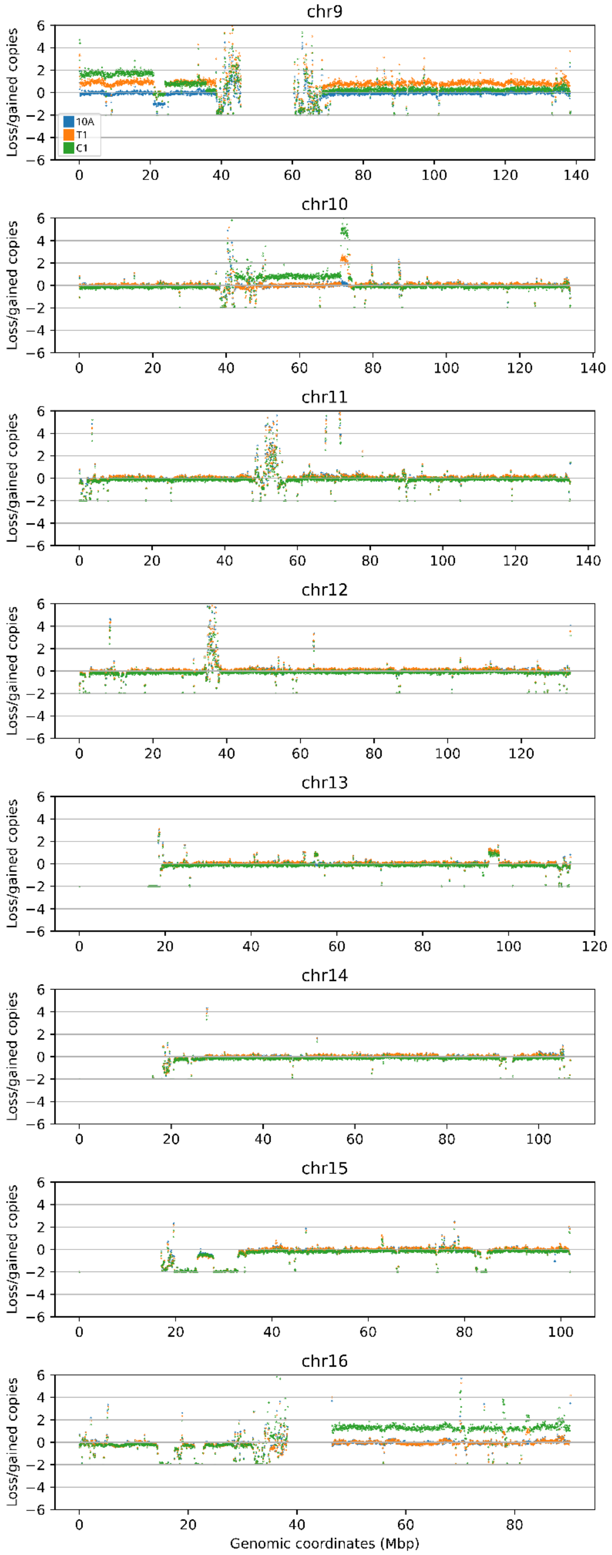


**Fig. S42**: Figure showing called CNVs across genome. Chrs 9-16

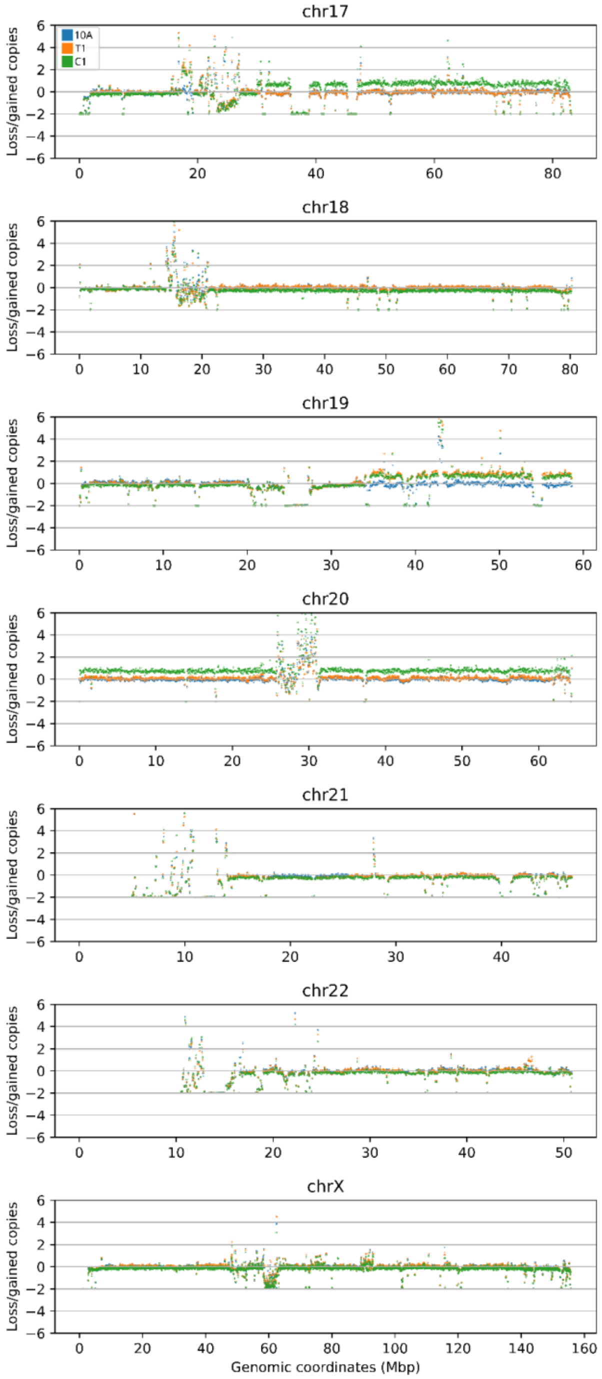


**Fig. S43**: Figure showing called CNVs across genome. Chrs 17-X

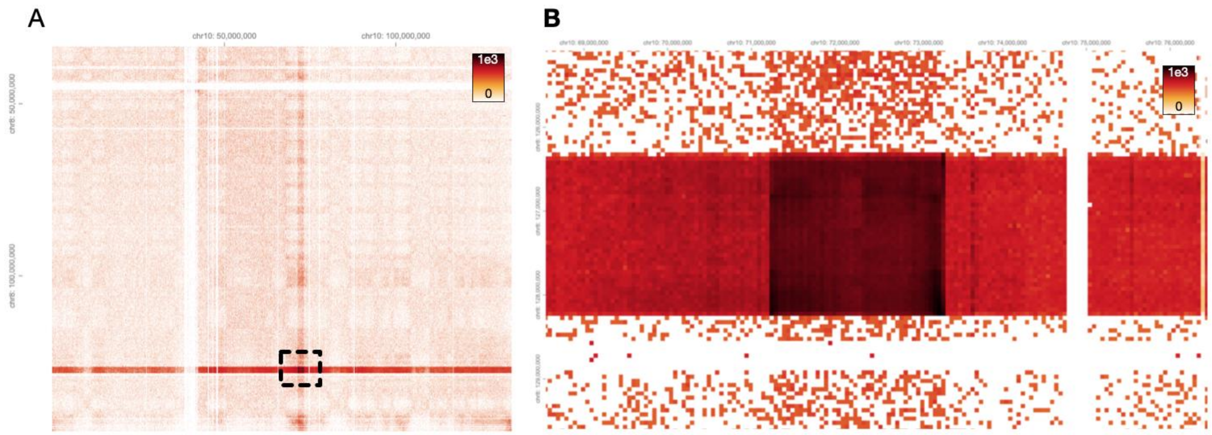


**Fig. S44**: **A**: Hi-C contact patterns (50 kbp bin resolution) in the genome region surrounding the *MYC*-insertion region on chromosome 8 (vertical axis) and chromosome 10 (horizontal axis) in C1 cells using the LOcal Iterative Correction (LOIC) balancing scheme, which takes copy number alterations int account [REF]. **B**: Zoom-in on the *MYC*-insertion region, showing enriched contacts between regions on 8 (126330000-128235000 bp; hg38) and chromosome 10 (71280000-73310000 bp; hg38).

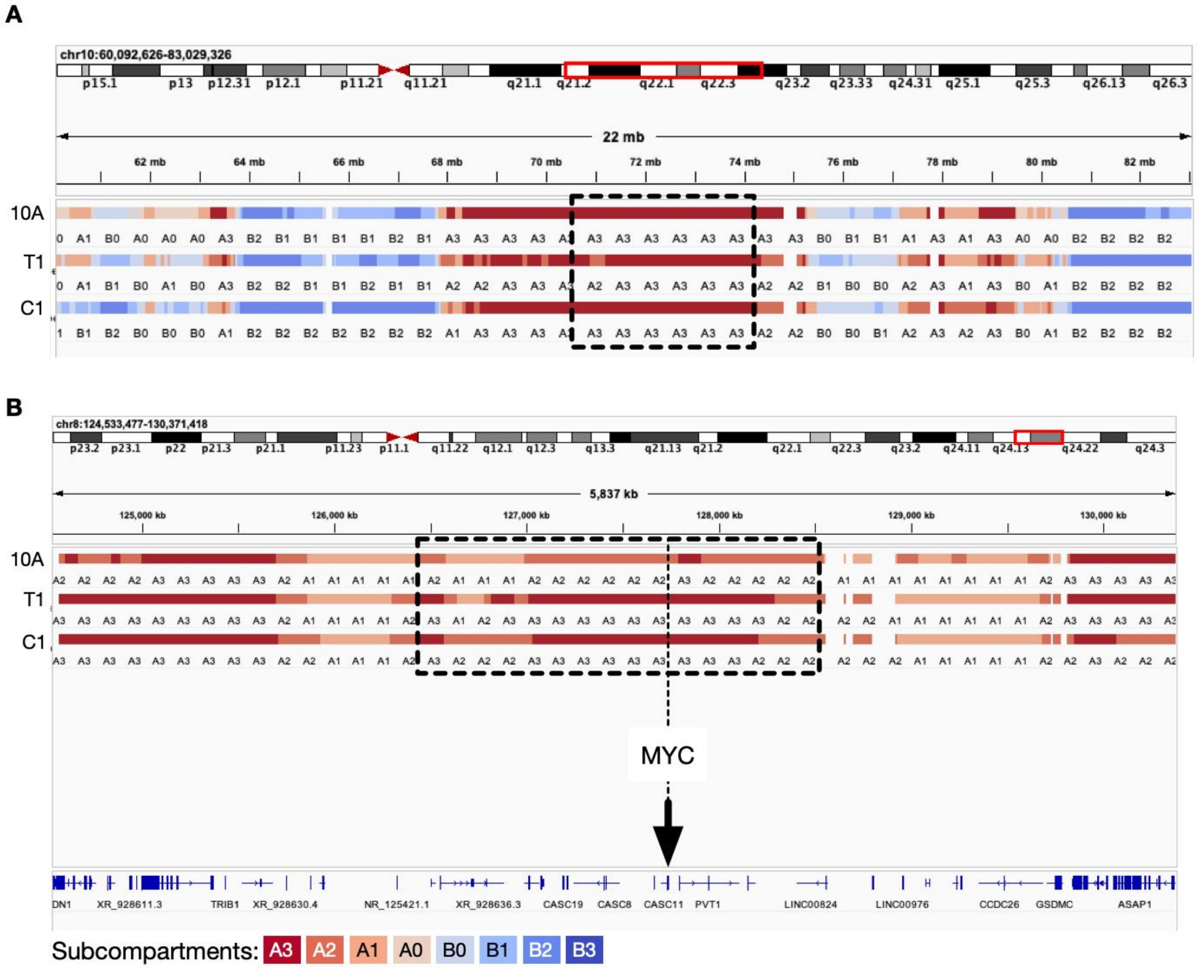


**Fig S45**: **A**: Genome browser view of subcompartments surrounding and including the amplified region in the three cell types on chromosome 10 (dotted lines) **B:** Genome browser view of subcompartments in the three cell types surrounding and including the amplified region on chromosome 8 (dotted lines)

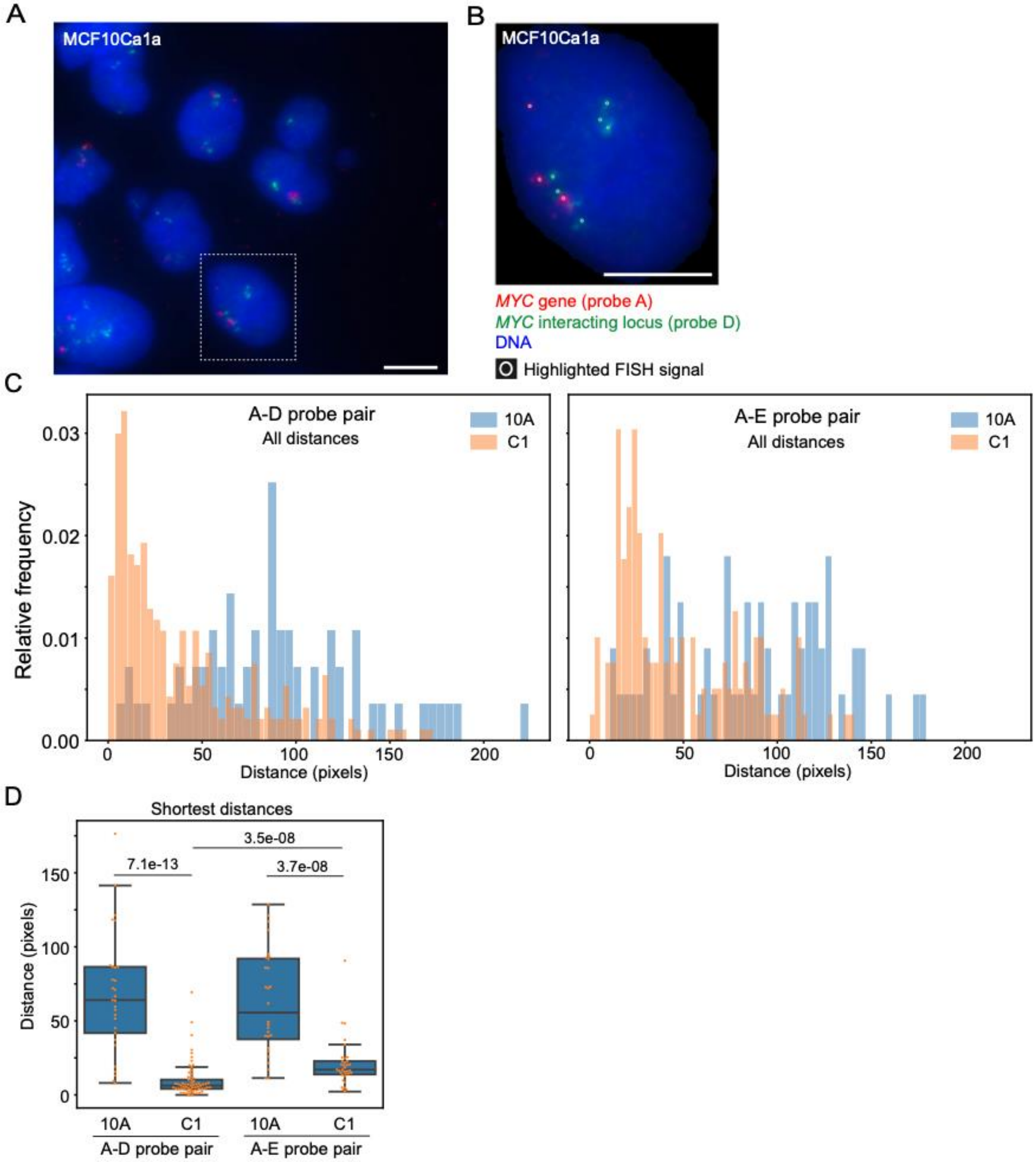


**Fig S46**: **A**: FISH image of the *MYC* gene (red) and the chromosome 10 enhancer (green; chr.10 position 73117711-73267436) in C1 cells. **B**: Zoom-in on region highlighted in A (dotted square). Bars, 7 µm. **C**: Left: Distribution of all distances (pixels) between the *MYC* gene and the enhancer probe in 10A (blue) and C1 (orange). Right: distribution of all distances (pixels) between the *MYC* gene and a downstream non-enhancer probe in 10A (blue) and C1 (orange). **D**: Quantification of shortest distance between the *MYC* probe and the nearest probe (either enhancer probe [D], or distal non-enhancer probe [E]). P-values are shown (Kolmogorov test).
