## Supplemental Tables for "Loss of multi-level 3D genome organization during breast cancer progression"

### Supplementary tables

**Table S1**: Hi-C interactions classified by type.

| | MCF10A_R1 | MCF10A_R2 | MCF10AT1_R1 | MCF10AT1_R2 | MCF10ACA1a_R1 | MCF10ACA1a_R2 |
|---|---|---|---|---|---|---|
| reported_pairs | 694.20 | 815.52 | 837.73 | 718.09 | 672.54 | 786.82 |
| valid_interaction | 664.17 | 774.48 | 821.28 | 704.73 | 650.15 | 760.91 |
| valid_interaction_rmdup | 598.02 | 681.51 | 687.67 | 603.61 | 559.65 | 639.96 |
| trans_interaction | 95.93 | 109.50 | 112.73 | 101.96 | 135.70 | 158.56 |
| cis_interaction | 502.10 | 572.01 | 574.94 | 501.65 | 423.95 | 481.40 |
| cis_shortRange | 106.91 | 121.64 | 105.22 | 89.85 | 75.64 | 85.27 |
| cis_longRange | 395.19 | 450.37 | 469.72 | 411.80 | 348.31 | 396.13 |
| cis_trans_ratio | 5.23 | 5.22 | 5.10 | 4.92 | 3.12 | 3.04 |
| cis_long_short_ratio | 3.70 | 3.70 | 4.46 | 4.58 | 4.61 | 4.65 |
| pct_valid_interaction_ff | 24.90 | 24.90 | 24.93 | 24.93 | 24.93 | 24.93 |
| pct_valid_interaction_rr | 24.92 | 24.92 | 24.95 | 24.96 | 24.96 | 24.96 |
| pct_valid_interaction_rf | 24.77 | 24.77 | 24.83 | 24.85 | 24.88 | 24.88 |
| pct_valid_interaction_fr | 25.40 | 25.41 | 25.30 | 25.26 | 25.24 | 25.24 |

**Table S2**: Coverage of compartment switches.

| MCF10A | MCF10AT1 | MCF10CA1a | Coverage (Mbp) | Rel. coverage (%) |
|---|---|---|---|---|
| B | B | B | 1024.22 | 37.58% |
| B | B | A | 119.23 | 4.37% |
| B | A | B | 76.1 | 2.79% |
| B | A | A | 146.15 | 5.36% |
| A | B | B | 101.57 | 3.73% |
| A | B | A | 60.85 | 2.23% |
| A | A | B | 97.91 | 3.59% |
| A | A | A | 1099.61 | 40.34% |

**Table S3**: Genome fraction (Mbp and relative) for WT, T1 and C1 for each of the 8 subcompartments.

| Subcompartment | Coverage (Mbp) | | | Coverage (relative) | | |
|---|---|---|---|---|---|---|
| | MCF10A | MCF10AT1 | MCF10CA1a | MCF10A | MCF10AT1 | MCF10CA1a |
| B3 | 164.82 | 180.49 | 172.04 | 5.34% | 5.84% | 5.57% |
| B2 | 509.71 | 449.93 | 468.55 | 16.50% | 14.57% | 15.17% |
| B1 | 517.25 | 581.01 | 580.49 | 16.75% | 18.81% | 18.80% |
| B0 | 355.62 | 268.17 | 251.65 | 11.52% | 8.68% | 8.15% |
| A0 | 213.53 | 253.13 | 216.38 | 6.91% | 8.20% | 7.01% |
| A1 | 546.21 | 538.50 | 482.78 | 17.69% | 17.44% | 15.63% |
| A2 | 398.25 | 438.56 | 523.43 | 12.90% | 14.20% | 16.95% |
| A3 | 382.88 | 378.47 | 392.95 | 12.40% | 12.26% | 12.72% |

**Table S4**: Wilcoxon rank sum test results of the median subcompartment distance from the nucleus center in each Chrom3D simulation model per condition and subcompartment pair.

| Condition 1 | Condition 2 | Subcomp. | P-value |
|---|---|---|---|
| 10A | C1 | A3 | < 2.2e-16 |
| 10A | C1 | A2 | 0.4977 |
| 10A | C1 | A1 | < 2.2e-16 |
| 10A | C1 | A0 | 3.175e-12 |
| 10A | C1 | B0 | 7.824e-05 |
| 10A | C1 | B1 | 4.367e-16 |
| 10A | C1 | B2 | 1.982e-12 |
| 10A | C1 | B3 | 1.369e-08 |
| 10A | T1 | A3 | < 2.2e-16 |
| 10A | T1 | A2 | 8.995e-07 |
| 10A | T1 | A1 | < 2.2e-16 |
| 10A | T1 | A0 | 0.001466 |
| 10A | T1 | B0 | 1.548e-06 |
| 10A | T1 | B1 | 0.0001672 |
| 10A | T1 | B2 | 0.6259 |
| 10A | T1 | B3 | 3.16e-13 |
| C1 | T1 | A3 | 0.0005997 |
| C1 | T1 | A2 | 1.586e-07 |
| C1 | T1 | A1 | 0.9251 |
| C1 | T1 | A0 | 6.97e-06 |
| C1 | T1 | B0 | 0.1222 |
| C1 | T1 | B1 | 2.925e-10 |
| C1 | T1 | B2 | 4.28e-16 |
| C1 | T1 | B3 | 0.05464 |

**Table S5**: Number of differentially expressed genes (lfc> 0.5; pvalue<0.01).

| contrast | condition | genes | DE genes | downreg | upreg |
|---|---|---|---|---|---|
| 10A | T1 | 21082 | 3180 | 1659 | 1521 |
| T1 | C1 | 21082 | 7510 | 3739 | 3771 |
| 10A | C1 | 21082 | 8362 | 4169 | 4193 |

**Table S6**: Table of subcompartment switches when comparing 10A with T1. Statistical significance was computed using the one-sided binomial test.

| Contrast (10A) | Condition (T1) | pvalue (downreg) | pvalue (upreg) |
|---|---|---|---|
| B3 | B3 | 1.000 | 1.000 |
| B3 | B2 | 0.938 | 0.500 |
| B3 | B1 | 0.875 | 0.125 |
| B3 | B0 | 1.000 | 1.000 |
| B3 | A0 | 1.000 | 1.000 |
| B3 | A1 | 1.000 | 1.000 |
| B3 | A2 | 1.000 | 1.000 |
| B3 | A3 | 1.000 | 1.000 |
| B2 | B3 | 0.227 | 0.813 |
| B2 | B2 | 1.000 | 1.000 |
| B2 | B1 | 1.000 | 0.059 |
| B2 | B0 | 1.000 | 0.500 |
| B2 | A0 | 1.000 | 0.688 |

| | | | |
|---|---|---|---|
| B2 | A1 | 1.000 | 0.500 |
| B2 | A2 | 1.000 | 1.000 |
| B2 | A3 | 1.000 | 0.500 |
| B1 | B3 | 0.500 | 1.000 |
| B1 | B2 | 0.000 | 0.982 |
| B1 | B1 | 1.000 | 1.000 |
| B1 | B0 | 1.000 | 0.046 |
| B1 | A0 | 1.000 | 0.072 |
| B1 | A1 | 0.945 | 0.004 |
| B1 | A2 | 1.000 | 0.125 |
| B1 | A3 | 0.063 | 1.000 |
| B0 | B3 | 1.000 | 1.000 |
| B0 | B2 | 0.125 | 1.000 |
| B0 | B1 | 0.000 | 0.989 |
| B0 | B0 | 1.000 | 1.000 |
| B0 | A0 | 0.125 | 1.000 |
| B0 | A1 | 0.989 | 0.095 |
| B0 | A2 | 1.000 | 0.063 |
| B0 | A3 | 0.500 | 0.500 |

| | | | |
|---|---|---|---|
| A0 | B3 | 0.250 | 1.000 |
| A0 | B2 | 1.000 | 0.688 |
| A0 | B1 | 0.004 | 0.975 |
| A0 | B0 | 1.000 | 0.250 |
| A0 | A0 | 1.000 | 1.000 |
| A0 | A1 | 0.846 | 0.166 |
| A0 | A2 | 0.344 | 0.313 |
| A0 | A3 | 1.000 | 1.000 |
| A1 | B3 | 1.000 | 1.000 |
| A1 | B2 | 0.250 | 1.000 |
| A1 | B1 | 0.172 | 1.000 |
| A1 | B0 | 0.038 | 0.961 |
| A1 | A0 | 0.271 | 0.928 |
| A1 | A1 | 1.000 | 1.000 |
| A1 | A2 | 0.081 | 0.500 |
| A1 | A3 | 0.773 | 0.019 |
| A2 | B3 | 1.000 | 1.000 |
| A2 | B2 | 1.000 | 1.000 |
| A2 | B1 | 0.500 | 1.000 |

| | | | |
|---|---|---|---|
| A2 | B0 | 0.500 | 1.000 |
| A2 | A0 | 0.891 | 0.938 |
| A2 | A1 | 0.960 | 0.622 |
| A2 | A2 | 1.000 | 1.000 |
| A2 | A3 | 0.953 | 0.868 |
| A3 | B3 | 1.000 | 1.000 |
| A3 | B2 | 1.000 | 1.000 |
| A3 | B1 | 1.000 | 1.000 |
| A3 | B0 | 0.875 | 0.875 |
| A3 | A0 | 1.000 | 1.000 |
| A3 | A1 | 0.500 | 0.997 |
| A3 | A2 | 0.105 | 0.229 |
| A3 | A3 | 1.000 | 1.000 |

**Table S7**: Table of subcompartment switches when comparing 10A with C1. Statistical significance was computed using the one-sided binomial test.

| Contrast (10A) | Condition (C1) | pvalue (downreg) | pvalue (upreg) |
|---|---|---|---|

| | | | |
|---|---|---|---|
| B3 | B3 | 1.000 | 1.000 |
| B3 | B2 | 0.377 | 0.212 |
| B3 | B1 | 0.969 | 0.031 |
| B3 | B0 | 1.000 | 1.000 |
| B3 | A0 | 1.000 | 1.000 |
| B3 | A1 | 1.000 | 1.000 |
| B3 | A2 | 1.000 | 1.000 |
| B3 | A3 | 1.000 | 1.000 |
| B2 | B3 | 0.828 | 0.910 |
| B2 | B2 | 1.000 | 1.000 |
| B2 | B1 | 1.000 | 0.229 |
| B2 | B0 | 0.637 | 0.500 |
| B2 | A0 | 0.997 | 0.623 |
| B2 | A1 | 0.813 | 0.194 |
| B2 | A2 | 1.000 | 0.500 |
| B2 | A3 | 1.000 | 1.000 |
| B1 | B3 | 0.188 | 1.000 |
| B1 | B2 | 0.000 | 0.868 |
| B1 | B1 | 1.000 | 1.000 |

| | | | |
|---|---|---|---|
| B1 | B0 | 0.885 | 0.077 |
| B1 | A0 | 0.997 | 0.908 |
| B1 | A1 | 0.708 | 0.001 |
| B1 | A2 | 0.746 | 0.013 |
| B1 | A3 | 0.969 | 1.000 |
| B0 | B3 | 1.000 | 1.000 |
| B0 | B2 | 0.637 | 1.000 |
| B0 | B1 | 0.196 | 0.960 |
| B0 | B0 | 1.000 | 1.000 |
| B0 | A0 | 0.891 | 0.980 |
| B0 | A1 | 0.394 | 0.101 |
| B0 | A2 | 0.145 | 0.004 |
| B0 | A3 | 0.688 | 0.063 |
| A0 | B3 | 0.500 | 0.500 |
| A0 | B2 | 0.019 | 0.623 |
| A0 | B1 | 0.008 | 0.172 |
| A0 | B0 | 0.344 | 0.090 |
| A0 | A0 | 1.000 | 1.000 |
| A0 | A1 | 0.994 | 0.087 |

| | | | |
|---|---|---|---|
| A0 | A2 | 0.166 | 0.004 |
| A0 | A3 | 0.938 | 0.500 |
| A1 | B3 | 1.000 | 1.000 |
| A1 | B2 | 0.500 | 0.927 |
| A1 | B1 | 0.428 | 1.000 |
| A1 | B0 | 0.705 | 0.941 |
| A1 | A0 | 0.014 | 0.952 |
| A1 | A1 | 1.000 | 1.000 |
| A1 | A2 | 0.165 | 0.000 |
| A1 | A3 | 0.231 | 0.001 |
| A2 | B3 | 1.000 | 1.000 |
| A2 | B2 | 1.000 | 1.000 |
| A2 | B1 | 0.500 | 0.996 |
| A2 | B0 | 0.965 | 0.999 |
| A2 | A0 | 0.928 | 0.999 |
| A2 | A1 | 0.879 | 1.000 |
| A2 | A2 | 1.000 | 1.000 |
| A2 | A3 | 0.913 | 0.584 |
| A3 | B3 | 1.000 | 1.000 |

| | | | |
|---|---|---|---|
| A3 | B2 | 1.000 | 1.000 |
| A3 | B1 | 0.188 | 0.250 |
| A3 | B0 | 0.688 | 1.000 |
| A3 | A0 | 0.313 | 1.000 |
| A3 | A1 | 0.849 | 1.000 |
| A3 | A2 | 0.126 | 0.500 |
| A3 | A3 | 1.000 | 1.000 |

**Table S8**: Contingency table with number of DE genes for Fisher's exact test comparing WT with T1 (Fisher test: 0.302975206611570248; P=4.0438076214854218e-09).

| | delta < 0 (more closed/B-like) | delta > 0 (more open/A-like) |
|---|---|---|
| lfc ≥ 2 | 72 | 132 |
| lfc ≤ -2 | 132 | 72 |

**Table S9**: Contingency table with number of DE genes for Fisher's exact test comparing WT with C1 (Fisher test: 0.42376178720527073; P=6.6567467681675556e-13).

| | delta < 0 (more closed/B-like) | delta > 0 (more open/A-like) |
|---|---|---|
| lfc ≥ 2 | 210 | 373 |
| lfc ≤ -2 | 341 | 259 |

**Table S10**: P-value table computed using the Mcnemar test to assess changes of clique sizes across conditions.

| **cond1** | **cond2** | **ratio** | **clique_size** | **mcnemar_stat** | **mcnemar_pval** |
|---|---|---|---|---|---|

| | | | | | |
|---|---|---|---|---|---|
| WT | T1 | 0.74 | 1 | 789 | 2.20E-09 |
| WT | T1 | 0.58 | 2 | 690 | 1.30E-31 |
| WT | T1 | 0.83 | 3 | 807 | 0.023 |
| WT | T1 | 1.53 | 4 | 477 | 8.60E-11 |
| WT | T1 | 2.27 | 5 | 241 | 4.70E-23 |
| WT | T1 | 2 | 6 | 158 | 9.60E-18 |
| WT | T1 | 2.38 | 7 | 60 | 2.70E-06 |
| WT | T1 | 3.83 | 8 | 8 | 5.00E-15 |
| WT | T1 | inf | 9 | 0 | 4.50E-13 |
| WT | T1 | inf | 10 | 0 | 1 |
| WT | T1 | inf | 11 | 0 | 1 |
| WT | C1 | 0.91 | 1 | 770 | 0.65 |
| WT | C1 | 0.81 | 2 | 690 | 0.004 |
| WT | C1 | 0.76 | 3 | 807 | 7.30E-08 |
| WT | C1 | 1.07 | 4 | 700 | 0.14 |
| WT | C1 | 1.78 | 5 | 315 | 1.20E-11 |
| WT | C1 | 1.6 | 6 | 225 | 2.10E-07 |
| WT | C1 | 1.15 | 7 | 90 | 0.024 |
| WT | C1 | 1.39 | 8 | 66 | 0.45 |

| | | | | | |
|---|---|---|---|---|---|
| WT | C1 | 2 | 9 | 23 | 0.025 |
| WT | C1 | inf | 10 | 0 | 1 |
| WT | C1 | inf | 11 | 0 | 1 |

**Table S11**: Table of numbers of members in HDBSCAN clusters in 10A, T1 and C1.

| cluster | MCF10A REP1 | MCF10A REP2 | MCF10AT1 REP1 | MCF10AT1 REP2 | MCF10CA1a REP1 | MCF10CA1a REP2 |
|---|---|---|---|---|---|---|
| Outliers | 847 | 811 | 526 | 597 | 595 | 563 |
| 0 | 135 | 127 | 173 | 176 | 182 | 170 |
| 1 | 97 | 85 | 56 | 65 | 74 | 74 |
| 2 | 210 | 195 | 219 | 234 | 274 | 252 |
| 3 | 41 | 37 | 23 | 24 | 56 | 53 |
| 4 | 323 | 301 | 316 | 330 | 341 | 334 |
| 5 | 165 | 154 | 80 | 81 | 130 | 117 |
| 6 | 534 | 520 | 260 | 296 | 521 | 483 |
| 7 | 348 | 340 | 168 | 177 | 289 | 274 |
| 8 | 44 | 39 | 25 | 30 | 54 | 49 |

**Table S12**: P-value table computed using the Mcnemar test to assess changes in associations with clique clusters across conditions.

| cond1 | cond2 | cluster | mcnemar_stat | mcnemar_pval |
|---|---|---|---|---|
| WT | T1 | Mixed | 526 | 6.40E-15 |

| | | | | |
|---|---|---|---|---|
| WT | T1 | 0 | 127 | 0.0093 |
| WT | T1 | 1 | 56 | 0.018 |
| WT | T1 | 2 | 195 | 0.26 |
| WT | T1 | 3 | 23 | 0.092 |
| WT | T1 | 4 | 301 | 0.57 |
| WT | T1 | 5 | 80 | 1.50E-06 |
| WT | T1 | 6 | 260 | 7.90E-21 |
| WT | T1 | 7 | 168 | 1.90E-14 |
| WT | T1 | 8 | 25 | 0.1 |
| WT | C1 | Mixed | 595 | 9.20E-09 |
| WT | C1 | 0 | 127 | 0.0021 |
| WT | C1 | 1 | 74 | 0.43 |
| WT | C1 | 2 | 195 | 0.00031 |
| WT | C1 | 3 | 37 | 0.061 |
| WT | C1 | 4 | 301 | 0.12 |
| WT | C1 | 5 | 130 | 0.17 |
| WT | C1 | 6 | 520 | 1 |
| WT | C1 | 7 | 289 | 0.046 |
| WT | C1 | 8 | 39 | 0.15 |

**Table S13**: Table of differentially expressed genes in T1 compared to 10A (lfc>= 0.1, svalue<0.01)

| Gene name | chrom | start | end | log2FoldChange | svalue |
|---|---|---|---|---|---|
| LRATD2 | chr8 | 126552443 | 126558478 | 1.53 | 4.32E-160 |
| CASC19 | chr8 | 127072694 | 127227541 | -0.36 | 1.16E-06 |
| CASC8 | chr8 | 127289808 | 127482139 | 1.16 | 7.19E-24 |
| ENSG00000286010 | chr8 | 127663280 | 127670990 | -1.32 | 0.00E+00 |
| PVT1 | chr8 | 127794526 | 128187101 | 0.28 | 2.95E-04 |
| UNC5B | chr10 | 71212570 | 71302864 | 0.62 | 2.66E-09 |
| SLC29A3 | chr10 | 71319259 | 71381423 | 0.65 | 1.02E-05 |
| VSIR | chr10 | 71747556 | 71773520 | 0.90 | 1.33E-26 |
| PSAP | chr10 | 71816298 | 71851251 | 0.59 | 7.25E-12 |
| CHST3 | chr10 | 71964395 | 72013558 | 0.69 | 9.67E-15 |
| SPOCK2 | chr10 | 72059034 | 72089032 | 1.53 | 7.87E-06 |
| ASCC1 | chr10 | 72096032 | 72217134 | 1.15 | 5.94E-71 |
| ANAPC16 | chr10 | 72216000 | 72235860 | 1.09 | 1.19E-36 |
| ENSG00000289506 | chr10 | 72272288 | 72273704 | 2.36 | 6.49E-04 |
| DDIT4 | chr10 | 72273919 | 72276036 | 1.45 | 7.05E-127 |
| DNAJB12 | chr10 | 72332830 | 72355149 | 0.71 | 3.01E-36 |
| MICU1 | chr10 | 72367340 | 72626131 | 1.03 | 3.08E-59 |

| | | | | | |
|---|---|---|---|---|---|
| MCU | chr10 | 72692143 | 72887694 | 1.36 | 2.49E-133 |
| P4HA1 | chr10 | 73007217 | 73096974 | 0.36 | 1.27E-05 |
| NUDT13 | chr10 | 73110375 | 73131828 | 1.96 | 7.12E-30 |
| ENSG00000272599 | chr10 | 73124573 | 73125532 | 1.45 | 4.49E-12 |
| ECD | chr10 | 73130155 | 73169055 | 1.30 | 3.06E-105 |
| FAM149B1 | chr10 | 73168119 | 73244504 | 0.66 | 3.85E-21 |
| ENSG00000288559 | chr10 | 73247342 | 73248268 | 0.87 | 1.71E-05 |
| MRPS16 | chr10 | 73248843 | 73252693 | 0.42 | 2.03E-17 |
| DNAJC9-AS1 | chr10 | 73252791 | 73254349 | 1.40 | 3.09E-11 |

**Table S14**: Table of differentially expressed genes in C1 compared to 10A (lfc>= 0.1, svalue<0.01)

| Gene name | chrom | start | end | log2FoldChange | svalue |
|---|---|---|---|---|---|
| LRATD2 | chr8 | 126552443 | 126558478 | 2.09 | 1.62E-304 |
| PCAT1 | chr8 | 126556323 | 127419050 | 2.11 | 1.13E-17 |
| CASC19 | chr8 | 127072694 | 127227541 | 0.86 | 1.38E-36 |
| ENSG00000224722 | chr8 | 127086263 | 127087510 | 2.13 | 1.26E-05 |
| ENSG00000287781 | chr8 | 127253213 | 127257630 | 3.13 | 7.17E-08 |
| CASC8 | chr8 | 127289808 | 127482139 | 2.90 | 1.54E-150 |
| POU5F1B | chr8 | 127322183 | 127420066 | 2.95 | 4.37E-53 |

| | | | | | |
|---|---|---|---|---|---|
| ENSG00000286010 | chr8 | 127663280 | 127670990 | -0.41 | 1.98E-04 |
| CASC11 | chr8 | 127686343 | 127738987 | 1.92 | 1.45E-05 |
| MYC | chr8 | 127735434 | 127742951 | 0.25 | 1.86E-04 |
| PVT1 | chr8 | 127794526 | 128187101 | 1.67 | 1.57E-122 |
| UNC5B | chr10 | 71212570 | 71302864 | 3.43 | 1.44E-266 |
| UNC5B-AS1 | chr10 | 71217220 | 71218294 | 1.33 | 2.33E-04 |
| SLC29A3 | chr10 | 71319259 | 71381423 | 0.74 | 6.98E-07 |
| CDH23 | chr10 | 71396920 | 71815947 | 1.77 | 1.40E-04 |
| VSIR | chr10 | 71747556 | 71773520 | 3.25 | 0.00E+00 |
| PSAP | chr10 | 71816298 | 71851251 | 3.09 | 0.00E+00 |
| ENSG00000289592 | chr10 | 71888499 | 71889171 | 3.21 | 1.39E-10 |
| CHST3 | chr10 | 71964395 | 72013558 | 1.03 | 2.71E-32 |
| SPOCK2 | chr10 | 72059034 | 72089032 | 4.04 | 5.13E-29 |
| ASCC1 | chr10 | 72096032 | 72217134 | 2.03 | 8.81E-228 |
| ANAPC16 | chr10 | 72216000 | 72235860 | 1.22 | 4.50E-46 |
| ENSG00000289506 | chr10 | 72272288 | 72273704 | 6.13 | 8.21E-14 |
| DDIT4 | chr10 | 72273919 | 72276036 | 2.14 | 6.95E-284 |
| DNAJB12 | chr10 | 72332830 | 72355149 | 0.77 | 5.06E-43 |
| MICU1 | chr10 | 72367340 | 72626131 | 1.09 | 2.50E-64 |

| | | | | | |
|---|---|---|---|---|---|
| MCU | chr10 | 72692143 | 72887694 | 2.08 | 0.00E+00 |
| P4HA1 | chr10 | 73007217 | 73096974 | 2.29 | 6.14E-203 |
| NUDT13 | chr10 | 73110375 | 73131828 | 2.63 | 1.71E-51 |
| ECD | chr10 | 73130155 | 73169055 | 2.22 | 0.00E+00 |
| FAM149B1 | chr10 | 73168119 | 73244504 | 1.78 | 8.33E-158 |
| DNAJC9 | chr10 | 73183362 | 73247255 | 0.46 | 2.33E-12 |
| ENSG00000288559 | chr10 | 73247342 | 73248268 | 1.94 | 6.42E-19 |
| MRPS16 | chr10 | 73248843 | 73252693 | 0.50 | 2.83E-25 |
| DNAJC9-AS1 | chr10 | 73252791 | 73254349 | 1.87 | 5.05E-18 |

**Table S15**: List of manually annotated translocations for 10A.

| chrom1 | start1 | end1 | chrom2 | start2 | end2 |
|---|---|---|---|---|---|
| chr3 | 894000 | 68025000 | chr9 | 23591000 | 138394717 |
| chr3 | 1235000 | 68434000 | chr5 | 118992000 | 181538259 |
| chr3 | 68025000 | 198295559 | chr9 | 0 | 20878000 |
| chr5 | 118960000 | 181538259 | chr9 | 23908000 | 138394717 |

**Table S16**: List of manually annotated translocations for T1.

| chrom1 | start1 | end1 | chrom2 | start2 | end2 |
|---|---|---|---|---|---|
| chr3 | 0 | 59225000 | chr17 | 16340000 | 83257441 |
| chr3 | 894000 | 68025000 | chr9 | 23591000 | 138394717 |
| chr3 | 1235000 | 68434000 | chr5 | 118992000 | 181538259 |
| chr3 | 56933000 | 90650000 | chr17 | 0 | 19300000 |
| chr3 | 68025000 | 198295559 | chr9 | 0 | 20878000 |
| chr5 | 118960000 | 181538259 | chr9 | 23908000 | 138394717 |
| chr6 | 0 | 170805979 | chr19 | 33715000 | 58617616 |
| chr8 | 126000000 | 128300000 | chr10 | 0 | 133797422 |

**Table S17**: List of manually annotated translocations for C1.

| chrom1 | start1 | end1 | chrom2 | start2 | end2 |
|---|---|---|---|---|---|
| chr2 | 0 | 92100000 | chr10 | 41600000 | 133797422 |

| | | | | | |
|---|---|---|---|---|---|
| chr3 | 0 | 59225000 | chr17 | 16340000 | 83257441 |
| chr3 | 0 | 68462000 | chr7 | 0 | 150000000 |
| chr3 | 894000 | 68025000 | chr9 | 23591000 | 138394717 |
| chr3 | 1235000 | 68434000 | chr5 | 118992000 | 181538259 |
| chr3 | 56933000 | 90650000 | chr17 | 0 | 19300000 |
| chr3 | 68025000 | 198295559 | chr9 | 0 | 20878000 |
| chr5 | 118960000 | 181538259 | chr9 | 23908000 | 138394717 |
| chr6 | 0 | 170805979 | chr19 | 33715000 | 58617616 |
| chr7 | 63700000 | 151700000 | chr9 | 24123000 | 35780000 |
| chr8 | 126000000 | 128300000 | chr10 | 0 | 133797422 |
| chr10 | 0 | 74500000 | chr17 | 29970000 | 83257441 |

**Table S18**: Number of nuclei and probes detected from FISH microscopy data. (A=*MYC* probe; D=enhancer probe; E=Distal non-enhancer probe.)

| | **10A (AD)** | **10A (AE)** | **C1 (AD)** | **C1 (AE)** |
|---|---|---|---|---|
| **num_valid_nuclei** | 34 | 28 | 95 | 43 |
| **num_nuclei** | 45 | 37 | 129 | 70 |
| **num_red_probes** | 62 | 62 | 269 | 140 |
| **num_green_probes** | 61 | 60 | 272 | 86 |
